# Hierarchical Approaches for Integrating Sparse, Multivariate Toxicological Effect Data in Whole Organism Molecular Dynamics

**DOI:** 10.1101/2025.05.09.652942

**Authors:** Florian Schunck, Wibke Busch, Andreas Focks

**Affiliations:** Osnabrück University, Barbarastr. 12, 49076 Osnabrück, Germany; Helmholtz-Centre for Environmental Research GmbH—UFZ, Permoserstr. 15, 04318 Leipzig, Germany

## Abstract

**Motivation:** Integrating time-resolved toxicity data observed along multiple steps along the whole-organism exposure-to-effect pathway into mechanistic ordinary differential equation (ODE) models promises improved chemical risk assessment and predictive toxicology. However, effective incorporation of such multivariate observations is mainly hindered by the inherent sparsity and variability of observation matrices characterized by low overlap in observation types between different experiments, due to tissue consumption and cost of whole-organism experiments.

**Results:** We analyzed the potential of bayesian hierarchical ODE models to estimate model parameters from sparse, non-overlapping, multivariate, time resolved datasets subject to experimental variability. The dataset in this study consisted of time series of chemical residue, gene-expression and survival obtained from 5 years of chemical exposure experiments with zebrafish embryos. Using the hierarchical approach, it was possible to increase the number of observations used in the model because experimental deviations in the exposure concentration could be accounted for. We identified inadequate dynamics of the uptake and metabolization kinetics in the ODE system, which led to strong overfitting when coupled to the hierarchical approach when wide prior distributions for the experimental error were used. In contrast, narrow distribution for the experimental error successfully avoided overfitting, despite model inadequacy. An additional simulation study of an adequate model confirmed that hierarchical models estimate parameters with low (*<* 10%) bias even under complete sparsity (i.e. no experimental overlap between measured variables); filling even small amounts (2.5%) of the missing information further reduced the parameter bias to (*<* 5%).

**Availability and Implementation:** The ODE model used in this study was implemented in the pymob model-building framework (https://github.com/flo-schu/pymob/), which provides the infrastructure for specifying high-dimensional ODE models and provides tools for bayesian inference. The case-study is available under https://github.com/flo-schu/hierarchical_molecular_tktd. The snakemake workflow for fitting pymob models is available under https://github.com/flo-schu/pymob-workflow.

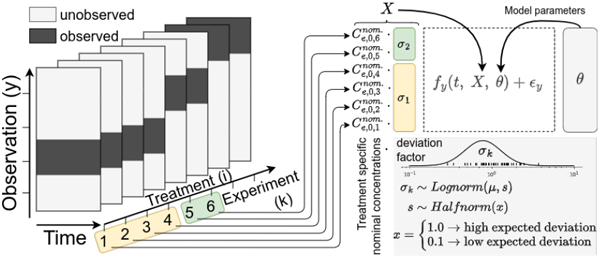

## 1 Introduction

Large amounts of diverse toxicological data from experiments in recent decades exist and are partly available in public repositories [1, 2, 3, 4]. In addition, molecular toxicity measurements from high-throughput methods are increasingly available and provide a vast data source [5, 6]. The adverse outcome pathway (AOP) framework aims to organize the different levels of biological observations into comprehensive pathways that lead to adverse effects [7, 8]. Here, integrating molecular data into mechanistic models can advance chemical risk assessment by increasing the level of biological understanding [9] and work toward predictive toxicology. However, there is still a lack of understanding and methods for integrating qualitatively and quantitatively different data sets from different experiments into mechanistic models [10], affecting not only toxicology, but also other research fields in computational biology like single-cell modeling [11]. We identified two main challenges for integrating heterogeneous data types from multiple experiments into mechanistic models:

### 1. Data sparsity

Although many datasets exist, they are not exhaustive. Experimental data have a high-dimensional data structure (experiment, treatment, measurement variable). However, due to experimental limitations, only sparse information exists in the ideal combinatorial observation matrix (Fig. 1).

**Figure 1.**
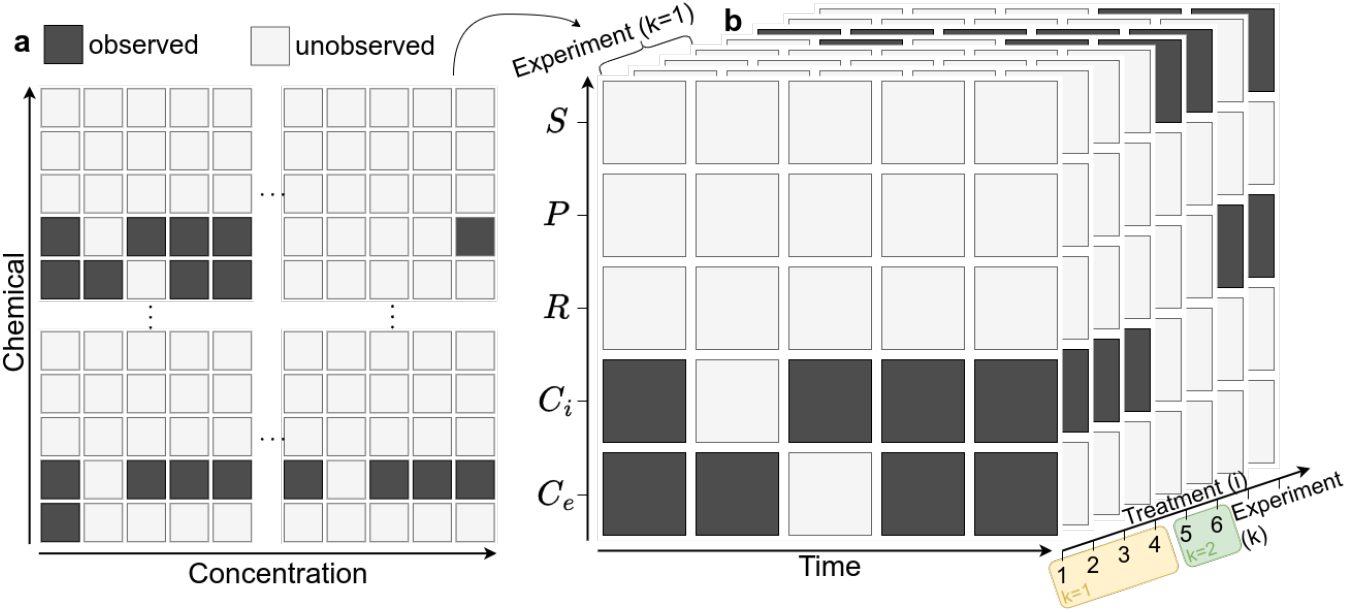
Sparse observation matrix of endpoints (rows) by time (columns). **a**: Treatments within one experiment (e.g. uptake kinetics) without overlap between concentration (Ci, Ce) and RNA (R) and survival (S). **b**: Ideal Information matrix with observed (dark boxes) and unobserved (light boxes) entries, usually a sparse matrix.

### 2. Data variabiltiy

Data sets from multiple sources and times have random (residual) errors and structured errors that are subject to experimenter differences or laboratory conditions (technical variability) or batch organism effects (biological variability) (Fig. 2).

**Figure 2.**
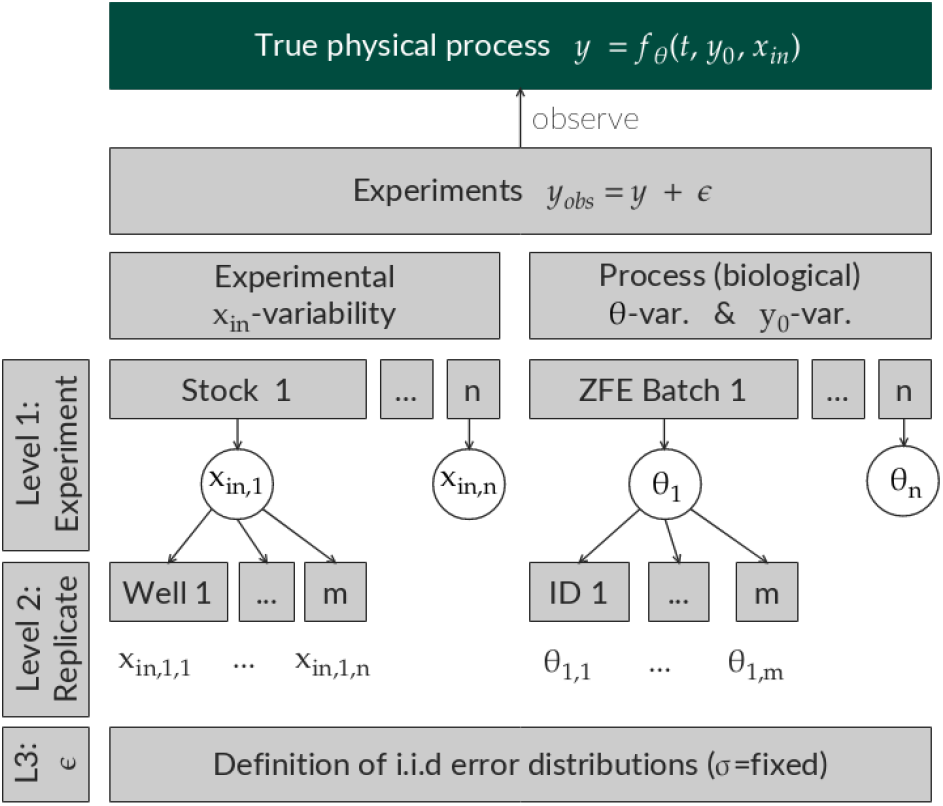
Sources of variation in experimental observations, related to the input variables in the underlying true physical process. *x*_*in*_: Input uncertainty (e.g. the exposure concentration in lab experiments, precipitation measurements in a hydrological model). *y*_0_: Starting value variability (e.g. the initial body weight, the initial RNA concentration, initial medium volume variability due to pipetting errors). *θ*: Parametric variability (e.g. the uptake rate constant variability between organism batches)

In dynamic modeling, most often ODE models are fitted to a single experimental dataset, which, if multiple variables are recorded, is a information matrix with overlapping endpoints in a single experiment. The situation of having scattered, multivariate data from different individuals, experiments or laboratories while attempting to make comprehensive assessments of the entire wealth of information is common not only in environmental toxicology but also in the medical sciences [12], marine ecology [13], machine learning [14] or social sciences [15].

Due to resource (time, money) constraints and lack of knowledge, measurements of biological systems are typically non-exhaustive, meaning that usually only few of many characteristic endpoints indicative for certain parts of biological processes are observed in a single experiment (Fig. 1-a). This is also the case in ecotoxicology where sampling is often destructive, meaning that repeated measurements over time are either not possible or require an enormous amount of samples. The few examples that record overlapping time series in the zebrafish literature include parallel measurement of internal concentrations and oxidative stress markers [16], or time resolved measurements of paraquat internal concentrations and effects on locomotion [17]. In most cases of ZFE studies, however, only a single endpoint is analyzed: internal concentrations [18, 19, 20] or survival [21, 22, 23]. This situation is not much different in other fields. If information is arranged from multiple experiments such as [24] used in [9], which compiles time-resolved toxicology measurements from zebrafish exposure studies of multiple observations, this results in sparse matrices with little overlap between observations across variables (e.g. tissue concentration, RNA, survival) and time between different experiments (Fig. 1-b). This data sparsity makes it *impossible* to separately calibrate dynamic multivariate models on single experiments. The simplest solution to this problem is to calibrate a model with a single set of parameters on the entire, sparse dataset. In this approach, models treat observational deviations as residual white noise, which can lead to biased parameter estimates if single data points or entire time series are extreme outliers [25].

The challenge of using the complete pooling approach is that parameters and boundary conditions describing the pathways may vary between organisms, treatments, and experiments due to biological variation and experimental handling errors (Fig. 2). This could lead to biased parameter estimates if, for instance, initial conditions diverge from nominal conditions and vary between individual timeseries but are not estimated separately [26]. The compromise between fitting all experiments separately and pooling all experiments in one estimate poses hierarchical modeling (also known as mixed-effects modeling or multilevel modeling), where experimental and biological variability is accounted for by clustering parameters according to the data structure of experiments and replicates. Consideration of inter-replica biological variability [27] and inter-experimental variability is crucial as it was, for example, named as a major constraint for identifying commonly expressed genes as markers for the stress response after chemical exposure in a meta study [2].

In linear regression, the concept of hierarchical models is well known and accepted as mixed-effects or multilevel models. Hierarchical models cluster the variability structure dependent on the data structure, and learn the hyper-parameter that drives e.g. the experimental variability [28]. In nonlinear dynamics, examples exist that explore the coupling of hierarchical modeling with ODE in biotechnology [26], gene-regulation models [29, 30], pharmacokinetic-pharmacodynamic (PBPK) models [31, 32], systems-biology (usually under the term non-linear mixed-effects, NLME) [33, 34], single-cell modeling [35, 36], HIV models [37, 38], and more generally [39]. The existing literature of hierarchical models, however, focuses mainly on the aspect of data variability [33]. Here, we turn to the aspect of data sparsity and probe the capabilities of hierarchical models to work with sparse data structures, which have so far not been studied for combinations of hierarchical models and ODE models to our knowledge.

Here, we study this with the example of a sparse data set without overlap between the observed variables [24] of chemical uptake and response in transcription and lethality in zebrafish embryos (ZFE). This dataset has been recently analyzed with a multivariate ODE approach [40], fitting an ODE model with a single parameter set on a reduced set of 35 experiments, due to inconsistencies between experimental conditions and observed endpoints.

We expand this work and apply a hierarchical approach to the complete dataset by estimating the *true* exposure concentration as a function of the nominal exposure concentration multiplied by an experiment-specific deviation factor *σ*_*k*_ (Fig. 1-c). This approach can be considered an extension to using a parametric distribution to estimate the initial conditions the quantity suspected to follow an extrinsic variability [41]. The approach described in this work additionally estimates the parameters of the distribution itself (hierarchically) [36]. We investigate the behavior of hierarchical models in sparse, non-overlapping data contexts and test the capacity of hierarchical models to deal with inconsistencies between nominal experimental conditions and observed experimental outcomes. We identify a erroneous behavior of the hierarchical approach when estimating the parameters of an inadequate model and cure the behavior by applying informed (narrow) priors for the experimental deviation from nominal concentrations (Fig. 1c). To study the capacity of hierarchical models to estimate parameters from sparse experimental matrices when the model description is adequate, we conduct a simulation study by systematically filling knowledge gaps in the ideal experimental matrix and calculating the resulting parameter bias. Finally, we investigate the influence of hierarchical models on loss and gradient functions for applications in gradient based parameter estimation algorithms like NUTS [42] and SVI [43].

## 2 Methods

### 2.1 Molecular TKTD Model

The differential equation model used in this work (Fig. 3c) has been described in depth [40] and is also available in the supporting information S2. In brief: The molecular TKTD model describes the multivariate response of ZFE to chemical exposure in terms of internal concentrations, regulation of the gene *nfe2l2b* coding for the transcription factor *Nrf2b* (henceforth *nrf2*) and lethality. The model was calibrated on a longitudinal (time-resolved) dataset of constant exposure to the photosynthesis inhibitor diuron, and the COX inhibitors diclofenac and naproxen between 24–120 hours post fertilization [24].

**Figure 3.**
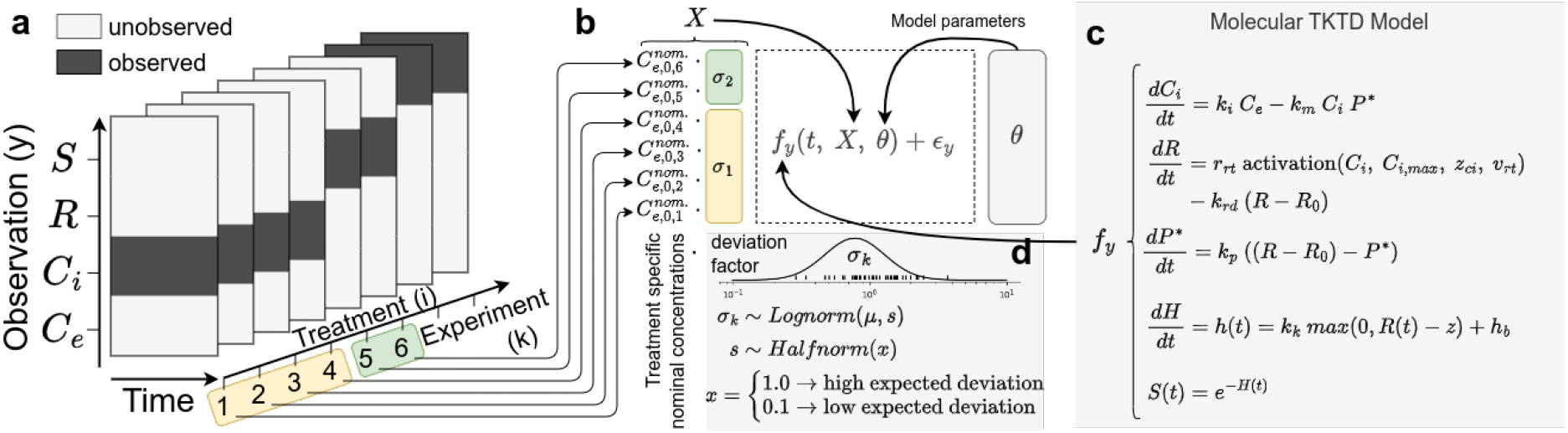
**a**: Exemplary sparse experimental observation matrix (simplified visualization compared to Fig. 1b). **b**: Quantifying technical variability between experiments by mapping treatments (*i*) and experiment-wise deviation factors (yellow and green boxes, *σ*_*k*_) to the model input *X* (Eq. 1) and relating the input *X*, parameters *θ* and time *t* to the noisy observations from the observation matrix with mechanistic model equations. **c**: Mechanistic equations from the molecular TKTD model described in [9]. **d**: Assumptions of different variability between experiments as priors for the estimation process.

### 2.2 Hierarchical Model

In this work, the method is extended to include hierarchical priors on the external concentration.

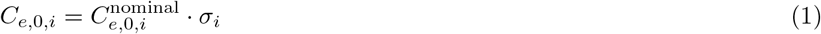

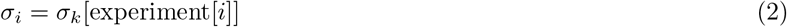

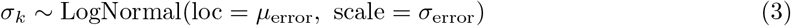

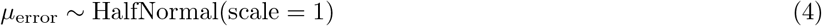

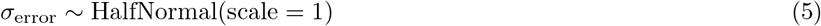

where “experiment” maps an experiment *k* (*n*_*k*_ = 42) to each id *i* (*n*_*i*_ = 314), assuming that the inter-experiment variation completely accounts for experimental variation in internal concentration. This assumption is plausible for the used dataset as the influence on replicate level variation was averaged already during the experimental process by pooling the replicates before observations are made. In addition, treatment-specific variation in the external concentration is also unlikely present, because the exposure solutions were only prepared using radial dilutions (as opposed to serial dilutions, where lower concentrations would be subject to a higher number of multiplicative pipetting errors). The remaining hierarchical error structure remains in the inter-experimental variation, where stock solutions are prepared freshly for each experiment, and here we can expect systematic deviations from the nominal concentrations for all treatments and replicates.

The models have been constructed and fitted in the open source model building platform pymob https://pymob.readthedocs.io/, which is capable of handling variably dimensional data, hierarchical variability and error specifications, gradient-based Bayesian inference for ODEs, and a suite of model development support tools.

### 2.3 Error Models

#### Survival error model

The model uses an updated likelihood formula for survival data. The conditional survival probability Pr(*t < T*| *t*_0_ *< T*) = *S*(*t*)*/S*(*t*_0_) in the interval [*t*_0_, *t*] is no longer computed in the solution of the differential equation but instead it is derived from the cumulative hazard state, by substituting *S*(*t*) = exp(*−H*(*t*)).

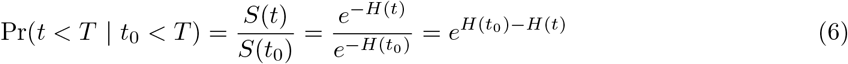

This equation is numerically more stable, since it does not involve the computation of potentially very large exponential functions, which leads to underflow issues, when the (unconditional) survival in the numerator or denominator approach the survival probability approaches the machine precision. This scenario can happen frequently in parameter estimation, and especially parameter sampling scenarios, when combinations of parameters induce the survival function to approach zero very fast.

#### Internal concentration, nrf2 error model

Qualitatively different datasets have qualitatively different error distributions. The error distributions of different observation types (e.g. Binomial survival data, continuous growth data, gene count data) may operate on different scales. Different error scales have consequences for calculating the probability of a datapoint given a distribution, because probability density values decrease with increasing scale at constant proportional deviation parameters. This is due to the property of probability distributions to have a unit integral. This means automatically, higher concentration values will be less likely than lower concentration values (even relative to the mean value). Which results in higher concentration values having a proportionally higher influence on the joint likelihood. Especially when using the deviation from predictions of ODE solutions as a metric for computing the likelihood this can lead to problems, because within one data-type and one timeseries, different observations will have different weights, due to scale differences because of the system dynamics over time.

For this reason, the models were calibrated on standardized residuals for concentrations and *nrf2* induction, assuming a multiplicative error models, using 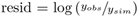. This ensures that the likelihoods of the corresponding data variables are scale invariant.

#### Weighing likelihood functions of different data types

In the used datasets, large differences in the number of observations between survival, internal concentration and *nrf2* induction data are present. The problem of weighting multivariate datasets has been discussed before [11]. We introduce a generic likelihood scaling factor *ω* to correct for unequal number of observations for *K* observed data types *k* with *n*_*k*_ observations, while keeping the general relationship between prior and likelihood function intact where.

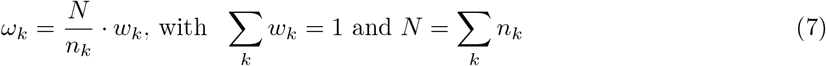

Weights *w*_*k*_ default to *w*_*k*_ = 1*/K* making all observation types equally important regardless of the number of observations. Weights can be manually adjusted, to account for differences in importance of the data types. *w*_*k*_ = *n*_*k*_*/N* to resort to an unscaled likelihood function

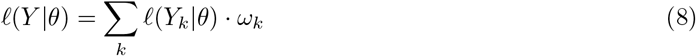

### 2.4 Parameter estimation

We use stochastic variational inference (SVI) to estimate the model parameters [44]. Variational inference approximates the posterior density of the parameters by optimizing a family of distributions to find that distribution that minimizes the Kullback-Leibner divergence [43]. SVI optimizes over random mini-batches of the data and updates the global parameters w.r.t in each iteration. SVI scales much better to large datasets than conventional MCMC sampling approaches, because the likelihood computation of mini batches is strongly reduced. This increased efficiency comes at the cost of reduced accuracy in the estimation of parameter covariance [43]. Here, we use the evidence lower bound (ELBO) to minimize KL-divergence. In particular we use a family of multivariate gaussians as target densities, to also approximate the correlation structure of the parameters. This approach is computationally somewhat more expensive than mean-field variational inference due to high-dimensional variance-covariance matrices, but in the case of ODEs with a limited amount of parameters it is a feasible and strongly suggested approach.

**Table 1.** Model comparison by Bayesian information criterion (BIC). Only RNA-Pulse-5 models with a substance-independent RNA protein module were compared. Additional models are available in the supporting information (Tab. S1)

| model | dataset | subtype | model type | priors | version | n | BIC |
| --- | --- | --- | --- | --- | --- | --- | --- |
| NU-I5-R | reduced | substance independent | non-hierarchical | uninformed priors | RNA Pulse 5 | 18 | 1761 |
| HU-I5-R | reduced | substance independent | hierarchical | uninformed priors | RNA Pulse 5 | 51 | 1826 |
| HI-I5-R | reduced | substance independent | hierarchical | informed priors | RNA Pulse 5 | 51 | 1812 |
| NU-I5-F | full | substance independent | non-hierarchical | uninformed priors | RNA Pulse 5 | 18 | 3363 |
| HU-I5-F | full | substance independent | hierarchical | uninformed priors | RNA Pulse 5 | 62 | 2754 |
| HI-I5-F | full | substance independent | hierarchical | informed priors | RNA Pulse 5 | 62 | 2984 |

### 2.5 Model building platform: pymob

Pymob was used as a platform for building the model and estimating parameters. Pymob is capable of handling variably dimensional data, hierarchical variability and error specifications, gradient based bayesian inference for ODEs with Markov Chain Monte Carlo (MCMC) and SVI, model development support tools, visualization tools and many more

## 3 Results

By modeling the external concentration as a function of the nominal external concentration multiplied by an experiment specific deviation factor *σ*_*k*_, the 42 experiments included in the entire ZFE dataset (for 24 hpf) [24] could be modeled, while previously [40], 9 experiments (IDs: 15, 16, 18, 37, 38, 39, 42, 45, 46) that clearly deviated from the nominal concentrations had to be excluded and two experiments (IDs: 2, 31) were previously not available. Using a stochastic variational inference algorithm, fits for over 1,400 data points were obtained in 5 minutes runtime on a local machine.

In total 12 models were compared (Tab. S1). For the evaluation of the hierarchical model performance, we only consider the performance of the models with the substance-independent RNA protein module (Tab. 1), as this module bears most relevance for environmental risk assessment and prevents parameter identifiability issues, due to parameter sharing of 7 out of 10 ODE parameters.

The resulting estimates allow the comparison between nominal external concentrations, estimated (posterior) external concentrations and observed external concentrations if available (Fig. 4). Experiments with the highest deviation between nominal and estimated concentrations coincide with the previously excluded experiments; especially experiment 16 (diuron exposure, *C*_*i*_ endpoint), which was previously excluded due to an evident discrepancy between the observed internal concentrations and the much lower reported nominal concentration, can now be included with an approximately 20-fold higher estimated external concentration. Also experiments 37–39 (diclofenac exposure, survival endpoint) were previously excluded, because they showed disproportionally low survival at higher concentrations than observed in other experiments. Under a hierarchical model, the external concentrations are estimated approximately an order of magnitude lower. Similarly this can be observed in experiments 45 and 46. Experiments 2 and 31 were previously not available and were under no doubts of quality issues and here we find exceptionally low deviation from the nominal concentrations, or even the observed concentrations. Finally, the bayesian information criterion (BIC) (HI-I5-F = 2984, HU-I5-F = 2754) of the hierarchical models is much lower than the non-hierarchical model’s BIC (NU-I5-F = 3363) when fitting to the full dataset, due to the reasons explained above. This is not the case if the reduced dataset is considered.

**Figure 4.**
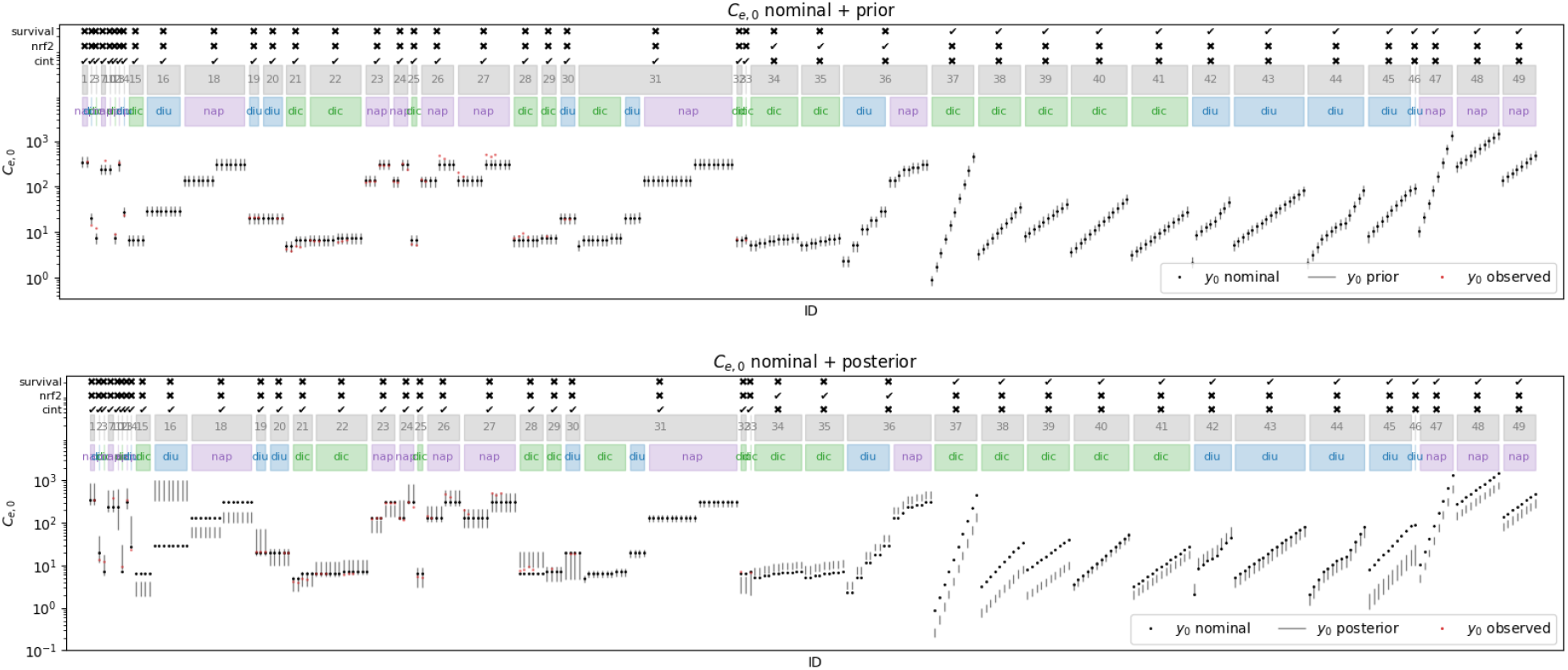
Nominal and posterior (modelled) external concentrations of the substance independent model fitted on the full dataset, using informed priors for the external concentrations (HI-I5-F). The grey boxes indicate the experiment ID and the colored boxes below indicate the used substance (blue: diuron, green: diclofenac, purple: naproxen).

The *informed* prior model (HI-I5-F) shows no systematic deviation in the modeled external concentrations *µ*_*error*_ = 1.0 *±* 0.1 from the nominal concentrations with an inter experimental variation of 0.6 log standard deviations. In contrast, the *uninformed* prior model (HI-I5-F) shows strong systematic deviation in the modeled external concentrations *µ*_*error*_ = 0.6 *±* 0.1 from the nominal concentrations with a large inter experimental variation of 1.1 log standard deviations. In the following the differences between informed (narrow) and uninformed (wide) prior distributions for the deviation from the nominal concentration are presented.

### 3.1 Informed vs. uninformed priors

The assumed scale (width of the distribution) of the lognormal *σ*_*k*_-prior, which estimates the deviation from the nominal external concentration, had a large effect on the estimated external concentrations *C*_*e*_ and correspondingly on the model fits. When an uninformed prior was applied *HalfNormal*(*σ*_*error*_ = 1.0) for the scale of the distribution, allowing extreme deviations from the external concentrations, even deviations of a factor of 1,000 are not considered improbable by the estimation algorithm. This led to extreme deviations of the external concentrations in model HU-I5-F (Fig. S20) and highly questionable model fits (Fig. S18). The increased degrees of freedom are reflected in the improved bayesian information criterion (*BIC* = 2754) for HU-I5-F compared to the non-hierarchical NU-I5-F model (*BIC* = 3363) and a low deviation from observed external concentrations for diuron and diclofenac (Fig. S21). This suggests a uncomfortable property of hierarchical models. The algorithm only maximizes the likelihood of the data given the model. If the model is not a sufficiently good approximation of the reality, the additional degrees of freedom can be exploited to minimize the likelihood especially in situation with scarce data availability. Such situations are present in this dataset, due to the sparsity of the dataset.

Using informed priors with a smaller scale parameter *HalfNormal*(*σ*_*error*_ = 0.1), considering large deviations from nominal concentrations as improbable but not impossible, the resulting extrapolations for diuron, diclofenac and naproxen are more reasonable (Fig. 5), compromised by an overall worse fit (*BIC* = 2984). Especially the diuron internal concentration data cannot be fitted in any model variant (Fig. 5j, diuron). This is caused by direct coupling between *C*_*i*_ *→nrf* 2 *→survival* and will be discussed in depth below. Correspondingly, the analysis of the residual deviations from internal concentrations showed an approximately 3 fold increase above the upper bound of the number of significant deviations from zero, compared to expected deviations if the residuals were normally distributed for diuron. Interestingly, diclofenac and naproxen showed an even higher number of significant deviations, which is attributable to the high density of initial observations, paired with a structural underestimation of the initial dynamics. The number of significant deviations for *nrf2* was at the upper bound of the expected deviations under normally distributed resiudals, which also reflects visual inspection of the residual plots (SI S4.8, S5.8). A number of additional model inadequacy metrics have been computed and are available in the SI for all computed model variants. They paint a similar picture accross all model variants and identify internal concentration as the most inadequate submodel.

**Figure 5.**
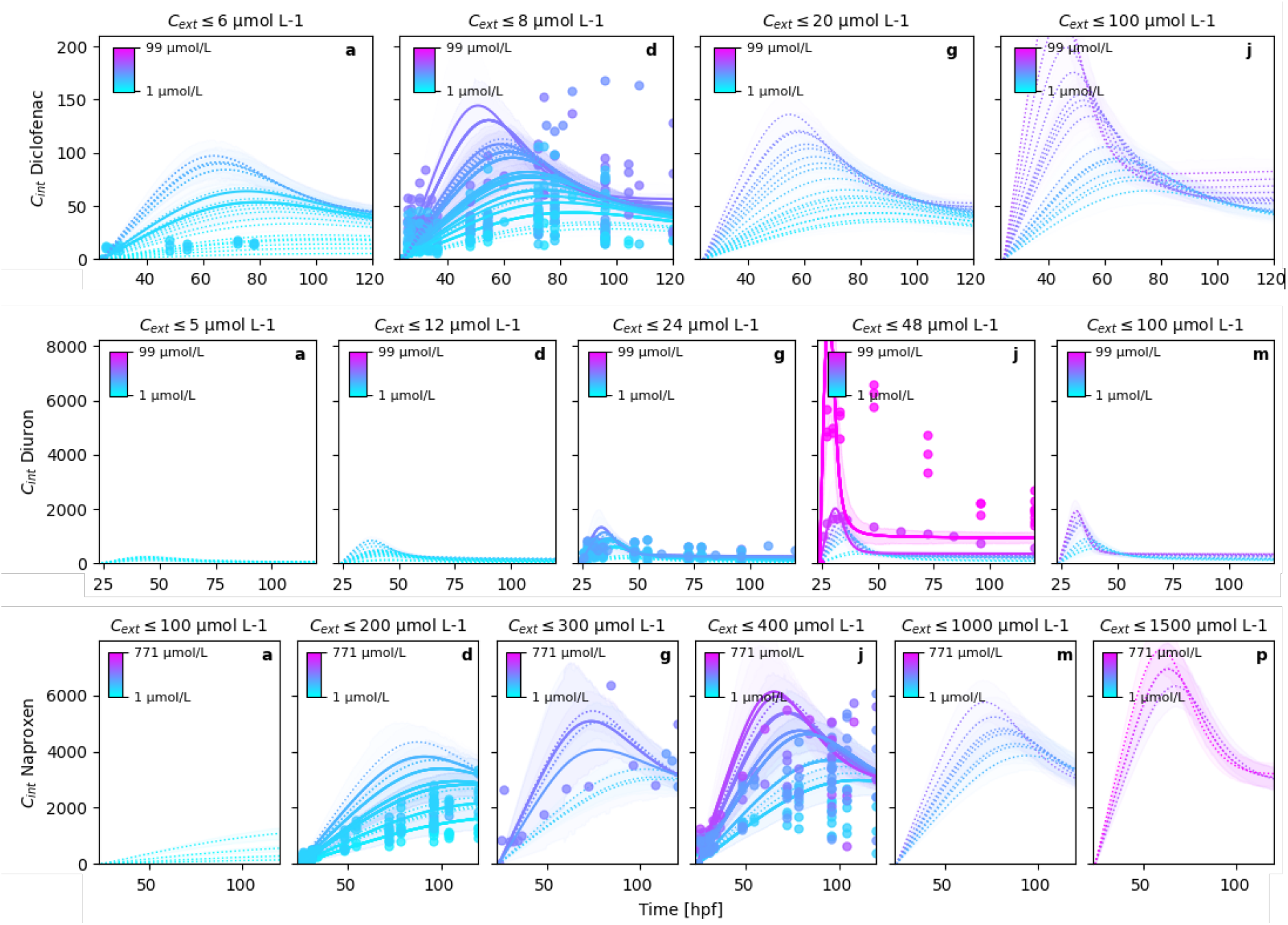
Posterior model fits and 95%-BCIs of the GUTS HI-I5-F model (hierarchical model, informed priors, substance-independent, RNA-pulse 5, full dataset) for internal concentration dynamics of diclofenac, diuron and naproxen. In order to improve the readability of the figure, columns distribute the modeled experimental data into nominal concentration classes. The solid lines are the mean posterior estimates of the endpoints over time and the dotted lines are those estimates where no data were available. The shaded areas are the 95%-BCIs indicate the posterior density intervals containing 95% probability of the posterior predictions. Note that the residual error in the observations is not included in the BCIs shown in the figures.

### 3.2 Investigating parameter estimation capabilities of hierarchical models and sparse datasets with a simulation study

To investigate whether hierarchical models can theoretically recover parameters from a sparse dataset, when the model is an adequate representation of reality, we conducted a small simulation study, where observations were simulated with the same structured technical variability as assumed above (Eq. 1). We assumed the parameter estimates from (SI S5.3), including the residual noise terms. The external concentration is fully correlated with the uptake rate coefficient. This means parameterizing a systematic offset at the same time as parameterizing the uptake coefficient leads to an identifiability issue. Therefore we have fixed the scale parameter of the lognormal distribution of experimental deviation at 1 (no systematic deviation) and only estimate the shape parameter (deviation of the experiments from nominal concentration). First we recovered the parameters from the sparse observational matrix of the dataset described in the previous sections and applied to the synthetic dataset. We then went on to randomly unmask observations to decrease the sparsity of the information matrix and increase the overlap of measured variables within experiments (Fig. 6).

**Figure 6.**
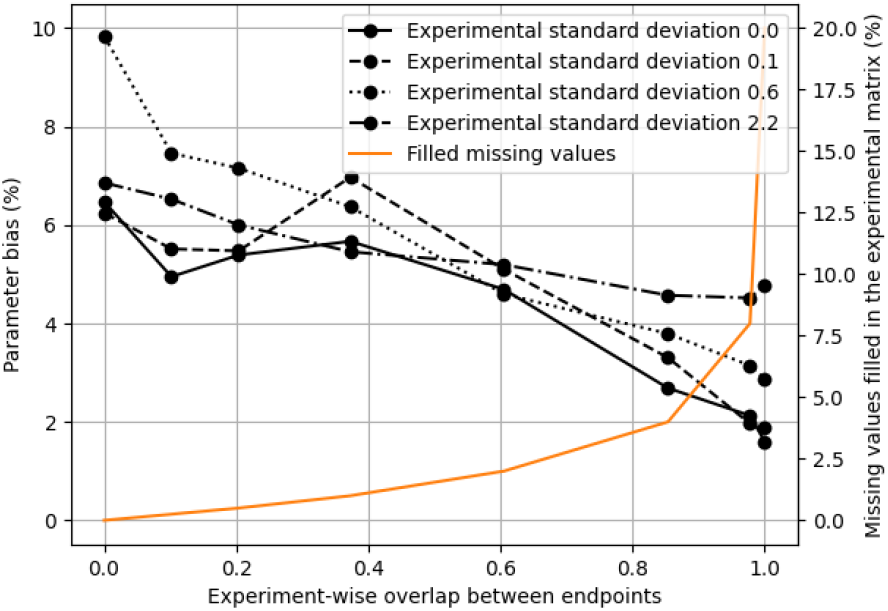
Influence of within experiment overlap of measured variables on parameter bias. The lines represent the average parameter bias from 20 model fits with random draws from lognormal distributions of experiment specific deviation factors with different standard deviations (0.0, 0.1, 0.6, 2.2) and random residual error drawn from lognormal distributions centered around the model fits with a shape parameter of 0.1. The overlap is computed as the fraction of experiments where at least two variables were observed at least once over time. The orange line is the fraction of missing observations from the ideal information matrix (id: 314 x time: 23 x variable: 3) compared to the sparse experimental matrix that has been unmasked from the synthetic dataset used in the simulation study.

The simulation study shows that the parameter bias is largely independent from the magnitude of within-experiment deviation from the nominal concentrations. Only at 2.2 standard deviations of the distribution of deviation factors, where experimental deviations of a factor of 10 can easily occur, the parameters cannot be estimated with a bias below 5%. Generally, the parameters can be extracted well from the synthetic data, indicated by a maximum bias of 10% in the sparse matrix with no variable overlap in a single experiment. The simulation study also shows that even low fractions of randomly added data (0.25%*≈* 50 data points) reduces the parameter bias and 2.5% of added data (*≈* 500 data points) reduce the parameter bias to 5%, irrespective of the magnitude of deviation. This finding indicates that recording and publication of even single measurements of additional variables within the same experiments can vastly improve parameter estimation of nonlinear models fitted on multivariate data sets.

### 3.3 Hierarchical modelling had no major effect on likelihood landscape near the global optimum

We finally went on to study the effects of hierarchical modelling approaches on likelihood landscapes and corresponding gradient vector fields. Using state-of-the-art tools like diffrax, implemented in pymob allows the automated computation of likelihood values and vector fields for parameter combinations of ODE models. We visualized the correlation structures between parameters that go beyond estimates obtained with SVI, and used it to improve the model specification and to identify problematic parameters in complex models.

The computed likelihood landscapes of the original model implementation [9] (TKTD-RNA-PULSE-4.0, Fig. 7a–c) exhibit rugged landscapes with erratic local minima and partly extreme errors in the vector field for the parameter combination *v*_*rt*_–*k*_*m*_.

**Figure 7.**
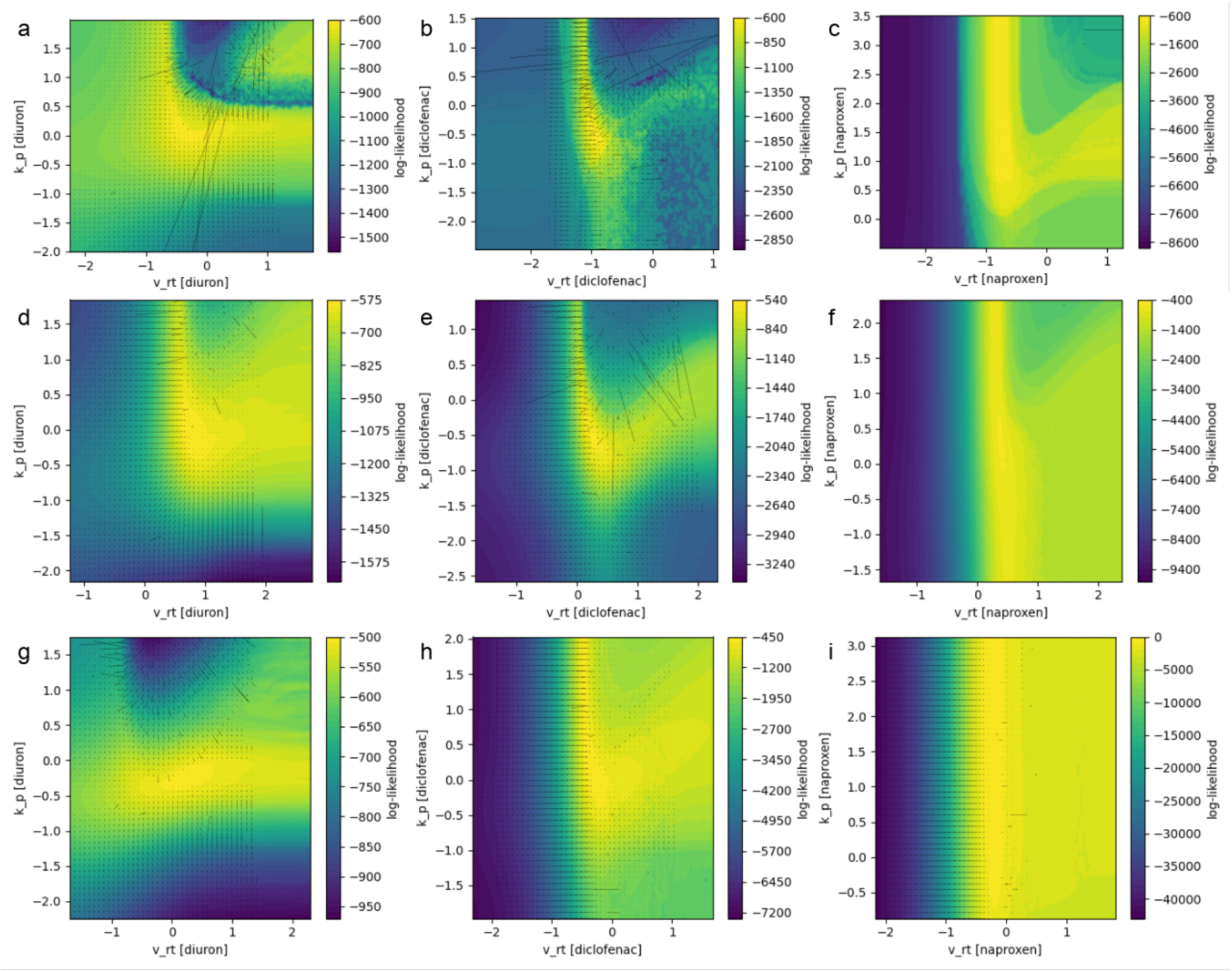
Exemplary marginal likelihood landscapes (color) and vector field of the gradients (arrows) of the parameter combinations activation velocity *v*_*rt*_ and dominant rate constant of the protein dynamic *k*_*p*_. The parameters are given on an standardized scale which is during fitting mapped to the transformed space of the prior. We centered the parameter exploration at the mode of the posterior distribution. **a–c** show the marginal parameter relationship for TKTD-RNA-PULSE-4.0; **d–f** show the marginal parameter relationship for TKTD-RNA-PULSE-5.0 (with a numerically stabilized likelihood function for conditional survival, Eq. 6), and **g–i** show the marginal parameter relationship for TKTD-RNA-PULSE-5.0 under the assumption of hierarchical errors in the external concentration (Eq. 1).

These problems led to model improvements with respect to the error model formulation (Eq. **??**) resolving most of the erroneous behavior of the likelihood function (Fig. 7d–e). In contrast, no evident improvement is visible comparing in the likelihood landscapes of the hierarchical approach (Fig. 7g–i) although the vector fields seem to become slightly more stable. It is noticeable that the gradients of the likelihood functions cannot be computed for large parameter values. Note that large in this case refers to the transformed (typically lognormal) space of the parameters. Currently we have no explanation for this behavior.

Finally, we observe that parameter sharing approaches produce the smoothest likelihood functions and the most stable vector fields, although the problems of computing gradients for very high values of the parameters persist. Figure 8.

**Figure 8.**
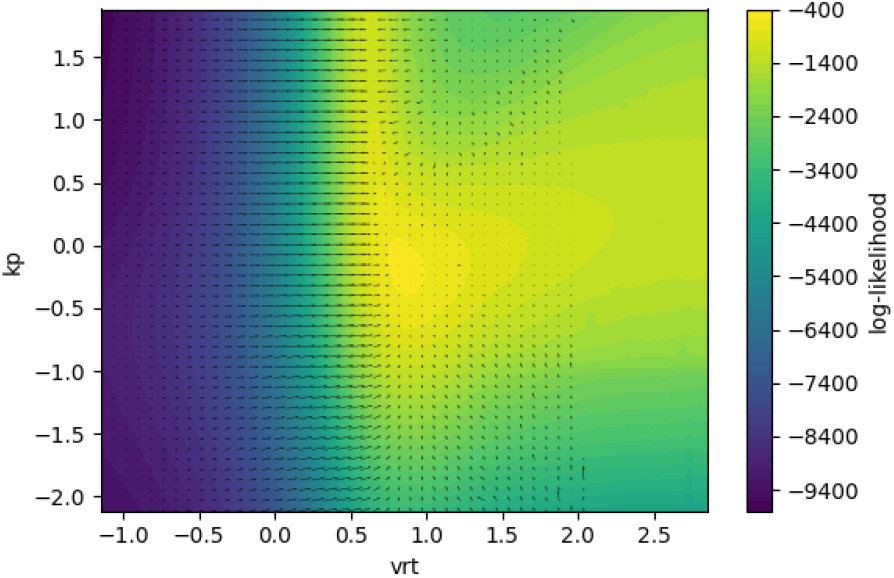
Exemplary marginal likelihood landscapes (color) and vector field of the gradients (arrows) of the parameter combinations activation velocity *v*_*rt*_ and dominant rate constant of the protein dynamic *k*_*p*_ for the TKTD-RNA-PULSE-5.0 (Eq. 6 under hierarchical assumption (Eq. 1) and parameter sharing in the RNA-protein module.

## 4 Discussion

Hierarchical modeling is well established in regression modeling [28, 45]. The coupling of dynamic models and hierarchical parameter estimation is gaining attention in computational biology [34, 11]. In general, multivariate analysis is considered at the forefront of research in dynamic modeling [46].

Here, we applied a hierarchical multivariate ODE model to an extremely sparse dataset with no overlap in data types between experiments, which has not been done before to our best knowledge. While sparsity in replicates of the same measurement type been successfully treated in NLME modelling, e.g. in cell-to-cell variability [41, 34], we are only aware of one application of NLME model to multivariate datasets [47], where Laplace approximation was necessary to ease the computational burden of the computations. It has been previously mentioned that NLME methods suffer under several limitations such as scalability of mechanistic model calibration due to high computational costs of the method, integration of multiple data types with imbalanced number of observations, availability of software tools [11]. The presented case study addresses these limitations: We provide an open-source modeling framework pymob (https://github.com/flo-schu/pymob/), capable of handling multidimensional data and coupling it to state-of-the art autodifferentiable numerical solvers [48], and inference engines [49] that even scale to massive datasets.

For the reasons mentioned above it is no surprise that meta studies of existing experimental data have so far not been conducted with mechanistic models, because the necessary tools for such an undertaking were not available. Here we have exemplified how hierarchical ODE models with informed priors on the experimental variability could rise to the challenge and be applied to re-analyze existing univariate datasets. For ecotoxicology (e.g. survival) and multimultivariate datasets, by accounting for the structured source of variation between experiments conducted in different laboratories and years. Such efforts would also increase the reliability of parameter estimates of mechanistic models, by accounting for multiple sources of variation, and hence increase the trust and acceptance of mechanistic models many research fields. Coupling of hierarchical modeling with whole-organism dynamic models could also help to separate experimental variation induced by batch effects from parameter uncertainty due to residual biological variation. This could help advance multi-species approaches [50] and multi-chemical approaches in (eco)toxicology.

### 4.1 Hierarchical models operating on sparse data structures should only be used with narrow priors on experimental variability

Using conservative, narrow priors for the error scale for technical variability (here expressed by the external concentration) in a hierarchical experiment design is highly recommended in a model that measures only one endpoint per experiment in a multi-endpoint process model, and is potentially inadequate for one or more measured endpoints. Therefore, we identify the substance-independent model fitted on the full dataset using informed priors for the external concentrations (HI-I5-F) as the overall best model in this work. While the model fit is inferior (*BIC* = 2984) to the model using uninformed priors (HU-I5-F, *BIC* = 2754), the capability of modeling the entire dataset, while not being prone to overfitting due to the additional degrees of freedom, made it the overall best model. In future work it might be interesting to investigate t-distributions, cauchy distributions or laplace distributions [25] as prior distributions for the experimental errors, because they may be better suited to detect outlying experiments than lognormal distributions, which allocate large amounts of probability mass into the tails of the distribution.

### 4.2 The inadequacy of the internal concentration submodule in the molecular TKTD model is driven by the joint role of *nrf2* in lethality and metabolization of the parent compound

While the hierarchical model worked well when informed (conservative) priors were used, we identify a risk of overfitting when the model is not a perfect description of the reality and uninformed priors are used. That inadequate ODE models can severely bias parameter estimates has been observed before in simulation studies of RNA expression [30]. In the presented case, the problem is exacerbated by the sparse data structure.

The diuron fit revealed the source of the problem. In all model variants, the internal concentration dynamic of diuron exposure could not be estimated. This is also true to a lesser degree for diclofenac and naproxen (SI S5.8, S4.8). In the RNA-Pulse-5 models, *nrf2* is expressed after internal concentrations *C*_*i*_ exceed a threshold. *nrf2* is also directly linked to lethality. The diuron case is characteristic: The sharp increase, followed by flattening of the curve in survival data, together with the pronounced peak in *nrf2* data force the model to describe the *C*_*i*_ dynamic as a very sharp peak, which is an inadequate model for internal concentrations. This is reflected by computed model inadequacy metrics that are most strongly increased for internal concentrations. While diuron modeled dynamics prominently deviate from the internal concentrations, residual dynamics figures and computed model inadequacy metrics indicate that the internal concentration dynamics are underpredicting the internal concentrations dynamics in the first 24 hours post fertilization (hpf), followed by a systematic overprediction in from 24–80 hpf for diclofenac and naproxen.

Model inadequacy is an active field of research and a variety of approaches have been proposed, such as the *statistical operator approach* for chemical kinetic modeling, which describes the residual dynamics in the ODE while respecting thermodynamic constraints [39]. While this approach is highly promising it requires immense effort to develop such descriptor. Another more generic approach was outlined in the KOH framework [51], which employes gaussian processes to describe and remedy model inadequacy, but is still plagued by parameter identifiability issues [52]. More recently physics informed neural networks have been employed to study model inadequacy [53]. We identified key limitations in the molecular TKTD model applied to the sparse, non-overlapping dataset, particularly the simplification of metabolization and lethality as driven by a single factor. Addressing this requires separate modeling of these processes, incorporating enzyme kinetics, and leveraging additional datasets. Furthermore, we propose integrating high-capacity function approximators within a universal differential equation framework [54, 48] to capture residual dynamics and improve model accuracy.

## 5 Conclusion

Hierarchical dynamic models are an emerging tool to integrate unrelated time resolved datasets into multivariate explanatory models. In this work we have probed the capacity of such models to handle complex situations of non-overlapping, noisy datasets and partial model inadequacy—a scenario that meets the research reality. While we’ve demonstrated the potential of this approach, we share a couple of recommendations for the application: Hierarchical dynamic models should always be used to pool information from multiple experiments if each experiment contains all types of data, because hierarchical methods can easily integrate outlying experiments, those that might otherwise be excluded from the analysis. This is especially helpful, when past data are analysed and not all information are available or can be procured in a reasonable effort. Under the situation we find in this work, i.e. the model is in parts an indequate representation of reality (diuron internal concentration dynamics) *and* if the data variables are only separately recorded in each experiment, hierarchical models should be used with great care. This is because the introduced experiment-wise hierarchy allows the optimization algorithm to hide unexplained dynamics in the data in the additional experiment-specific parameters. Ways to cope with such problems are: using more narrow hyperpriors, using information about the experimental variability directly as parameters for the additional hierarchy level, using a more restrictive clustering levels, or estimating the residual dynamics.

## Acknowledgements

The authors thank Michael Osthege for the helpful discussions on the specification of priors for hierarchical ODE models in Bayesian parameter inference. The study was supported by the Helmholtz Program oriented funding (POF IV) Earth and Environment Research Program, Topic 9 - Healthy Planet. Computations on the HPC of Osnabrück University were funded by the Deutsche Forschungsgemeinschaft (DFG, German Research Foundation) - 456666331.

## Supporting information

- The model description of the molecular TKTD model, complete model comparison as well as model reports referenced in the manuscript are available in the file supporting-information.pdf
- The code used to generate the results including install and usage instructions are available on https://gitlab.uni-osnabrueck.de/molecular-tktd/hierarchical_tktd
- The molecular TKTD model variants are available on https://github.com/flo-schu/tktd_rna_pulse
- The model building and parameter estimation platform pymob is available https://github.com/flo-schu/pymob and documentation is available on https://pymob.readthedocs.io/en/latest/
- The datasets used in this study is available on https://doi.org/10.12751/g-node.l6jqgf
- Additional results for the GUTS-RNA-pulse models are available on https://gin.g-node.org/flo-schu/tktd_rna_pulse results
- Additional results for the standard GUTS models (reduced, scaled-damage, full GUTS) models are available on https://gin.g-node.org/flo-schu/guts results

