## supplementary-information.pdf for "Hierarchical Approaches for Integrating Sparse, Multivariate Toxicological Effect Data in Whole Organism Molecular Dynamics"

This document contains 42 Figures and 2 Tables on 61 pages.

#### Contents

**S1 Complete model comparison** S3

**S2 Molecular TKTD model** S4

**S3 Report(case_study=tktd_rna_pulse, scenario=rna_pulse_5_substance_independent_rna_protein_moc**
- S3.1 Report: Model ✓ . . . S6
  - S3.1.1 Model . . . S6
  - S3.1.2 Solver post processing . . . S8
  - S3.1.3 Probability model . . . S8
- S3.2 Report: Parameters ✓ . . . S8
  - S3.2.1 $x_{in}$ . . . S8
  - S3.2.2 $y_0$ . . . S8
  - S3.2.3 Free parameters . . . S14
  - S3.2.4 Fixed parameters . . . S15
- S3.3 Report: Table parameter estimates ✓ . . . S15
- S3.4 Report: Goodness of fit ✓ . . . S15
- S3.5 Report: Diagnostics ✓ . . . S16
- S3.6 Report: Visualizations ✓ . . . S18
- S3.7 Report: Model inadequacy metrics ✓ . . . S20
  - S3.7.1 Residuals . . . S20
  - S3.7.2 Model inadequacy . . . S22

**S4 Report(case_study=hierarchical_molecular_tktd, scenario=hierarchical_cext_nested_sigma_hyperpr**
- S4.1 Report: Model ✓ . . . S24
  - S4.1.1 Model . . . S24
  - S4.1.2 Solver post processing . . . S26
  - S4.1.3 Probability model . . . S26
- S4.2 Report: Parameters ✓ . . . S26
  - S4.2.1 $x_{in}$ . . . S26
  - S4.2.2 $y_0$ . . . S26
  - S4.2.3 Free parameters . . . S32
  - S4.2.4 Fixed parameters . . . S33

S4.3 Report: Table parameter estimates ✓ . . . S33
S4.4 Report: Goodness of fit ✓ . . . S33
S4.5 Report: Diagnostics ✓ . . . S34
S4.6 Report: Visualizations ✓ . . . S36
S4.7 Report: Visualizations ✓ . . . S36
S4.7.1 $y_0$ estimation of external concentrations . . . S38
S4.8 Report: Model inadequacy metrics ✓ . . . S39
S4.8.1 Residuals . . . S39
S4.8.2 Model inadequacy . . . S41

**S5 Report(case_study=hierarchical_molecular_tktd, scenario=hierarchical_cext_nested_sigma_hyperpr**
S5.1 Report: Model ✓ . . . S43
S5.1.1 Model . . . S43
S5.1.2 Solver post processing . . . S45
S5.1.3 Probability model . . . S45
S5.2 Report: Parameters ✓ . . . S45
S5.2.1 $x_{in}$ . . . S45
S5.2.2 $y_0$ . . . S45
S5.2.3 Free parameters . . . S51
S5.2.4 Fixed parameters . . . S52
S5.3 Report: Table parameter estimates ✓ . . . S52
S5.4 Report: Goodness of fit ✓ . . . S52
S5.5 Report: Diagnostics ✓ . . . S53
S5.6 Report: Visualizations ✓ . . . S55
S5.7 Report: Visualizations ✓ . . . S55
S5.7.1 $y_0$ estimation of external concentrations . . . S57
S5.8 Report: Model inadequacy metrics ✓ . . . S58
S5.8.1 Residuals . . . S58
S5.8.2 Model inadequacy . . . S60

### List of Figures

Figure S1 Directed acyclic graph (DAG) of the probability model. . . . S8
Figure S2 Paired parameter estimates . . . S16
Figure S3 Psuedo trace, generated for draws from the optimized SVI distribution . . . S17
Figure S4 Posterior model fits . . . S18
Figure S5 Posterior model fits . . . S19
Figure S6 Posterior model fits . . . S19
Figure S7 Resiudal cint dynamics of diuron . . . S20
Figure S8 Resiudal cint dynamics of diclofenac . . . S20
Figure S9 Resiudal cint dynamics of naproxen . . . S21
Figure S10 Resiudal nrf2 dynamics of diuron . . . S21
Figure S11 Resiudal nrf2 dynamics of diclofenac . . . S22
Figure S12 Resiudal nrf2 dynamics of naproxen . . . S22
Figure S13 Directed acyclic graph (DAG) of the probability model. . . . S26
Figure S14 Paired parameter estimates . . . S34
Figure S15 Psuedo trace, generated for draws from the optimized SVI distribution . . . S35
Figure S16 Posterior model fits . . . S36
Figure S17 Posterior model fits . . . S37
Figure S18 Posterior model fits . . . S37
Figure S19 Prior $C_{ext,0}$ estimates and nominal concentrations . . . S38
Figure S20 Posterior $C_{ext,0}$ estimates and nominal concentrations . . . S38

Figure S21 $C_{ext,0}$ comparison between nominal, measured and estimated concentrations . . S38
Figure S22 Resiudal cint dynamics of diuron . . . . . . . . . . . . . . . . . . . . . . . . . . S39
Figure S23 Resiudal cint dynamics of diclofenac . . . . . . . . . . . . . . . . . . . . . . . . S39
Figure S24 Resiudal cint dynamics of naproxen . . . . . . . . . . . . . . . . . . . . . . . . S40
Figure S25 Resiudal nrf2 dynamics of diuron . . . . . . . . . . . . . . . . . . . . . . . . . . S40
Figure S26 Resiudal nrf2 dynamics of diclofenac . . . . . . . . . . . . . . . . . . . . . . . . S41
Figure S27 Resiudal nrf2 dynamics of naproxen . . . . . . . . . . . . . . . . . . . . . . . . S41
Figure S28 Directed acyclic graph (DAG) of the probability model. . . . . . . . . . . . . . S45
Figure S29 Paired parameter estimates . . . . . . . . . . . . . . . . . . . . . . . . . . . . . S53
Figure S30 Psuedo trace, generated for draws from the optimized SVI distribution . . . . . S54
Figure S31 Posterior model fits . . . . . . . . . . . . . . . . . . . . . . . . . . . . . . . . . S55
Figure S32 Posterior model fits . . . . . . . . . . . . . . . . . . . . . . . . . . . . . . . . . S56
Figure S33 Posterior model fits . . . . . . . . . . . . . . . . . . . . . . . . . . . . . . . . . S56
Figure S34 Prior $C_{ext,0}$ estimates and nominal concentrations . . . . . . . . . . . . . . . S57
Figure S35 Posterior $C_{ext,0}$ estimates and nominal concentrations . . . . . . . . . . . . . S57
Figure S36 $C_{ext,0}$ comparison between nominal, measured and estimated concentrations . . S57
Figure S37 Resiudal cint dynamics of diuron . . . . . . . . . . . . . . . . . . . . . . . . . . S58
Figure S38 Resiudal cint dynamics of diclofenac . . . . . . . . . . . . . . . . . . . . . . . . S58
Figure S39 Resiudal cint dynamics of naproxen . . . . . . . . . . . . . . . . . . . . . . . . S59
Figure S40 Resiudal nrf2 dynamics of diuron . . . . . . . . . . . . . . . . . . . . . . . . . . S59
Figure S41 Resiudal nrf2 dynamics of diclofenac . . . . . . . . . . . . . . . . . . . . . . . . S60
Figure S42 Resiudal nrf2 dynamics of naproxen . . . . . . . . . . . . . . . . . . . . . . . . S60

#### List of Tables

Table S1 Compared models and bayesian information criterion (BIC). . . . . . . . . . . . S3
Table S2 TKTD state variables and parameters used in the GUTS-RNA-pulse model. The column “Assumed substance independence” indicates whether a parameter is supposed to be shared for multiple substances. . . . . . . . . . . . . . . . . . . . . . . . . . . . . S5

### S1 Complete model comparison

**Table S1.** Compared models and bayesian information criterion (BIC).

| model | dataset | subtype | model type | priors | version | n | BIC |
|---|---|---|---|---|---|---|---|
| NU-S5-R | reduced | substance specific | non-hierarchical | uninformed priors | RNA Pulse 5 | 32 | 1790 |
| NU-S4-R | reduced | substance specific | non-hierarchical | uninformed priors | RNA Pulse 4 | 32 | 2019 |
| HU-S5-R | reduced | substance specific | hierarchical | uninformed priors | RNA Pulse 5 | 65 | 1835 |
| HI-S5-R | reduced | substance specific | hierarchical | informed priors | RNA Pulse 5 | 65 | 1823 |
| NU-I5-R | reduced | substance independent | non-hierarchical | uninformed priors | RNA Pulse 5 | 18 | 1761 |
| NU-I4-R | reduced | substance independent | non-hierarchical | uninformed priors | RNA Pulse 4 | 18 | 1816 |
| HU-I5-R | reduced | substance independent | hierarchical | uninformed priors | RNA Pulse 5 | 51 | 1826 |
| HI-I5-R | reduced | substance independent | hierarchical | informed priors | RNA Pulse 5 | 51 | 1812 |
| NU-S5-F | full | substance specific | non-hierarchical | uninformed priors | RNA Pulse 5 | 32 | 3234 |
| HU-S5-F | full | substance specific | hierarchical | uninformed priors | RNA Pulse 5 | 76 | 2970 |
| HI-S5-F | full | substance specific | hierarchical | informed priors | RNA Pulse 5 | 76 | 2955 |
| NU-I5-F | full | substance independent | non-hierarchical | uninformed priors | RNA Pulse 5 | 18 | 3363 |
| HU-I5-F | full | substance independent | hierarchical | uninformed priors | RNA Pulse 5 | 62 | 2754 |
| HI-I5-F | full | substance independent | hierarchical | informed priors | RNA Pulse 5 | 62 | 2984 |

#### S2 Molecular TKTD model

$$\frac{dC_i}{dt} = k_i \ C_e - k_m \ C_i \ P^* \qquad \text{(Eq. S1)}$$

$$\frac{dR}{dt} = r_{rt} \ \text{activation}(C_i, \ C_{i,max}, \ z_{ci}, \ v_{rt}) - k_{rd} \ (R - R_0) \qquad \text{(Eq. S2)}$$

$$\frac{dP^*}{dt} = k_p \ ((R - R_0) - P^*) \qquad \text{(Eq. S3)}$$

$$\frac{dH}{dt} = h(t) = k_k \ max(0, R(t) - z) + h_b \qquad \text{(Eq. S4)}$$

$$S(t) = e^{-H(t)} \qquad \text{(Eq. S5)}$$

with

$$activation(C_i, \ C_{i,max}, \ z_{ci}, \ v_{rt}) = 0.5 + \frac{1}{\pi} \ arctan(v_{rt} \ (\frac{C_i}{C_{i,max}} - z_{ci})) \qquad \text{(Eq. S6)}$$

**Table S2.** TKTD state variables and parameters used in the GUTS-RNA-pulse model. The column "Assumed substance independence" indicates whether a parameter is supposed to be shared for multiple substances.

| Symbol | Definition | Unit | Assumed substance independence |
|---|---|---|---|
| Model state variables | | | |
| $C_e$ | Environmental concentration in the aqueous medium | $\mu mol\ L^{-1}$ | |
| $C_i$ | Internal concentration of the homogenized ZFE | $\mu mol\ L^{-1}$ | |
| $R$ | Relative differential RNA transcription in the ZFE | fc [c] | |
| $R_0$ | Relative differential initial RNA transcription in the ZFE | fc [c] | |
| $P^*$ | Scaled protein concentration in the ZFE | fc [c] | |
| $h$ | Instantaneous hazard rate at time $t$ | $h^{-1}$ | |
| $H$ | Cumulative hazard at time $t$ | − | |
| $S$ | Survival probability of a ZFE at time $t$ | − | |
| Model parameters | | | |
| $k_i$ | Uptake rate constant of the chemical into the internal compartment of the ZFE | $h^{-1}$ | no |
| $k_m$ | Scaled metabolization rate constant from the internal compartment of the ZFE | $h^{-1}$ | no |
| $z_{\text{ci}}$ | Scaled internal concentration threshold for the activation of *nrf2* expression | − [d] | no |
| $v_{\text{rt}}$ | Scaled responsiveness of the *nrf2* activation (slope of the activation function) | − [d] | yes/no [a] |
| $r_{\text{rt}}$ | Constant *nrf2* expression rate after activation [b] | fc [c] | yes |
| $k_{\text{rd}}$ | Nrf2 decay rate constant | $h^{-1}$ | yes |
| $k_p$ | Dominant rate constant of synthesis and decay of metabolizing proteins | $h^{-1}$ | yes |
| $z$ | Effect *nrf2*-threshold of the hazard function [b] | fc [c] | yes |
| $k_k$ | killing rate constant for *nrf2* [b] | $\text{fc}^{-1}\ h^{-1}$ [c] | yes |
| $h_b$ | background hazard rate | $h^{-1}$ | yes |
| $\sigma_{\text{cint}}$ | Standard deviation of the lognormal distribution of the internal concentration | − | yes |
| $\sigma_{nrf2}$ | Standard deviation of the lognormal distribution of the *nrf2* expression [b] | − | yes |

[a] In an unscaled version of the activation function, $v_{rt}$ is not considered substance independent, due to an inverse relationship between $v_{rt}$ and $C_{i,max}$

[b] relative to the *nrf2* concentration in untreated ZFE (fold-change)

[c] fc: fold change $\frac{\mu mol\ nrf2\text{-treatment}\ L^{-1}}{\mu mol\ nrf2\text{-control}\ L^{-1}}$

[d] scaled internal concentrations: $\frac{\mu mol\ C_i(t)\ L^{-1}}{\mu mol\ C_{i,max}\ L^{-1}}$

#### S3 Report(case_study=tktd_rna_pulse, scenario=rna_pulse_5_substance_indep

- Using `tktd_rna_pulse==0.2.9`
- Using `pymob==0.5.6a3`
- Using backend: `NumpyroBackend`
- Using settings: `../tktd_rna_pulse/scenarios/rna_pulse_5_substance_independent_rna_protein_modul`

##### S3.1 Report: Model ✓

###### S3.1.1 Model

```python
def tktd_rna_5(t, X, r_0, k_i, r_rt, r_rd, z_ci, v_rt, k_p, k_m, h_b, kk, z, ci_max):
    """
    A simplified RNA pulse model.

    This function models gene expression and metabolization of the internal
    concentration of a substance. The gene expression is controlled by a arctan
    step function based on the internal concentration (Ci) and switches on the
    gene's expression. The gene then translates a Protein, which metabolizes
    the internal concentration proportional to its expression level. The concept
    of protein must be understood not as a single Protein but as a collection
    of detoxification measures, which reduce the internal concentration of the
    compound and keep it at a reasonable level.

    Changes w.r.t. to RNA 4 model
    -------------------------------
    - The model does not evolve the survival probability S over time any more
      This is done more efficiently in the post processing, by simply taking the
      exponent


    Parameters
    ----------
    t : float
        Timestep at which the model is evaluated.

    X : tuple
        A tuple containing three elements:
        - Ce : float
            The external concentration.
        - Ci : float
            The internal concentration.
        - R : float
            The gene expression level.
        - P : float
            The protein level.
        - H : float
            The cumulative hazard.

    r_0 : float
        Initial value of the gene expression level.

    k_i : float
```

```
        Internal consumption rate constant.

    k_m : float
        Metabolization rate constant.

    r_rt : float
        Maximum gene expression rate constant. Termed k_rt in the paper

    r_rd : float
        Gene degradation rate constant. Termed k_rd in the paper

    z_ci : float
        The threshold for gene expression.

    v_rt : float, optional
        The slope parameter for the inverse tangent step function. This
        parameter regulates the responsiveness of the gene expression induction.

    k_p : float
        Dominant protein translation rate konstant.

    Returns
    --------
    dCe_dt : float
        The rate of change of external concentration.

    dCi_dt : float
        The rate of change of internal concentration.

    dR_dt : float
        The rate of change of gene expression level.

    dR_dt : float
        The rate of change of protein level.

    dH_dt : float
        The hazard rate h(t) = b * max(D, 0) + h_b
    """
    Ce, Ci, R, P, H = X

    # active = 0.5 + (1 / jnp.pi) * jnp.arctan(v_rt * (Ci / ci_max - z_ci))
    active = 1 / (1 + jnp.exp(- v_rt * (Ci/ci_max - z_ci)))

    dCe_dt = 0.0
    dCi_dt = Ce * k_i - Ci * P * k_m
    dR_dt = r_rt * active - (R - r_0) * r_rd
    dP_dt = k_p * ((R - r_0) - P)

    dH_dt = kk * jnp.maximum(R - z, jnp.array([0.0], dtype=float)) + h_b

    return dCe_dt, dCi_dt, dR_dt, dP_dt, dH_dt
```

##### S3.1.2 Solver post processing

```python
def survival(results, t, interpolation):
    results["survival"] = jnp.exp(-results["H"])
    return results
```

##### S3.1.3 Probability model

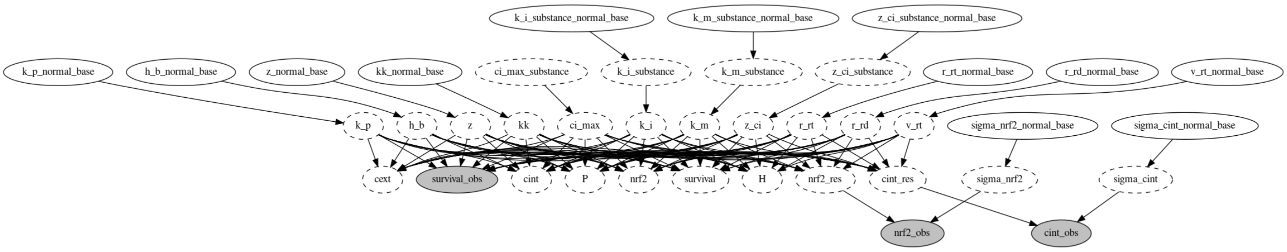


**Figure S1.** Directed acyclic graph (DAG) of the probability model.

#### S3.2 Report: Parameters ✓

### S3.2.1 $x_{in}$

No model input

### S3.2.2 $y_0$

| id | cext | cint | nrf2 | P | H |
|---|---|---|---|---|---|
| 101_0 | 2.34 | 0 | 1 | 0 | 0 |
| 101_1 | 2.34 | 0 | 1 | 0 | 0 |
| 106_0 | 5.16 | 0 | 1 | 0 | 0 |
| 106_1 | 5.16 | 0 | 1 | 0 | 0 |
| 112_0 | 11.72 | 0 | 1 | 0 | 0 |
| 112_1 | 11.72 | 0 | 1 | 0 | 0 |
| 118_0 | 18.14 | 0 | 1 | 0 | 0 |
| 118_1 | 18.14 | 0 | 1 | 0 | 0 |
| 124_0 | 29.44 | 0 | 1 | 0 | 0 |
| 124_1 | 29.44 | 0 | 1 | 0 | 0 |
| 184_0 | 2.12727 | 0 | 1 | 0 | 0 |
| 185_0 | 8.5091 | 0 | 1 | 0 | 0 |
| 186_0 | 10.6364 | 0 | 1 | 0 | 0 |
| 187_0 | 12.7636 | 0 | 1 | 0 | 0 |
| 188_0 | 14.8909 | 0 | 1 | 0 | 0 |
| 189_0 | 17.0182 | 0 | 1 | 0 | 0 |
| 190_0 | 25.5273 | 0 | 1 | 0 | 0 |
| 191_0 | 34.0364 | 0 | 1 | 0 | 0 |
| 192_0 | 45.7364 | 0 | 1 | 0 | 0 |
| 193_0 | 5.31819 | 0 | 1 | 0 | 0 |
| 194_0 | 6.38182 | 0 | 1 | 0 | 0 |
| 195_0 | 7.78583 | 0 | 1 | 0 | 0 |
| 196_0 | 9.31746 | 0 | 1 | 0 | 0 |
| 197_0 | 11.232 | 0 | 1 | 0 | 0 |
| 198_0 | 13.4869 | 0 | 1 | 0 | 0 |
| 199_0 | 15.7418 | 0 | 1 | 0 | 0 |

| id | cext | cint | nrf2 | P | H |
|---|---|---|---|---|---|
| 200_0 | 19.3582 | 0 | 1 | 0 | 0 |
| 201_0 | 23.2724 | 0 | 1 | 0 | 0 |
| 202_0 | 27.9098 | 0 | 1 | 0 | 0 |
| 203_0 | 33.5259 | 0 | 1 | 0 | 0 |
| 204_0 | 40.2055 | 0 | 1 | 0 | 0 |
| 205_0 | 48.2466 | 0 | 1 | 0 | 0 |
| 206_0 | 57.9044 | 0 | 1 | 0 | 0 |
| 207_0 | 69.4768 | 0 | 1 | 0 | 0 |
| 208_0 | 83.3892 | 0 | 1 | 0 | 0 |
| 209_0 | 2.08473 | 0 | 1 | 0 | 0 |
| 210_0 | 3.19091 | 0 | 1 | 0 | 0 |
| 211_0 | 4.7651 | 0 | 1 | 0 | 0 |
| 212_0 | 7.19019 | 0 | 1 | 0 | 0 |
| 213_0 | 8.5091 | 0 | 1 | 0 | 0 |
| 214_0 | 10.764 | 0 | 1 | 0 | 0 |
| 215_0 | 12.7636 | 0 | 1 | 0 | 0 |
| 216_0 | 14.8909 | 0 | 1 | 0 | 0 |
| 217_0 | 16.1247 | 0 | 1 | 0 | 0 |
| 218_0 | 24.1658 | 0 | 1 | 0 | 0 |
| 219_0 | 36.2913 | 0 | 1 | 0 | 0 |
| 220_0 | 54.4582 | 0 | 1 | 0 | 0 |
| 221_0 | 81.6874 | 0 | 1 | 0 | 0 |
| 222_0 | 8.01201 | 0 | 1 | 0 | 0 |
| 223_0 | 10.4156 | 0 | 1 | 0 | 0 |
| 224_0 | 13.5403 | 0 | 1 | 0 | 0 |
| 225_0 | 17.6024 | 0 | 1 | 0 | 0 |
| 226_0 | 22.8831 | 0 | 1 | 0 | 0 |
| 227_0 | 29.748 | 0 | 1 | 0 | 0 |
| 228_0 | 38.6724 | 0 | 1 | 0 | 0 |
| 229_0 | 50.2742 | 0 | 1 | 0 | 0 |
| 230_0 | 65.3564 | 0 | 1 | 0 | 0 |
| 231_0 | 84.9634 | 0 | 1 | 0 | 0 |
| 232_0 | 91.2601 | 0 | 1 | 0 | 0 |
| 42_0 | 23.3532 | 0 | 1 | 0 | 0 |
| 44_0 | 29.462 | 0 | 1 | 0 | 0 |
| 44_1 | 29.462 | 0 | 1 | 0 | 0 |
| 44_2 | 29.462 | 0 | 1 | 0 | 0 |
| 44_3 | 29.462 | 0 | 1 | 0 | 0 |
| 44_4 | 29.462 | 0 | 1 | 0 | 0 |
| 44_5 | 29.462 | 0 | 1 | 0 | 0 |
| 44_6 | 29.462 | 0 | 1 | 0 | 0 |
| 44_7 | 29.462 | 0 | 1 | 0 | 0 |
| 51_0 | 20.8579 | 0 | 1 | 0 | 0 |
| 51_1 | 20.9121 | 0 | 1 | 0 | 0 |
| 51_2 | 20.9456 | 0 | 1 | 0 | 0 |
| 52_0 | 19.9914 | 0 | 1 | 0 | 0 |
| 52_1 | 19.9914 | 0 | 1 | 0 | 0 |
| 52_2 | 19.9914 | 0 | 1 | 0 | 0 |
| 53_0 | 19.8758 | 0 | 1 | 0 | 0 |
| 53_1 | 20.0048 | 0 | 1 | 0 | 0 |

| id | cext | cint | nrf2 | P | H |
|---|---|---|---|---|---|
| 69_0 | 18.7886 | 0 | 1 | 0 | 0 |
| 70_0 | 19.0905 | 0 | 1 | 0 | 0 |
| 70_1 | 18.943 | 0 | 1 | 0 | 0 |
| 70_2 | 19.9914 | 0 | 1 | 0 | 0 |
| 74_0 | 19.9914 | 0 | 1 | 0 | 0 |
| 74_1 | 19.9914 | 0 | 1 | 0 | 0 |
| 74_2 | 19.9914 | 0 | 1 | 0 | 0 |
| 75_0 | 20.5062 | 0 | 1 | 0 | 0 |
| 8_0 | 14.0585 | 0 | 1 | 0 | 0 |
| 10_0 | 12.1997 | 0 | 1 | 0 | 0 |
| 126_0 | 0.878067 | 0 | 1 | 0 | 0 |
| 127_0 | 1.75613 | 0 | 1 | 0 | 0 |
| 128_0 | 3.51227 | 0 | 1 | 0 | 0 |
| 129_0 | 7.02453 | 0 | 1 | 0 | 0 |
| 130_0 | 14.0491 | 0 | 1 | 0 | 0 |
| 131_0 | 28.0981 | 0 | 1 | 0 | 0 |
| 132_0 | 56.1963 | 0 | 1 | 0 | 0 |
| 133_0 | 112.393 | 0 | 1 | 0 | 0 |
| 134_0 | 224.785 | 0 | 1 | 0 | 0 |
| 136_0 | 449.57 | 0 | 1 | 0 | 0 |
| 138_0 | 3.29335 | 0 | 1 | 0 | 0 |
| 139_0 | 4.28135 | 0 | 1 | 0 | 0 |
| 140_0 | 5.56576 | 0 | 1 | 0 | 0 |
| 141_0 | 7.23549 | 0 | 1 | 0 | 0 |
| 142_0 | 9.40613 | 0 | 1 | 0 | 0 |
| 143_0 | 12.228 | 0 | 1 | 0 | 0 |
| 144_0 | 15.8964 | 0 | 1 | 0 | 0 |
| 145_0 | 20.6653 | 0 | 1 | 0 | 0 |
| 146_0 | 26.8649 | 0 | 1 | 0 | 0 |
| 147_0 | 34.9243 | 0 | 1 | 0 | 0 |
| 148_0 | 7.99998 | 0 | 1 | 0 | 0 |
| 149_0 | 9.59998 | 0 | 1 | 0 | 0 |
| 150_0 | 11.52 | 0 | 1 | 0 | 0 |
| 151_0 | 13.824 | 0 | 1 | 0 | 0 |
| 152_0 | 16.5888 | 0 | 1 | 0 | 0 |
| 153_0 | 19.9065 | 0 | 1 | 0 | 0 |
| 154_0 | 23.8878 | 0 | 1 | 0 | 0 |
| 155_0 | 28.6654 | 0 | 1 | 0 | 0 |
| 156_0 | 34.3985 | 0 | 1 | 0 | 0 |
| 157_0 | 41.2781 | 0 | 1 | 0 | 0 |
| 158_0 | 3.66326 | 0 | 1 | 0 | 0 |
| 159_0 | 4.57907 | 0 | 1 | 0 | 0 |
| 160_0 | 5.72384 | 0 | 1 | 0 | 0 |
| 161_0 | 7.1548 | 0 | 1 | 0 | 0 |
| 162_0 | 8.94349 | 0 | 1 | 0 | 0 |
| 163_0 | 11.1794 | 0 | 1 | 0 | 0 |
| 164_0 | 13.9742 | 0 | 1 | 0 | 0 |
| 165_0 | 17.4678 | 0 | 1 | 0 | 0 |
| 166_0 | 21.8347 | 0 | 1 | 0 | 0 |
| 167_0 | 27.2934 | 0 | 1 | 0 | 0 |

| id | cext | cint | nrf2 | P | H |
|---|---|---|---|---|---|
| 168_0 | 34.1167 | 0 | 1 | 0 | 0 |
| 169_0 | 42.6459 | 0 | 1 | 0 | 0 |
| 170_0 | 53.3074 | 0 | 1 | 0 | 0 |
| 171_0 | 3.14673 | 0 | 1 | 0 | 0 |
| 172_0 | 3.77607 | 0 | 1 | 0 | 0 |
| 173_0 | 4.53128 | 0 | 1 | 0 | 0 |
| 174_0 | 5.43754 | 0 | 1 | 0 | 0 |
| 175_0 | 6.52505 | 0 | 1 | 0 | 0 |
| 176_0 | 7.83006 | 0 | 1 | 0 | 0 |
| 177_0 | 9.39607 | 0 | 1 | 0 | 0 |
| 178_0 | 11.2753 | 0 | 1 | 0 | 0 |
| 179_0 | 13.5303 | 0 | 1 | 0 | 0 |
| 180_0 | 16.2364 | 0 | 1 | 0 | 0 |
| 181_0 | 19.4837 | 0 | 1 | 0 | 0 |
| 182_0 | 23.3804 | 0 | 1 | 0 | 0 |
| 183_0 | 28.0565 | 0 | 1 | 0 | 0 |
| 38_0 | 9.22851 | 0 | 1 | 0 | 0 |
| 43_0 | 6.60108 | 0 | 1 | 0 | 0 |
| 43_1 | 6.60108 | 0 | 1 | 0 | 0 |
| 43_2 | 6.60108 | 0 | 1 | 0 | 0 |
| 43_3 | 6.60108 | 0 | 1 | 0 | 0 |
| 54_0 | 4.25084 | 0 | 1 | 0 | 0 |
| 54_1 | 3.93463 | 0 | 1 | 0 | 0 |
| 55_0 | 4.83931 | 0 | 1 | 0 | 0 |
| 55_1 | 4.77683 | 0 | 1 | 0 | 0 |
| 55_2 | 6.60108 | 0 | 1 | 0 | 0 |
| 56_0 | 6.5343 | 0 | 1 | 0 | 0 |
| 56_1 | 6.30297 | 0 | 1 | 0 | 0 |
| 56_2 | 6.36656 | 0 | 1 | 0 | 0 |
| 56_3 | 3.64386 | 0 | 1 | 0 | 0 |
| 56_4 | 4.75085 | 0 | 1 | 0 | 0 |
| 56_5 | 5.17322 | 0 | 1 | 0 | 0 |
| 57_0 | 6.21088 | 0 | 1 | 0 | 0 |
| 57_1 | 6.49514 | 0 | 1 | 0 | 0 |
| 57_2 | 6.56835 | 0 | 1 | 0 | 0 |
| 57_3 | 4.30287 | 0 | 1 | 0 | 0 |
| 57_4 | 4.853 | 0 | 1 | 0 | 0 |
| 57_5 | 5.56204 | 0 | 1 | 0 | 0 |
| 62_0 | 5.42553 | 0 | 1 | 0 | 0 |
| 62_1 | 5.1399 | 0 | 1 | 0 | 0 |
| 67_0 | 7.4669 | 0 | 1 | 0 | 0 |
| 67_1 | 8.07566 | 0 | 1 | 0 | 0 |
| 67_2 | 9.49718 | 0 | 1 | 0 | 0 |
| 67_3 | 7.83615 | 0 | 1 | 0 | 0 |
| 67_4 | 6.60108 | 0 | 1 | 0 | 0 |
| 67_5 | 6.60108 | 0 | 1 | 0 | 0 |
| 68_0 | 8.57659 | 0 | 1 | 0 | 0 |
| 68_1 | 8.33878 | 0 | 1 | 0 | 0 |
| 68_2 | 7.22975 | 0 | 1 | 0 | 0 |
| 68_3 | 7.22975 | 0 | 1 | 0 | 0 |

| id | cext | cint | nrf2 | P | H |
|---|---|---|---|---|---|
| 71_0 | 5.02939 | 0 | 1 | 0 | 0 |
| 72_0 | 6.60108 | 0 | 1 | 0 | 0 |
| 72_1 | 6.60108 | 0 | 1 | 0 | 0 |
| 72_2 | 6.60108 | 0 | 1 | 0 | 0 |
| 72_3 | 6.60108 | 0 | 1 | 0 | 0 |
| 72_4 | 6.60108 | 0 | 1 | 0 | 0 |
| 72_5 | 6.60108 | 0 | 1 | 0 | 0 |
| 73_0 | 7.22975 | 0 | 1 | 0 | 0 |
| 73_1 | 7.22975 | 0 | 1 | 0 | 0 |
| 73_2 | 7.22975 | 0 | 1 | 0 | 0 |
| 78_0 | 7.10197 | 0 | 1 | 0 | 0 |
| 78_1 | 6.60108 | 0 | 1 | 0 | 0 |
| 80_0 | 6.86279 | 0 | 1 | 0 | 0 |
| 82_0 | 5.1 | 0 | 1 | 0 | 0 |
| 82_1 | 5.1 | 0 | 1 | 0 | 0 |
| 84_0 | 5.77 | 0 | 1 | 0 | 0 |
| 84_1 | 5.77 | 0 | 1 | 0 | 0 |
| 86_0 | 6.52 | 0 | 1 | 0 | 0 |
| 86_1 | 6.52 | 0 | 1 | 0 | 0 |
| 88_0 | 6.93 | 0 | 1 | 0 | 0 |
| 88_1 | 6.93 | 0 | 1 | 0 | 0 |
| 88_2 | 6.93 | 0 | 1 | 0 | 0 |
| 90_0 | 7.36 | 0 | 1 | 0 | 0 |
| 90_1 | 7.36 | 0 | 1 | 0 | 0 |
| 91_0 | 5.1 | 0 | 1 | 0 | 0 |
| 91_1 | 5.1 | 0 | 1 | 0 | 0 |
| 92_0 | 5.77 | 0 | 1 | 0 | 0 |
| 92_1 | 5.77 | 0 | 1 | 0 | 0 |
| 93_0 | 6.52 | 0 | 1 | 0 | 0 |
| 93_1 | 6.52 | 0 | 1 | 0 | 0 |
| 94_0 | 6.93 | 0 | 1 | 0 | 0 |
| 94_1 | 6.93 | 0 | 1 | 0 | 0 |
| 95_0 | 7.36 | 0 | 1 | 0 | 0 |
| 102_0 | 134.58 | 0 | 1 | 0 | 0 |
| 102_1 | 134.58 | 0 | 1 | 0 | 0 |
| 107_0 | 177.57 | 0 | 1 | 0 | 0 |
| 113_0 | 234.29 | 0 | 1 | 0 | 0 |
| 113_1 | 234.29 | 0 | 1 | 0 | 0 |
| 119_0 | 269.13 | 0 | 1 | 0 | 0 |
| 119_1 | 269.13 | 0 | 1 | 0 | 0 |
| 125_0 | 309.14 | 0 | 1 | 0 | 0 |
| 125_1 | 309.14 | 0 | 1 | 0 | 0 |
| 233_0 | 10.5849 | 0 | 1 | 0 | 0 |
| 234_0 | 21.1698 | 0 | 1 | 0 | 0 |
| 235_0 | 42.3396 | 0 | 1 | 0 | 0 |
| 236_0 | 84.6793 | 0 | 1 | 0 | 0 |
| 237_0 | 169.359 | 0 | 1 | 0 | 0 |
| 238_0 | 338.717 | 0 | 1 | 0 | 0 |
| 239_0 | 677.434 | 0 | 1 | 0 | 0 |
| 240_0 | 1354.87 | 0 | 1 | 0 | 0 |

| id | cext | cint | nrf2 | P | H |
|---|---|---|---|---|---|
| 241_0 | 281.571 | 0 | 1 | 0 | 0 |
| 242_0 | 337.886 | 0 | 1 | 0 | 0 |
| 243_0 | 405.463 | 0 | 1 | 0 | 0 |
| 244_0 | 486.556 | 0 | 1 | 0 | 0 |
| 245_0 | 583.867 | 0 | 1 | 0 | 0 |
| 247_0 | 700.64 | 0 | 1 | 0 | 0 |
| 249_0 | 840.768 | 0 | 1 | 0 | 0 |
| 251_0 | 1008.92 | 0 | 1 | 0 | 0 |
| 253_0 | 1210.71 | 0 | 1 | 0 | 0 |
| 254_0 | 1452.85 | 0 | 1 | 0 | 0 |
| 255_0 | 137.422 | 0 | 1 | 0 | 0 |
| 256_0 | 164.906 | 0 | 1 | 0 | 0 |
| 257_0 | 197.888 | 0 | 1 | 0 | 0 |
| 258_0 | 237.465 | 0 | 1 | 0 | 0 |
| 259_0 | 284.958 | 0 | 1 | 0 | 0 |
| 260_0 | 341.95 | 0 | 1 | 0 | 0 |
| 261_0 | 410.34 | 0 | 1 | 0 | 0 |
| 262_0 | 492.408 | 0 | 1 | 0 | 0 |
| 27_0 | 238.256 | 0 | 1 | 0 | 0 |
| 28_0 | 385.144 | 0 | 1 | 0 | 0 |
| 33_0 | 200.606 | 0 | 1 | 0 | 0 |
| 40_0 | 341.171 | 0 | 1 | 0 | 0 |
| 48_0 | 134.792 | 0 | 1 | 0 | 0 |
| 48_1 | 134.792 | 0 | 1 | 0 | 0 |
| 48_2 | 134.792 | 0 | 1 | 0 | 0 |
| 48_3 | 134.792 | 0 | 1 | 0 | 0 |
| 48_4 | 134.792 | 0 | 1 | 0 | 0 |
| 48_5 | 134.792 | 0 | 1 | 0 | 0 |
| 48_6 | 134.792 | 0 | 1 | 0 | 0 |
| 49_0 | 309.229 | 0 | 1 | 0 | 0 |
| 49_1 | 309.229 | 0 | 1 | 0 | 0 |
| 49_2 | 309.229 | 0 | 1 | 0 | 0 |
| 49_3 | 309.229 | 0 | 1 | 0 | 0 |
| 49_4 | 309.229 | 0 | 1 | 0 | 0 |
| 49_5 | 309.229 | 0 | 1 | 0 | 0 |
| 49_6 | 309.229 | 0 | 1 | 0 | 0 |
| 58_0 | 127.212 | 0 | 1 | 0 | 0 |
| 58_1 | 129.451 | 0 | 1 | 0 | 0 |
| 58_2 | 130.777 | 0 | 1 | 0 | 0 |
| 59_0 | 295.007 | 0 | 1 | 0 | 0 |
| 59_1 | 294.15 | 0 | 1 | 0 | 0 |
| 59_2 | 299.496 | 0 | 1 | 0 | 0 |
| 5_0 | 291.323 | 0 | 1 | 0 | 0 |
| 60_0 | 133.312 | 0 | 1 | 0 | 0 |
| 60_1 | 121.34 | 0 | 1 | 0 | 0 |
| 61_0 | 320.009 | 0 | 1 | 0 | 0 |
| 61_1 | 235.269 | 0 | 1 | 0 | 0 |
| 63_0 | 145.646 | 0 | 1 | 0 | 0 |
| 63_1 | 131.266 | 0 | 1 | 0 | 0 |
| 63_2 | 134.792 | 0 | 1 | 0 | 0 |

| id | cext | cint | nrf2 | P | H |
|---|---|---|---|---|---|
| 63_3 | 134.792 | 0 | 1 | 0 | 0 |
| 64_0 | 475.162 | 0 | 1 | 0 | 0 |
| 64_1 | 411.483 | 0 | 1 | 0 | 0 |
| 64_2 | 309.229 | 0 | 1 | 0 | 0 |
| 64_3 | 309.229 | 0 | 1 | 0 | 0 |
| 65_0 | 208.492 | 0 | 1 | 0 | 0 |
| 65_1 | 164.104 | 0 | 1 | 0 | 0 |
| 65_2 | 162.544 | 0 | 1 | 0 | 0 |
| 65_3 | 134.792 | 0 | 1 | 0 | 0 |
| 65_4 | 134.792 | 0 | 1 | 0 | 0 |
| 65_5 | 134.792 | 0 | 1 | 0 | 0 |
| 66_0 | 505.273 | 0 | 1 | 0 | 0 |
| 66_1 | 456.728 | 0 | 1 | 0 | 0 |
| 66_2 | 514.382 | 0 | 1 | 0 | 0 |
| 66_3 | 309.229 | 0 | 1 | 0 | 0 |
| 66_4 | 309.229 | 0 | 1 | 0 | 0 |
| 66_5 | 309.229 | 0 | 1 | 0 | 0 |
| 6_0 | 349.539 | 0 | 1 | 0 | 0 |
| 76_0 | 134.792 | 0 | 1 | 0 | 0 |
| 76_1 | 134.792 | 0 | 1 | 0 | 0 |
| 76_10 | 134.792 | 0 | 1 | 0 | 0 |
| 76_2 | 134.792 | 0 | 1 | 0 | 0 |
| 76_3 | 134.792 | 0 | 1 | 0 | 0 |
| 76_4 | 134.792 | 0 | 1 | 0 | 0 |
| 76_5 | 134.792 | 0 | 1 | 0 | 0 |
| 76_6 | 134.792 | 0 | 1 | 0 | 0 |
| 76_7 | 134.792 | 0 | 1 | 0 | 0 |
| 76_8 | 134.792 | 0 | 1 | 0 | 0 |
| 76_9 | 134.792 | 0 | 1 | 0 | 0 |
| 77_0 | 309.229 | 0 | 1 | 0 | 0 |
| 77_1 | 309.229 | 0 | 1 | 0 | 0 |
| 77_2 | 309.229 | 0 | 1 | 0 | 0 |
| 77_3 | 309.229 | 0 | 1 | 0 | 0 |
| 77_4 | 309.229 | 0 | 1 | 0 | 0 |
| 77_5 | 309.229 | 0 | 1 | 0 | 0 |
| 77_6 | 309.229 | 0 | 1 | 0 | 0 |
| 77_7 | 309.229 | 0 | 1 | 0 | 0 |
| 77_8 | 309.229 | 0 | 1 | 0 | 0 |

##### S3.2.3 Free parameters

- k_i_substance ∼ lognorm(scale=[1.0,1.0,1.0],s=2,dims=('substance',))
- k_m_substance ∼ lognorm(scale=[0.05,0.05,0.05],s=2,dims=('substance',))
- z_ci_substance ∼ lognorm(scale=[0.5,0.5,0.5],s=2,dims=('substance',))
- r_rt ∼ lognorm(scale=1.0,s=2,dims=())
- r_rd ∼ lognorm(scale=0.5,s=2,dims=())
- v_rt ∼ lognorm(scale=1.0,s=2,dims=())
- k_p ∼ lognorm(scale=0.02,s=2,dims=())
- h_b ∼ lognorm(scale=1e-08,s=2,dims=())
- z ∼ lognorm(scale=1.0,s=2,dims=())

- kk ~ lognorm(scale=0.02,s=2,dims=())
- sigma_cint ~ halfnorm(scale=5.0,dims=())
- sigma_nrf2 ~ halfnorm(scale=5.0,dims=())

###### S3.2.4 Fixed parameters

- k_i = k_i_substance[substance_index], dims=('id',)
- k_m = k_m_substance[substance_index], dims=('id',)
- z_ci = z_ci_substance[substance_index], dims=('id',)
- ci_max_substance = [1757.0, 168.1, 6364.8], dims=('substance',)
- ci_max = ci_max_substance[substance_index], dims=('id',)
- r_0 = 1.0, dims=()

##### S3.3 Report: Table parameter estimates ✓

| | ('index', ' ') | ('diuron', 'mean ± std') | ('diclofenac', 'mean ± std') | ('naproxen', 'mean ± std') |
|---|---|---|---|---|
| 0 | ci_max_substance | 1757.0 ± 0.0 | 168.1 ± 0.0 | 6364.8 ± 0.0 |
| 1 | h_b | 0.0 ± 0.0 | 0.0 ± 0.0 | 0.0 ± 0.0 |
| 2 | k_i_substance | 13.97 ± 2.493 | 0.38 ± 0.022 | 0.283 ± 0.016 |
| 3 | k_m_substance | 1.785 ± 0.441 | 0.088 ± 0.013 | 0.031 ± 0.005 |
| 4 | k_p | 0.025 ± 0.005 | 0.025 ± 0.005 | 0.025 ± 0.005 |
| 5 | kk | 0.043 ± 0.006 | 0.043 ± 0.006 | 0.043 ± 0.006 |
| 6 | r_rd | 0.138 ± 0.017 | 0.138 ± 0.017 | 0.138 ± 0.017 |
| 7 | r_rt | 0.399 ± 0.045 | 0.399 ± 0.045 | 0.399 ± 0.045 |
| 8 | sigma_cint | 0.787 ± 0.026 | 0.787 ± 0.026 | 0.787 ± 0.026 |
| 9 | sigma_nrf2 | 0.298 ± 0.01 | 0.298 ± 0.01 | 0.298 ± 0.01 |
| 10 | v_rt | 5.681 ± 0.475 | 5.681 ± 0.475 | 5.681 ± 0.475 |
| 11 | z | 2.238 ± 0.079 | 2.238 ± 0.079 | 2.238 ± 0.079 |
| 12 | z_ci_substance | 0.627 ± 0.055 | 0.38 ± 0.032 | 0.495 ± 0.039 |

Report 'table_parameter_estimates' was successfully generated and saved in '../tktd_rna_pulse/results/rna_pulse_5_

##### S3.4 Report: Goodness of fit ✓

| | cint | nrf2 | survival | model |
|---|---|---|---|---|
| NRMSE | 0.122078 | 0.176929 | 0.35283 | nan |
| NRMSE (95%-hdi[lower]) | 0.11748 | 0.170892 | 0.341109 | nan |
| NRMSE (95%-hdi[upper]) | 0.128475 | 0.182609 | 0.366025 | nan |
| Log-Likelihood | -1055.13 | -37.6129 | -522.912 | -1615.65 |
| Log-Likelihood (95%-hdi[lower]) | -1071.03 | -40.5499 | -533.442 | -1629.05 |
| Log-Likelihood (95%-hdi[upper]) | -1036.87 | -35.1349 | -512.258 | -1602.07 |
| n (data) | 913 | 169 | 392 | 1474 |
| k (parameters) | nan | nan | nan | 18 |
| BIC | nan | nan | nan | 3362.63 |
| BIC (95%-hdi[lower]) | nan | nan | nan | 3335.46 |
| BIC (95%-hdi[upper]) | nan | nan | nan | 3389.42 |

Report 'goodness_of_fit' was successfully generated and saved in '../tktd_rna_pulse/results/rna_pulse_5_substance_i

#### S3.5 Report: Diagnostics ✓

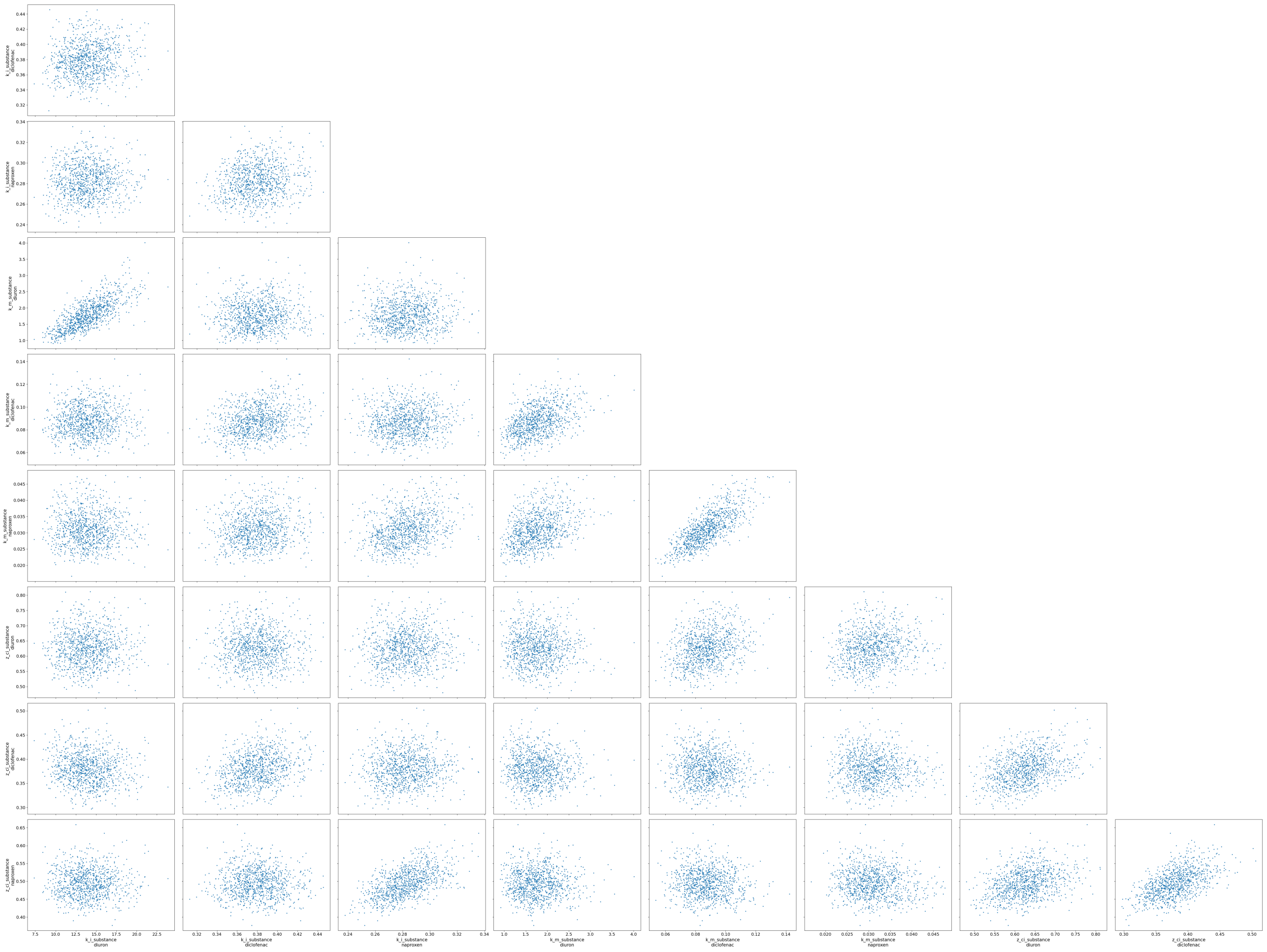

**Figure S2.** Paired parameter estimates

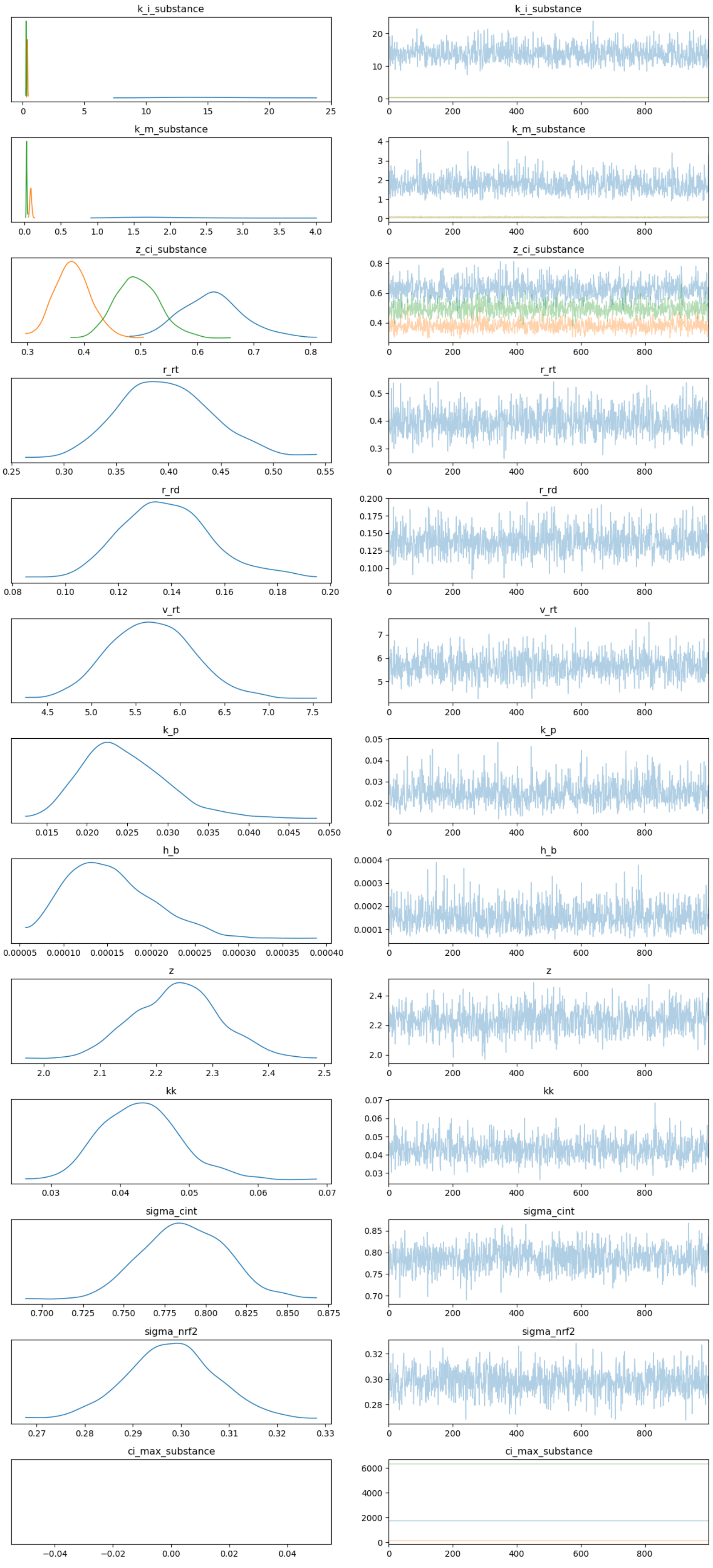


**Figure S3.** Psuedo trace, generated for draws from the optimized SVI distribution

Report ‘diagnostics’ was successfully generated and saved in ‘(’../tktd_rna_pulse/results/rna_pulse_5_substance_indep[illegible]
‘../tktd_rna_pulse/results/rna_pulse_5_substance_independent_rna_protein_module_full_dataset/posterior_trace.png’)’

##### S3.6 Report: Visualizations ✓

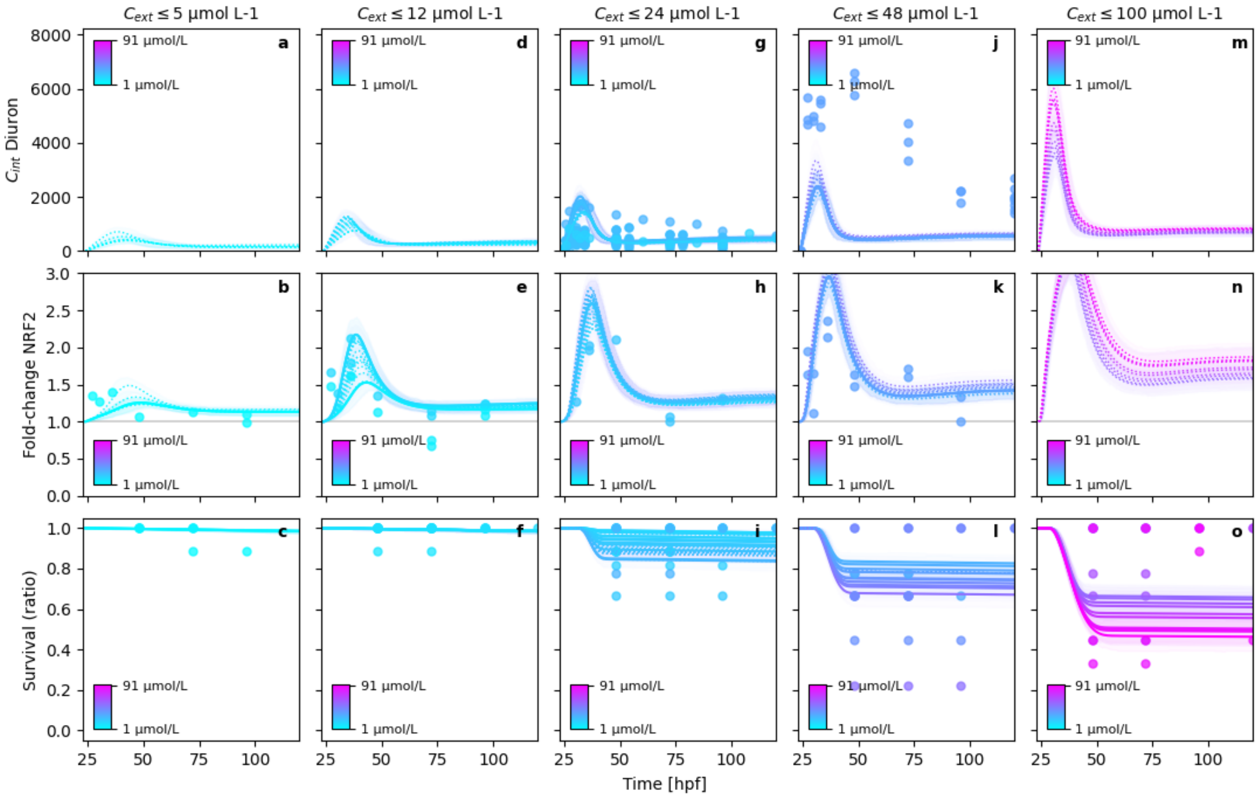


**Figure S4.** Posterior model fits

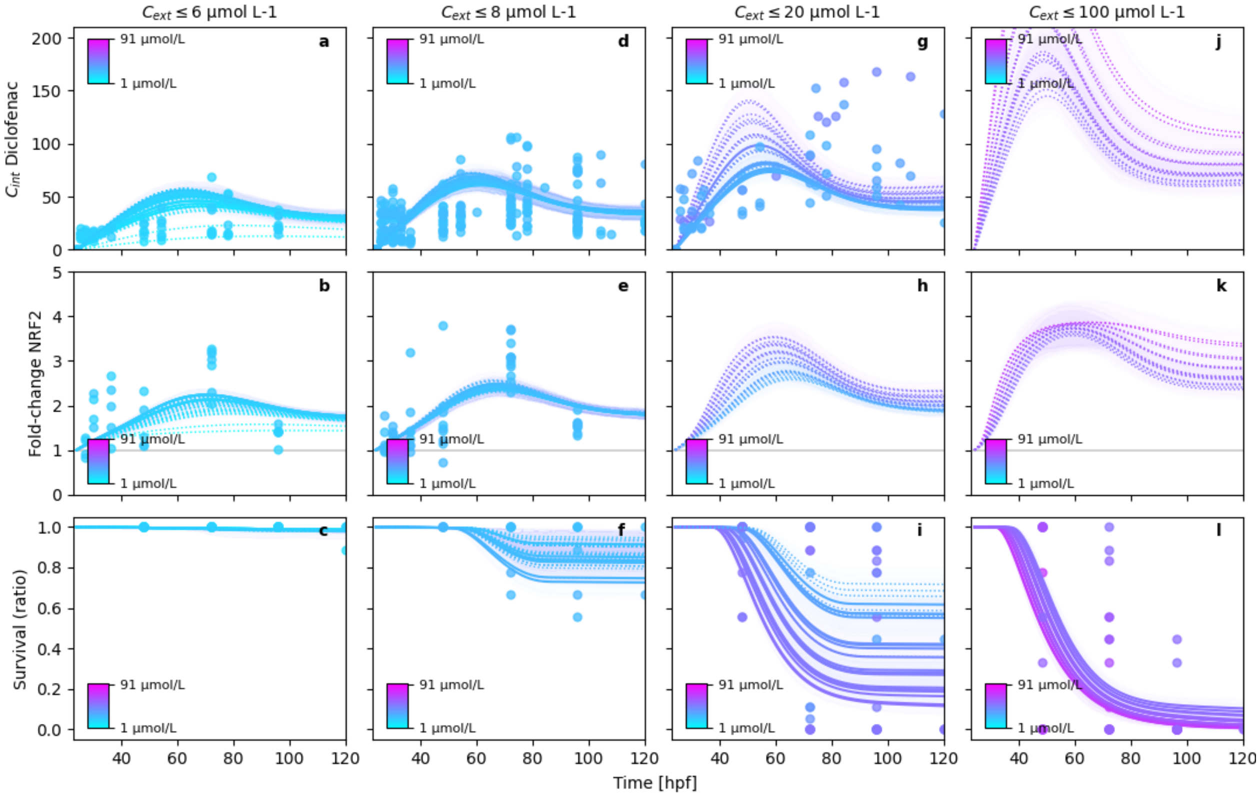


**Figure S5.** Posterior model fits

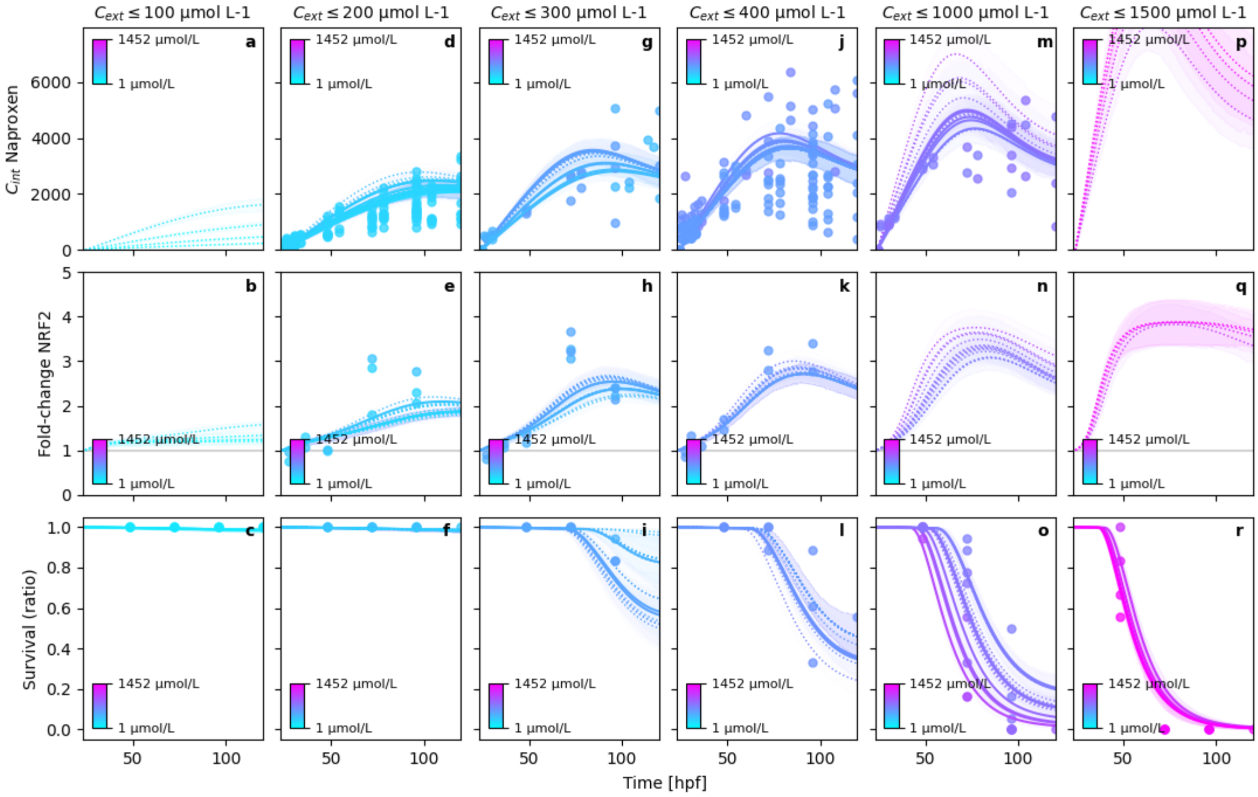


**Figure S6.** Posterior model fits

Report 'visualizations' was successfully generated and saved in '['../tktd_rna_pulse/results/rna_pulse_5_substance_inde
'../tktd_rna_pulse/results/rna_pulse_5_substance_independent_rna_protein_module_full_dataset/combined_pps_figure_

'../tktd_rna_pulse/results/rna_pulse_5_substance_independent_rna_protein_module_full_dataset/combined_pps_figure_

##### S3.7 Report: Model inadequacy metrics ✓

###### S3.7.1 Residuals

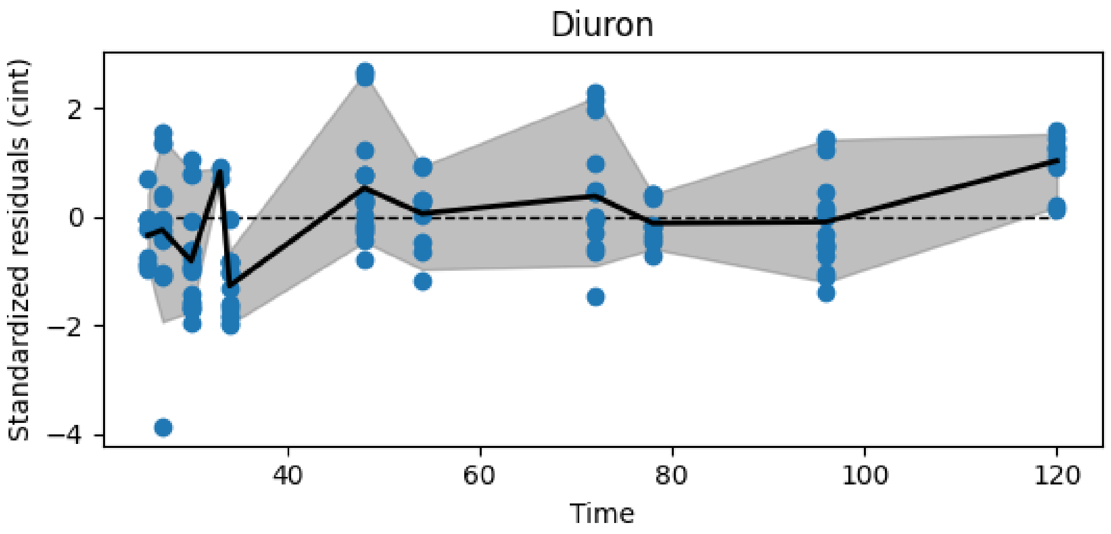


**Figure S7.** Resiudal cint dynamics of diuron

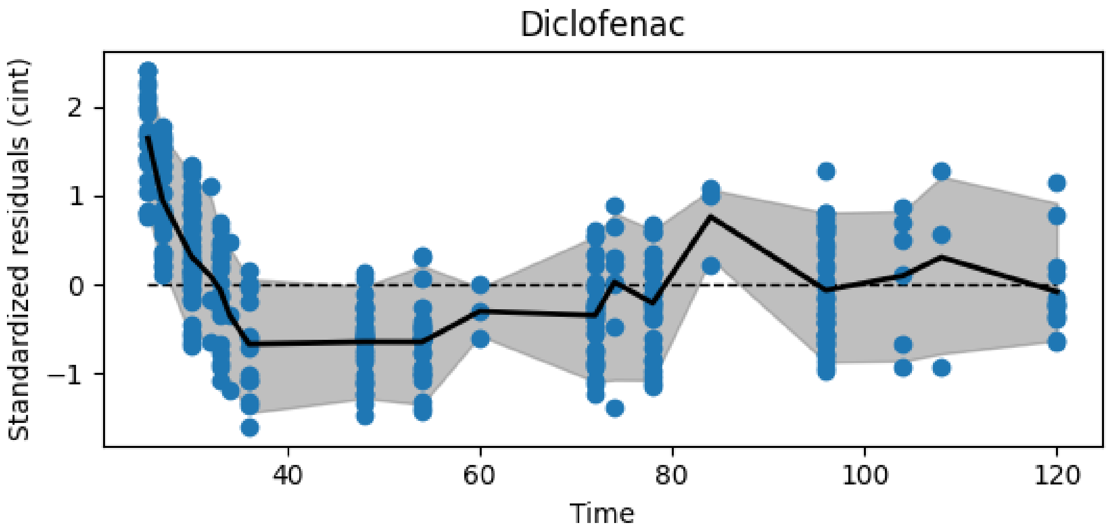


**Figure S8.** Resiudal cint dynamics of diclofenac

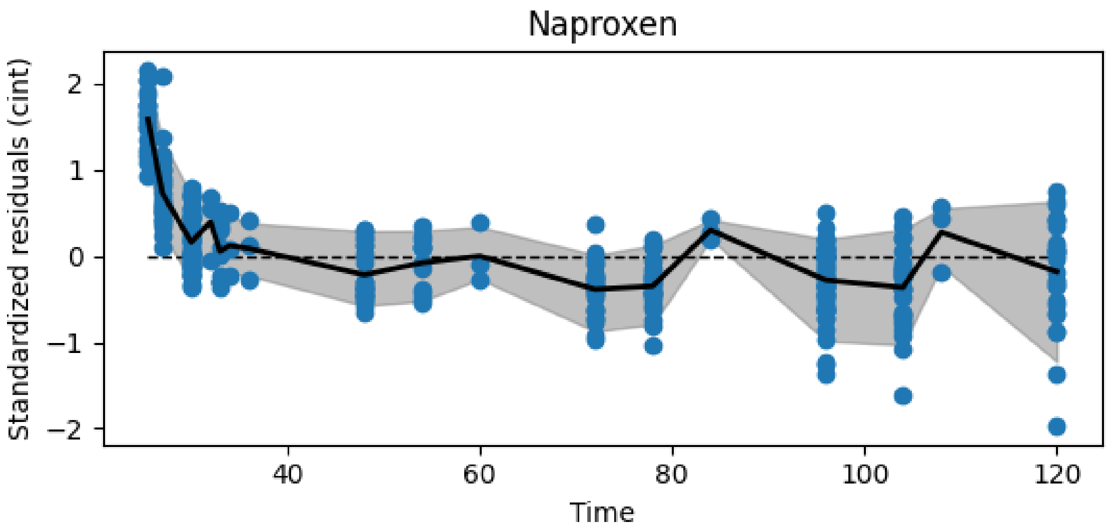


**Figure S9.** Resiudal cint dynamics of naproxen

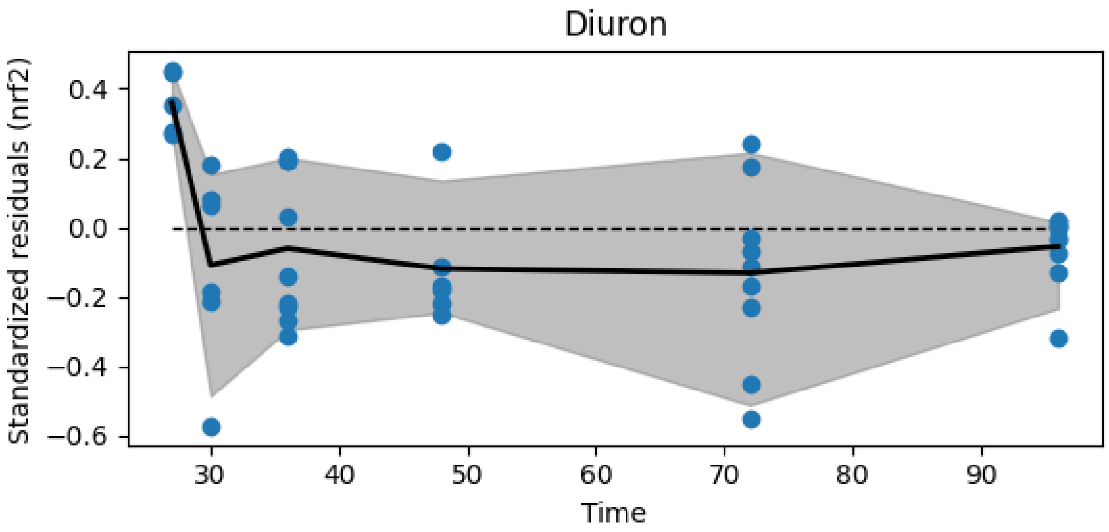


**Figure S10.** Resiudal nrf2 dynamics of diuron

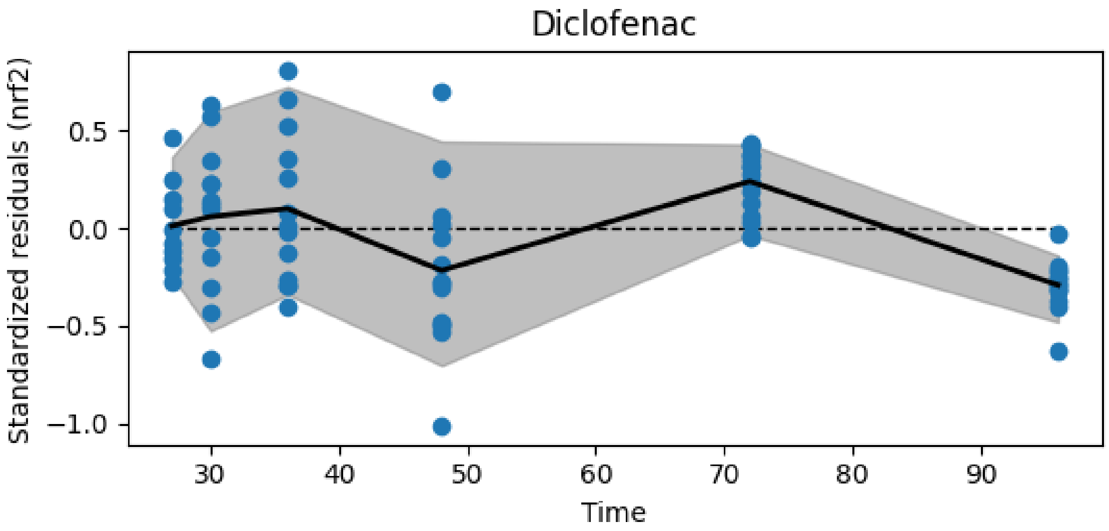


**Figure S11.** Resiudal nrf2 dynamics of diclofenac

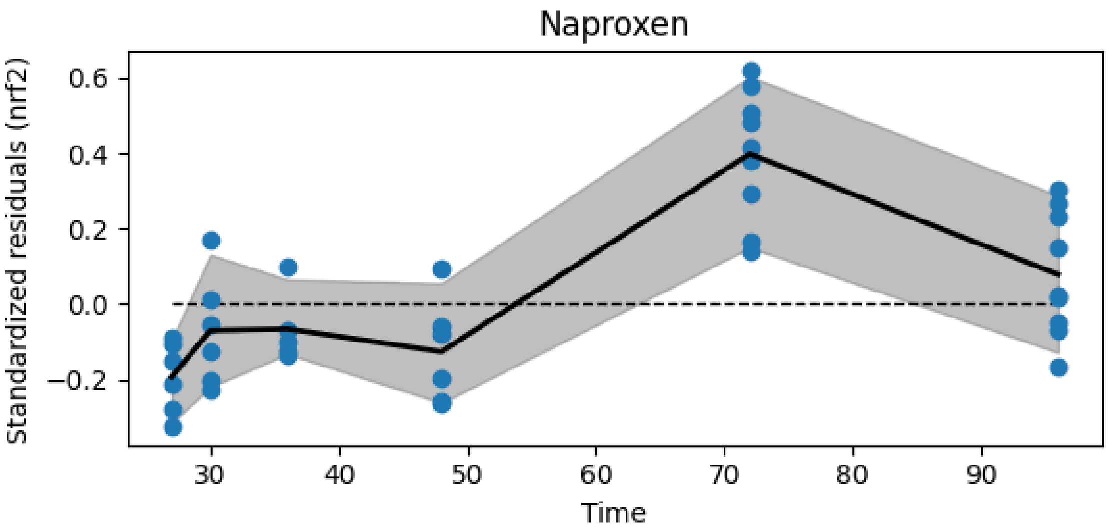


**Figure S12.** Resiudal nrf2 dynamics of naproxen

##### S3.7.2 Model inadequacy

The different metrics are measures for the model inadequacy. The comparison to `metric_value_if_normal_dist` is a simulation of normally distributed residuals that have the same data structure in terms of dimensionality (id x time) and missing values. If the `metric_value` falls within that interval, the model can be assumed as not inadequate.

- **autocorrelation**: Measures the correlation of the residuals with themselves with a lag of 1. High absolute autocorrelation means, the variable is not normally distributed. Ideal would be values close to 0.
- **deviation log-prob**: Uses a t-test to estimate the probability of the replicates at a time t being different from zero. The result is the summed log-probability. Low (negative) log probs indicate high probability for deviation.
- **significant deviations**: Uses a t-test to estimate the probability of the replicates at a time t being different from zero. The result is the number of significant deviations (for an alpha level of 0.05). High number of deviations indicate an inadequate model

- **local/global variance**: This metric calculates the local variance as a rolling variance of always 3 direct neighboring residuals. The local variances are then averaged and divided by the global averages of all residuals. The basis for the calculation is the residuals averaged by id. Values close to 1 indicate an adequate model
- **replicate/global variance**: This metric calculates the replicate variance at time t, averages it and divides the number by the global variance. The local variances are then averaged and divided by the global averages of all residuals. The basis for the calculation is the residuals averaged by id. Values close to 1 indicate an adequate model

| | metric | data_variable | index | metric_value | metric_value_if_normal_dist |
|---|---|---|---|---|---|
| 0 | autocorrelation | cint | diclofenac | 0.694237 | -0.04[-0.34,0.29] |
| 1 | autocorrelation | cint | diuron | -0.546305 | -0.12[-0.53,0.43] |
| 2 | autocorrelation | cint | naproxen | 0.532375 | -0.06[-0.38,0.3] |
| 3 | autocorrelation | nrf2 | diclofenac | -0.890841 | -0.25[-0.76,0.36] |
| 4 | autocorrelation | nrf2 | diuron | -0.272144 | -0.22[-0.79,0.51] |
| 5 | autocorrelation | nrf2 | naproxen | 0.0503981 | -0.16[-0.72,0.52] |
| 6 | deviation log-prob | cint | diclofenac | -127.32 | -18.03[-27.51,-11.16] |
| 7 | deviation log-prob | cint | diuron | -46.927 | -10.58[-16.2,-6.04] |
| 8 | deviation log-prob | cint | naproxen | -124.608 | -17.25[-24.09,-11.25] |
| 9 | deviation log-prob | nrf2 | diclofenac | -30.8218 | -6.15[-10.58,-2.52] |
| 10 | deviation log-prob | nrf2 | diuron | -14.618 | -5.4[-11.1,-2.54] |
| 11 | deviation log-prob | nrf2 | naproxen | -21.5593 | -6.31[-11.63,-2.7] |
| 12 | local/global variance | cint | diclofenac | 0.22326 | 0.73[0.51,0.89] |
| 13 | local/global variance | cint | diuron | 0.840779 | 0.75[0.44,1.0] |
| 14 | local/global variance | cint | naproxen | 0.289806 | 0.74[0.56,0.92] |
| 15 | local/global variance | nrf2 | diclofenac | 0.839068 | 0.85[0.54,1.21] |
| 16 | local/global variance | nrf2 | diuron | 0.39773 | 0.84[0.44,1.24] |
| 17 | local/global variance | nrf2 | naproxen | 0.697706 | 0.83[0.44,1.23] |
| 18 | replicate/global variance | cint | diclofenac | 0.469347 | 0.86[0.74,1.0] |
| 19 | replicate/global variance | cint | diuron | 0.527751 | 0.91[0.8,1.02] |
| 20 | replicate/global variance | cint | naproxen | 0.302548 | 0.84[0.71,1.01] |
| 21 | replicate/global variance | nrf2 | diclofenac | 0.717106 | 0.93[0.85,1.0] |
| 22 | replicate/global variance | nrf2 | diuron | 0.614125 | 0.9[0.76,1.04] |
| 23 | replicate/global variance | nrf2 | naproxen | 0.269133 | 0.85[0.72,1.0] |
| 24 | significant deviations | cint | diclofenac | 7 | 0.89[0.0,3.0] |
| 25 | significant deviations | cint | diuron | 5 | 0.39[0.0,1.0] |
| 26 | significant deviations | cint | naproxen | 8 | 0.9[0.0,2.0] |
| 27 | significant deviations | nrf2 | diclofenac | 2 | 0.33[0.0,1.0] |
| 28 | significant deviations | nrf2 | diuron | 1 | 0.27[0.0,1.0] |
| 29 | significant deviations | nrf2 | naproxen | 2 | 0.37[0.0,1.0] |

Report 'model_inadequacy_metrics' was successfully generated and saved in '../tktd_rna_pulse/results/rna_pulse_5

### S4 Report(case_study=hierarchical_molecular_tktd, scenario=hierarchical_ce[illegible]

- Using `hierarchical_molecular_tktd==0.1.6`
- Using `pymob==0.5.6a3`
- Using backend: `NumpyroBackend`
- Using settings: `../hierarchical_molecular_tktd/scenarios/hierarchical_cext_nested_sigma_hyperp`[illegible]

#### S4.1 Report: Model ✓

##### S4.1.1 Model

```
def tktd_rna_5(t, X, r_0, k_i, r_rt, r_rd, z_ci, v_rt, k_p, k_m, h_b, kk, z, ci_max):
    """
    A simplified RNA pulse model.

    This function models gene expression and metabolization of the internal
    concentration of a substance. The gene expression is controlled by a arctan
    step function based on the internal concentration (Ci) and switches on the
    gene's expression. The gene then translates a Protein, which metabolizes
    the internal concentration proportional to its expression level. The concept
    of protein must be understood not as a single Protein but as a collection
    of detoxification measures, which reduce the internal concentration of the
    compound and keep it at a reasonable level.

    Changes w.r.t. to RNA 4 model
    -------------------------------
    - The model does not evolve the survival probability S over time any more
      This is done more efficiently in the post processing, by simply taking the
      exponent


    Parameters
    ----------
    t : float
        Timestep at which the model is evaluated.

    X : tuple
        A tuple containing three elements:
        - Ce : float
            The external concentration.
        - Ci : float
            The internal concentration.
        - R : float
            The gene expression level.
        - P : float
            The protein level.
        - H : float
            The cumulative hazard.

    r_0 : float
        Initial value of the gene expression level.

    k_i : float
```

```
    Internal consumption rate constant.

k_m : float
    Metabolization rate constant.

r_rt : float
    Maximum gene expression rate constant. Termed k_rt in the paper

r_rd : float
    Gene degradation rate constant. Termed k_rd in the paper

z_ci : float
    The threshold for gene expression.

v_rt : float, optional
    The slope parameter for the inverse tangent step function. This
    parameter regulates the responsiveness of the gene expression induction.

k_p : float
    Dominant protein translation rate konstant.

Returns
--------
dCe_dt : float
    The rate of change of external concentration.

dCi_dt : float
    The rate of change of internal concentration.

dR_dt : float
    The rate of change of gene expression level.

dR_dt : float
    The rate of change of protein level.

dH_dt : float
    The hazard rate h(t) = b * max(D, 0) + h_b
"""
Ce, Ci, R, P, H = X

# active = 0.5 + (1 / jnp.pi) * jnp.arctan(v_rt * (Ci / ci_max - z_ci))
active = 1 / (1 + jnp.exp(- v_rt * (Ci/ci_max - z_ci)))

dCe_dt = 0.0
dCi_dt = Ce * k_i - Ci * P * k_m
dR_dt = r_rt * active - (R - r_0) * r_rd
dP_dt = k_p * ((R - r_0) - P)

dH_dt = kk * jnp.maximum(R - z, jnp.array([0.0], dtype=float)) + h_b

return dCe_dt, dCi_dt, dR_dt, dP_dt, dH_dt
```

##### S4.1.2 Solver post processing

```
def survival(results, t, interpolation):
    results["survival"] = jnp.exp(-results["H"])
    return results
```

##### S4.1.3 Probability model

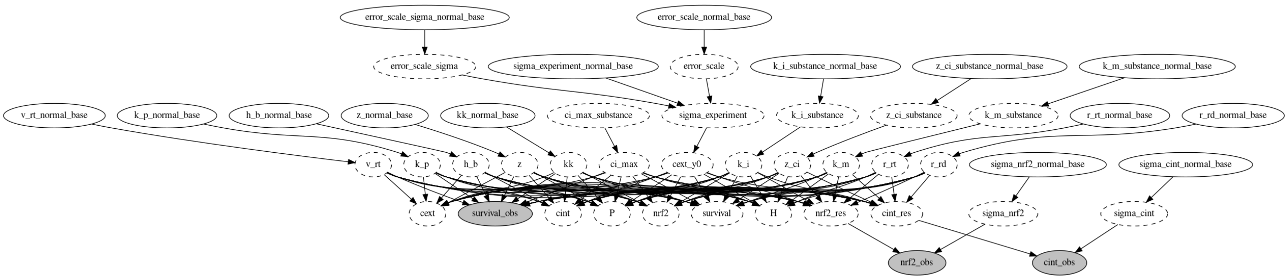


**Figure S13.** Directed acyclic graph (DAG) of the probability model.

#### S4.2 Report: Parameters ✓

### S4.2.1 $x_{in}$

No model input

### S4.2.2 $y_0$

| id | cext | cint | nrf2 | P | H |
|---|---|---|---|---|---|
| 101_0 | 2.34 | 0 | 1 | 0 | 0 |
| 101_1 | 2.34 | 0 | 1 | 0 | 0 |
| 106_0 | 5.16 | 0 | 1 | 0 | 0 |
| 106_1 | 5.16 | 0 | 1 | 0 | 0 |
| 112_0 | 11.72 | 0 | 1 | 0 | 0 |
| 112_1 | 11.72 | 0 | 1 | 0 | 0 |
| 118_0 | 18.14 | 0 | 1 | 0 | 0 |
| 118_1 | 18.14 | 0 | 1 | 0 | 0 |
| 124_0 | 29.44 | 0 | 1 | 0 | 0 |
| 124_1 | 29.44 | 0 | 1 | 0 | 0 |
| 184_0 | 2.12727 | 0 | 1 | 0 | 0 |
| 185_0 | 8.5091 | 0 | 1 | 0 | 0 |
| 186_0 | 10.6364 | 0 | 1 | 0 | 0 |
| 187_0 | 12.7636 | 0 | 1 | 0 | 0 |
| 188_0 | 14.8909 | 0 | 1 | 0 | 0 |
| 189_0 | 17.0182 | 0 | 1 | 0 | 0 |
| 190_0 | 25.5273 | 0 | 1 | 0 | 0 |
| 191_0 | 34.0364 | 0 | 1 | 0 | 0 |
| 192_0 | 45.7364 | 0 | 1 | 0 | 0 |
| 193_0 | 5.31819 | 0 | 1 | 0 | 0 |
| 194_0 | 6.38182 | 0 | 1 | 0 | 0 |
| 195_0 | 7.78583 | 0 | 1 | 0 | 0 |
| 196_0 | 9.31746 | 0 | 1 | 0 | 0 |
| 197_0 | 11.232 | 0 | 1 | 0 | 0 |
| 198_0 | 13.4869 | 0 | 1 | 0 | 0 |

| id | cext | cint | nrf2 | P | H |
|---|---|---|---|---|---|
| 199_0 | 15.7418 | 0 | 1 | 0 | 0 |
| 200_0 | 19.3582 | 0 | 1 | 0 | 0 |
| 201_0 | 23.2724 | 0 | 1 | 0 | 0 |
| 202_0 | 27.9098 | 0 | 1 | 0 | 0 |
| 203_0 | 33.5259 | 0 | 1 | 0 | 0 |
| 204_0 | 40.2055 | 0 | 1 | 0 | 0 |
| 205_0 | 48.2466 | 0 | 1 | 0 | 0 |
| 206_0 | 57.9044 | 0 | 1 | 0 | 0 |
| 207_0 | 69.4768 | 0 | 1 | 0 | 0 |
| 208_0 | 83.3892 | 0 | 1 | 0 | 0 |
| 209_0 | 2.08473 | 0 | 1 | 0 | 0 |
| 210_0 | 3.19091 | 0 | 1 | 0 | 0 |
| 211_0 | 4.7651 | 0 | 1 | 0 | 0 |
| 212_0 | 7.19019 | 0 | 1 | 0 | 0 |
| 213_0 | 8.5091 | 0 | 1 | 0 | 0 |
| 214_0 | 10.764 | 0 | 1 | 0 | 0 |
| 215_0 | 12.7636 | 0 | 1 | 0 | 0 |
| 216_0 | 14.8909 | 0 | 1 | 0 | 0 |
| 217_0 | 16.1247 | 0 | 1 | 0 | 0 |
| 218_0 | 24.1658 | 0 | 1 | 0 | 0 |
| 219_0 | 36.2913 | 0 | 1 | 0 | 0 |
| 220_0 | 54.4582 | 0 | 1 | 0 | 0 |
| 221_0 | 81.6874 | 0 | 1 | 0 | 0 |
| 222_0 | 8.01201 | 0 | 1 | 0 | 0 |
| 223_0 | 10.4156 | 0 | 1 | 0 | 0 |
| 224_0 | 13.5403 | 0 | 1 | 0 | 0 |
| 225_0 | 17.6024 | 0 | 1 | 0 | 0 |
| 226_0 | 22.8831 | 0 | 1 | 0 | 0 |
| 227_0 | 29.748 | 0 | 1 | 0 | 0 |
| 228_0 | 38.6724 | 0 | 1 | 0 | 0 |
| 229_0 | 50.2742 | 0 | 1 | 0 | 0 |
| 230_0 | 65.3564 | 0 | 1 | 0 | 0 |
| 231_0 | 84.9634 | 0 | 1 | 0 | 0 |
| 232_0 | 91.2601 | 0 | 1 | 0 | 0 |
| 42_0 | 28.098 | 0 | 1 | 0 | 0 |
| 44_0 | 29.462 | 0 | 1 | 0 | 0 |
| 44_1 | 29.462 | 0 | 1 | 0 | 0 |
| 44_2 | 29.462 | 0 | 1 | 0 | 0 |
| 44_3 | 29.462 | 0 | 1 | 0 | 0 |
| 44_4 | 29.462 | 0 | 1 | 0 | 0 |
| 44_5 | 29.462 | 0 | 1 | 0 | 0 |
| 44_6 | 29.462 | 0 | 1 | 0 | 0 |
| 44_7 | 29.462 | 0 | 1 | 0 | 0 |
| 51_0 | 20.5062 | 0 | 1 | 0 | 0 |
| 51_1 | 20.5062 | 0 | 1 | 0 | 0 |
| 51_2 | 20.5062 | 0 | 1 | 0 | 0 |
| 52_0 | 19.9914 | 0 | 1 | 0 | 0 |
| 52_1 | 19.9914 | 0 | 1 | 0 | 0 |
| 52_2 | 19.9914 | 0 | 1 | 0 | 0 |
| 53_0 | 19.9914 | 0 | 1 | 0 | 0 |

| id | cext | cint | nrf2 | P | H |
|---:|---:|---:|---:|---:|---:|
| 53_1 | 19.9914 | 0 | 1 | 0 | 0 |
| 69_0 | 19.9914 | 0 | 1 | 0 | 0 |
| 70_0 | 19.9914 | 0 | 1 | 0 | 0 |
| 70_1 | 19.9914 | 0 | 1 | 0 | 0 |
| 70_2 | 19.9914 | 0 | 1 | 0 | 0 |
| 74_0 | 19.9914 | 0 | 1 | 0 | 0 |
| 74_1 | 19.9914 | 0 | 1 | 0 | 0 |
| 74_2 | 19.9914 | 0 | 1 | 0 | 0 |
| 75_0 | 20.5062 | 0 | 1 | 0 | 0 |
| 8_0 | 20 | 0 | 1 | 0 | 0 |
| 10_0 | 7.36 | 0 | 1 | 0 | 0 |
| 126_0 | 0.878067 | 0 | 1 | 0 | 0 |
| 127_0 | 1.75613 | 0 | 1 | 0 | 0 |
| 128_0 | 3.51227 | 0 | 1 | 0 | 0 |
| 129_0 | 7.02453 | 0 | 1 | 0 | 0 |
| 130_0 | 14.0491 | 0 | 1 | 0 | 0 |
| 131_0 | 28.0981 | 0 | 1 | 0 | 0 |
| 132_0 | 56.1963 | 0 | 1 | 0 | 0 |
| 133_0 | 112.393 | 0 | 1 | 0 | 0 |
| 134_0 | 224.785 | 0 | 1 | 0 | 0 |
| 136_0 | 449.57 | 0 | 1 | 0 | 0 |
| 138_0 | 3.29335 | 0 | 1 | 0 | 0 |
| 139_0 | 4.28135 | 0 | 1 | 0 | 0 |
| 140_0 | 5.56576 | 0 | 1 | 0 | 0 |
| 141_0 | 7.23549 | 0 | 1 | 0 | 0 |
| 142_0 | 9.40613 | 0 | 1 | 0 | 0 |
| 143_0 | 12.228 | 0 | 1 | 0 | 0 |
| 144_0 | 15.8964 | 0 | 1 | 0 | 0 |
| 145_0 | 20.6653 | 0 | 1 | 0 | 0 |
| 146_0 | 26.8649 | 0 | 1 | 0 | 0 |
| 147_0 | 34.9243 | 0 | 1 | 0 | 0 |
| 148_0 | 7.99998 | 0 | 1 | 0 | 0 |
| 149_0 | 9.59998 | 0 | 1 | 0 | 0 |
| 150_0 | 11.52 | 0 | 1 | 0 | 0 |
| 151_0 | 13.824 | 0 | 1 | 0 | 0 |
| 152_0 | 16.5888 | 0 | 1 | 0 | 0 |
| 153_0 | 19.9065 | 0 | 1 | 0 | 0 |
| 154_0 | 23.8878 | 0 | 1 | 0 | 0 |
| 155_0 | 28.6654 | 0 | 1 | 0 | 0 |
| 156_0 | 34.3985 | 0 | 1 | 0 | 0 |
| 157_0 | 41.2781 | 0 | 1 | 0 | 0 |
| 158_0 | 3.66326 | 0 | 1 | 0 | 0 |
| 159_0 | 4.57907 | 0 | 1 | 0 | 0 |
| 160_0 | 5.72384 | 0 | 1 | 0 | 0 |
| 161_0 | 7.1548 | 0 | 1 | 0 | 0 |
| 162_0 | 8.94349 | 0 | 1 | 0 | 0 |
| 163_0 | 11.1794 | 0 | 1 | 0 | 0 |
| 164_0 | 13.9742 | 0 | 1 | 0 | 0 |
| 165_0 | 17.4678 | 0 | 1 | 0 | 0 |
| 166_0 | 21.8347 | 0 | 1 | 0 | 0 |

| id | cext | cint | nrf2 | P | H |
|---|---|---|---|---|---|
| 167_0 | 27.2934 | 0 | 1 | 0 | 0 |
| 168_0 | 34.1167 | 0 | 1 | 0 | 0 |
| 169_0 | 42.6459 | 0 | 1 | 0 | 0 |
| 170_0 | 53.3074 | 0 | 1 | 0 | 0 |
| 171_0 | 3.14673 | 0 | 1 | 0 | 0 |
| 172_0 | 3.77607 | 0 | 1 | 0 | 0 |
| 173_0 | 4.53128 | 0 | 1 | 0 | 0 |
| 174_0 | 5.43754 | 0 | 1 | 0 | 0 |
| 175_0 | 6.52505 | 0 | 1 | 0 | 0 |
| 176_0 | 7.83006 | 0 | 1 | 0 | 0 |
| 177_0 | 9.39607 | 0 | 1 | 0 | 0 |
| 178_0 | 11.2753 | 0 | 1 | 0 | 0 |
| 179_0 | 13.5303 | 0 | 1 | 0 | 0 |
| 180_0 | 16.2364 | 0 | 1 | 0 | 0 |
| 181_0 | 19.4837 | 0 | 1 | 0 | 0 |
| 182_0 | 23.3804 | 0 | 1 | 0 | 0 |
| 183_0 | 28.0565 | 0 | 1 | 0 | 0 |
| 38_0 | 7.364 | 0 | 1 | 0 | 0 |
| 43_0 | 6.60108 | 0 | 1 | 0 | 0 |
| 43_1 | 6.60108 | 0 | 1 | 0 | 0 |
| 43_2 | 6.60108 | 0 | 1 | 0 | 0 |
| 43_3 | 6.60108 | 0 | 1 | 0 | 0 |
| 54_0 | 5.02939 | 0 | 1 | 0 | 0 |
| 54_1 | 5.02939 | 0 | 1 | 0 | 0 |
| 55_0 | 6.60108 | 0 | 1 | 0 | 0 |
| 55_1 | 6.60108 | 0 | 1 | 0 | 0 |
| 55_2 | 6.60108 | 0 | 1 | 0 | 0 |
| 56_0 | 6.60108 | 0 | 1 | 0 | 0 |
| 56_1 | 6.60108 | 0 | 1 | 0 | 0 |
| 56_2 | 6.60108 | 0 | 1 | 0 | 0 |
| 56_3 | 6.60108 | 0 | 1 | 0 | 0 |
| 56_4 | 6.60108 | 0 | 1 | 0 | 0 |
| 56_5 | 6.60108 | 0 | 1 | 0 | 0 |
| 57_0 | 7.22975 | 0 | 1 | 0 | 0 |
| 57_1 | 7.22975 | 0 | 1 | 0 | 0 |
| 57_2 | 7.22975 | 0 | 1 | 0 | 0 |
| 57_3 | 7.22975 | 0 | 1 | 0 | 0 |
| 57_4 | 7.22975 | 0 | 1 | 0 | 0 |
| 57_5 | 7.22975 | 0 | 1 | 0 | 0 |
| 62_0 | 6.60108 | 0 | 1 | 0 | 0 |
| 62_1 | 6.60108 | 0 | 1 | 0 | 0 |
| 67_0 | 6.60108 | 0 | 1 | 0 | 0 |
| 67_1 | 6.60108 | 0 | 1 | 0 | 0 |
| 67_2 | 6.60108 | 0 | 1 | 0 | 0 |
| 67_3 | 6.60108 | 0 | 1 | 0 | 0 |
| 67_4 | 6.60108 | 0 | 1 | 0 | 0 |
| 67_5 | 6.60108 | 0 | 1 | 0 | 0 |
| 68_0 | 7.22975 | 0 | 1 | 0 | 0 |
| 68_1 | 7.22975 | 0 | 1 | 0 | 0 |
| 68_2 | 7.22975 | 0 | 1 | 0 | 0 |

| id | cext | cint | nrf2 | P | H |
|---|---|---|---|---|---|
| 68_3 | 7.22975 | 0 | 1 | 0 | 0 |
| 71_0 | 5.02939 | 0 | 1 | 0 | 0 |
| 72_0 | 6.60108 | 0 | 1 | 0 | 0 |
| 72_1 | 6.60108 | 0 | 1 | 0 | 0 |
| 72_2 | 6.60108 | 0 | 1 | 0 | 0 |
| 72_3 | 6.60108 | 0 | 1 | 0 | 0 |
| 72_4 | 6.60108 | 0 | 1 | 0 | 0 |
| 72_5 | 6.60108 | 0 | 1 | 0 | 0 |
| 73_0 | 7.22975 | 0 | 1 | 0 | 0 |
| 73_1 | 7.22975 | 0 | 1 | 0 | 0 |
| 73_2 | 7.22975 | 0 | 1 | 0 | 0 |
| 78_0 | 6.60108 | 0 | 1 | 0 | 0 |
| 78_1 | 6.60108 | 0 | 1 | 0 | 0 |
| 80_0 | 7.364 | 0 | 1 | 0 | 0 |
| 82_0 | 5.1 | 0 | 1 | 0 | 0 |
| 82_1 | 5.1 | 0 | 1 | 0 | 0 |
| 84_0 | 5.77 | 0 | 1 | 0 | 0 |
| 84_1 | 5.77 | 0 | 1 | 0 | 0 |
| 86_0 | 6.52 | 0 | 1 | 0 | 0 |
| 86_1 | 6.52 | 0 | 1 | 0 | 0 |
| 88_0 | 6.93 | 0 | 1 | 0 | 0 |
| 88_1 | 6.93 | 0 | 1 | 0 | 0 |
| 88_2 | 6.93 | 0 | 1 | 0 | 0 |
| 90_0 | 7.36 | 0 | 1 | 0 | 0 |
| 90_1 | 7.36 | 0 | 1 | 0 | 0 |
| 91_0 | 5.1 | 0 | 1 | 0 | 0 |
| 91_1 | 5.1 | 0 | 1 | 0 | 0 |
| 92_0 | 5.77 | 0 | 1 | 0 | 0 |
| 92_1 | 5.77 | 0 | 1 | 0 | 0 |
| 93_0 | 6.52 | 0 | 1 | 0 | 0 |
| 93_1 | 6.52 | 0 | 1 | 0 | 0 |
| 94_0 | 6.93 | 0 | 1 | 0 | 0 |
| 94_1 | 6.93 | 0 | 1 | 0 | 0 |
| 95_0 | 7.36 | 0 | 1 | 0 | 0 |
| 102_0 | 134.58 | 0 | 1 | 0 | 0 |
| 102_1 | 134.58 | 0 | 1 | 0 | 0 |
| 107_0 | 177.57 | 0 | 1 | 0 | 0 |
| 113_0 | 234.29 | 0 | 1 | 0 | 0 |
| 113_1 | 234.29 | 0 | 1 | 0 | 0 |
| 119_0 | 269.13 | 0 | 1 | 0 | 0 |
| 119_1 | 269.13 | 0 | 1 | 0 | 0 |
| 125_0 | 309.14 | 0 | 1 | 0 | 0 |
| 125_1 | 309.14 | 0 | 1 | 0 | 0 |
| 233_0 | 10.5849 | 0 | 1 | 0 | 0 |
| 234_0 | 21.1698 | 0 | 1 | 0 | 0 |
| 235_0 | 42.3396 | 0 | 1 | 0 | 0 |
| 236_0 | 84.6793 | 0 | 1 | 0 | 0 |
| 237_0 | 169.359 | 0 | 1 | 0 | 0 |
| 238_0 | 338.717 | 0 | 1 | 0 | 0 |
| 239_0 | 677.434 | 0 | 1 | 0 | 0 |

| id | cext | cint | nrf2 | P | H |
|---|---|---|---|---|---|
| 240_0 | 1354.87 | 0 | 1 | 0 | 0 |
| 241_0 | 281.571 | 0 | 1 | 0 | 0 |
| 242_0 | 337.886 | 0 | 1 | 0 | 0 |
| 243_0 | 405.463 | 0 | 1 | 0 | 0 |
| 244_0 | 486.556 | 0 | 1 | 0 | 0 |
| 245_0 | 583.867 | 0 | 1 | 0 | 0 |
| 247_0 | 700.64 | 0 | 1 | 0 | 0 |
| 249_0 | 840.768 | 0 | 1 | 0 | 0 |
| 251_0 | 1008.92 | 0 | 1 | 0 | 0 |
| 253_0 | 1210.71 | 0 | 1 | 0 | 0 |
| 254_0 | 1452.85 | 0 | 1 | 0 | 0 |
| 255_0 | 137.422 | 0 | 1 | 0 | 0 |
| 256_0 | 164.906 | 0 | 1 | 0 | 0 |
| 257_0 | 197.888 | 0 | 1 | 0 | 0 |
| 258_0 | 237.465 | 0 | 1 | 0 | 0 |
| 259_0 | 284.958 | 0 | 1 | 0 | 0 |
| 260_0 | 341.95 | 0 | 1 | 0 | 0 |
| 261_0 | 410.34 | 0 | 1 | 0 | 0 |
| 262_0 | 492.408 | 0 | 1 | 0 | 0 |
| 27_0 | 238.256 | 0 | 1 | 0 | 0 |
| 28_0 | 238.256 | 0 | 1 | 0 | 0 |
| 33_0 | 238.256 | 0 | 1 | 0 | 0 |
| 40_0 | 306.652 | 0 | 1 | 0 | 0 |
| 48_0 | 134.792 | 0 | 1 | 0 | 0 |
| 48_1 | 134.792 | 0 | 1 | 0 | 0 |
| 48_2 | 134.792 | 0 | 1 | 0 | 0 |
| 48_3 | 134.792 | 0 | 1 | 0 | 0 |
| 48_4 | 134.792 | 0 | 1 | 0 | 0 |
| 48_5 | 134.792 | 0 | 1 | 0 | 0 |
| 48_6 | 134.792 | 0 | 1 | 0 | 0 |
| 49_0 | 309.229 | 0 | 1 | 0 | 0 |
| 49_1 | 309.229 | 0 | 1 | 0 | 0 |
| 49_2 | 309.229 | 0 | 1 | 0 | 0 |
| 49_3 | 309.229 | 0 | 1 | 0 | 0 |
| 49_4 | 309.229 | 0 | 1 | 0 | 0 |
| 49_5 | 309.229 | 0 | 1 | 0 | 0 |
| 49_6 | 309.229 | 0 | 1 | 0 | 0 |
| 58_0 | 134.792 | 0 | 1 | 0 | 0 |
| 58_1 | 134.792 | 0 | 1 | 0 | 0 |
| 58_2 | 134.792 | 0 | 1 | 0 | 0 |
| 59_0 | 309.229 | 0 | 1 | 0 | 0 |
| 59_1 | 309.229 | 0 | 1 | 0 | 0 |
| 59_2 | 309.229 | 0 | 1 | 0 | 0 |
| 5_0 | 348.834 | 0 | 1 | 0 | 0 |
| 60_0 | 134.792 | 0 | 1 | 0 | 0 |
| 60_1 | 134.792 | 0 | 1 | 0 | 0 |
| 61_0 | 309.229 | 0 | 1 | 0 | 0 |
| 61_1 | 309.229 | 0 | 1 | 0 | 0 |
| 63_0 | 134.792 | 0 | 1 | 0 | 0 |
| 63_1 | 134.792 | 0 | 1 | 0 | 0 |

| id | cext | cint | nrf2 | P | H |
|---|---|---|---|---|---|
| 63_2 | 134.792 | 0 | 1 | 0 | 0 |
| 63_3 | 134.792 | 0 | 1 | 0 | 0 |
| 64_0 | 309.229 | 0 | 1 | 0 | 0 |
| 64_1 | 309.229 | 0 | 1 | 0 | 0 |
| 64_2 | 309.229 | 0 | 1 | 0 | 0 |
| 64_3 | 309.229 | 0 | 1 | 0 | 0 |
| 65_0 | 134.792 | 0 | 1 | 0 | 0 |
| 65_1 | 134.792 | 0 | 1 | 0 | 0 |
| 65_2 | 134.792 | 0 | 1 | 0 | 0 |
| 65_3 | 134.792 | 0 | 1 | 0 | 0 |
| 65_4 | 134.792 | 0 | 1 | 0 | 0 |
| 65_5 | 134.792 | 0 | 1 | 0 | 0 |
| 66_0 | 309.229 | 0 | 1 | 0 | 0 |
| 66_1 | 309.229 | 0 | 1 | 0 | 0 |
| 66_2 | 309.229 | 0 | 1 | 0 | 0 |
| 66_3 | 309.229 | 0 | 1 | 0 | 0 |
| 66_4 | 309.229 | 0 | 1 | 0 | 0 |
| 66_5 | 309.229 | 0 | 1 | 0 | 0 |
| 6_0 | 348.834 | 0 | 1 | 0 | 0 |
| 76_0 | 134.792 | 0 | 1 | 0 | 0 |
| 76_1 | 134.792 | 0 | 1 | 0 | 0 |
| 76_10 | 134.792 | 0 | 1 | 0 | 0 |
| 76_2 | 134.792 | 0 | 1 | 0 | 0 |
| 76_3 | 134.792 | 0 | 1 | 0 | 0 |
| 76_4 | 134.792 | 0 | 1 | 0 | 0 |
| 76_5 | 134.792 | 0 | 1 | 0 | 0 |
| 76_6 | 134.792 | 0 | 1 | 0 | 0 |
| 76_7 | 134.792 | 0 | 1 | 0 | 0 |
| 76_8 | 134.792 | 0 | 1 | 0 | 0 |
| 76_9 | 134.792 | 0 | 1 | 0 | 0 |
| 77_0 | 309.229 | 0 | 1 | 0 | 0 |
| 77_1 | 309.229 | 0 | 1 | 0 | 0 |
| 77_2 | 309.229 | 0 | 1 | 0 | 0 |
| 77_3 | 309.229 | 0 | 1 | 0 | 0 |
| 77_4 | 309.229 | 0 | 1 | 0 | 0 |
| 77_5 | 309.229 | 0 | 1 | 0 | 0 |
| 77_6 | 309.229 | 0 | 1 | 0 | 0 |
| 77_7 | 309.229 | 0 | 1 | 0 | 0 |
| 77_8 | 309.229 | 0 | 1 | 0 | 0 |

##### S4.2.3 Free parameters

- error_scale ~ halfnorm(scale=1,dims=())
- error_scale_sigma ~ halfnorm(scale=1,dims=())
- sigma_experiment ~ lognorm(scale=error_scale,s=error_scale_sigma,dims=('experiment_id',))
- k_i_substance ~ lognorm(scale=[1.0,1.0,1.0],s=2,dims=('substance',))
- z_ci_substance ~ lognorm(scale=[0.5,0.5,0.5],s=2,dims=('substance',))
- k_m_substance ~ lognorm(scale=[0.05,0.05,0.05],s=2,dims=('substance',))
- r_rt ~ lognorm(scale=1.0,s=2,dims=())
- r_rd ~ lognorm(scale=0.5,s=2,dims=())

- v_rt ~ lognorm(scale=1.0,s=2,dims=())
- k_p ~ lognorm(scale=0.02,s=2,dims=())
- h_b ~ lognorm(scale=1e-08,s=2,dims=())
- z ~ lognorm(scale=1.0,s=2,dims=())
- kk ~ lognorm(scale=0.02,s=2,dims=())
- sigma_nrf2 ~ halfnorm(scale=5.0,dims=())
- sigma_cint ~ halfnorm(scale=5.0,dims=())

###### S4.2.4 Fixed parameters

- cext_y0 = cext_y0 * sigma_experiment[experiment_id_index], dims=('id',)
- k_i = k_i_substance[substance_index], dims=('id',)
- z_ci = z_ci_substance[substance_index], dims=('id',)
- k_m = k_m_substance[substance_index], dims=('id',)
- ci_max_substance = [1757.0, 168.1, 6364.8], dims=('substance',)
- ci_max = ci_max_substance[substance_index], dims=('id',)
- r_0 = 1.0, dims=()

##### S4.3 Report: Table parameter estimates ✓

Excluding parameters: ['sigma_experiment'] for meaningful visualization

| | ('index', ' ') | ('diuron', 'mean ± std') | ('diclofenac', 'mean ± std') | ('naproxen', 'mean ± std') |
|---|---|---|---|---|
| 0 | ci_max_substance | 1757.0 ± 0.0 | 168.1 ± 0.0 | 6364.8 ± 0.0 |
| 1 | error_scale | 0.577 ± 0.085 | 0.577 ± 0.085 | 0.577 ± 0.085 |
| 2 | error_scale_sigma | 1.079 ± 0.084 | 1.079 ± 0.084 | 1.079 ± 0.084 |
| 3 | h_b | 0.0 ± 0.0 | 0.0 ± 0.0 | 0.0 ± 0.0 |
| 4 | k_i_substance | 6.882 ± 1.156 | 0.741 ± 0.085 | 0.568 ± 0.055 |
| 5 | k_m_substance | 0.88 ± 0.157 | 0.055 ± 0.005 | 0.023 ± 0.002 |
| 6 | k_p | 0.038 ± 0.006 | 0.038 ± 0.006 | 0.038 ± 0.006 |
| 7 | kk | 0.078 ± 0.008 | 0.078 ± 0.008 | 0.078 ± 0.008 |
| 8 | r_rd | 0.495 ± 0.044 | 0.495 ± 0.044 | 0.495 ± 0.044 |
| 9 | r_rt | 3.742 ± 0.398 | 3.742 ± 0.398 | 3.742 ± 0.398 |
| 10 | sigma_cint | 0.588 ± 0.02 | 0.588 ± 0.02 | 0.588 ± 0.02 |
| 11 | sigma_nrf2 | 0.293 ± 0.009 | 0.293 ± 0.009 | 0.293 ± 0.009 |
| 12 | v_rt | 10.805 ± 0.398 | 10.805 ± 0.398 | 10.805 ± 0.398 |
| 13 | z | 1.471 ± 0.032 | 1.471 ± 0.032 | 1.471 ± 0.032 |
| 14 | z_ci_substance | 0.412 ± 0.027 | 0.371 ± 0.016 | 0.435 ± 0.016 |

Report 'table_parameter_estimates' was successfully generated and saved in '../hierarchical_molecular_tktd/results

##### S4.4 Report: Goodness of fit ✓

| | cint | nrf2 | survival | model |
|---|---|---|---|---|
| NRMSE | 0.178896 | 0.166608 | 0.356855 | nan |
| NRMSE (95%-hdi[lower]) | 0.109386 | 0.160374 | 0.337507 | nan |
| NRMSE (95%-hdi[upper]) | 0.255927 | 0.172948 | 0.376839 | nan |
| Log-Likelihood | -829.362 | -30.6933 | -290.642 | -1150.7 |
| Log-Likelihood (95%-hdi[lower]) | -855.101 | -33.956 | -317.592 | -1186.47 |
| Log-Likelihood (95%-hdi[upper]) | -804.335 | -27.7943 | -275.123 | -1118.86 |

| | cint | nrf2 | survival | model |
|---|---|---|---|---|
| n (data) | 913 | 169 | 392 | 1474 |
| k (parameters) | nan | nan | nan | 62 |
| BIC | nan | nan | nan | 2753.73 |
| BIC (95%-hdi[lower]) | nan | nan | nan | 2690.05 |
| BIC (95%-hdi[upper]) | nan | nan | nan | 2825.28 |

Report ‘goodness_of_fit’ was successfully generated and saved in ‘../hierarchical_molecular_tktd/results/hierarchica

#### S4.5 Report: Diagnostics ✓

Excluding parameters: [‘sigma_experiment’] for meaningful visualization

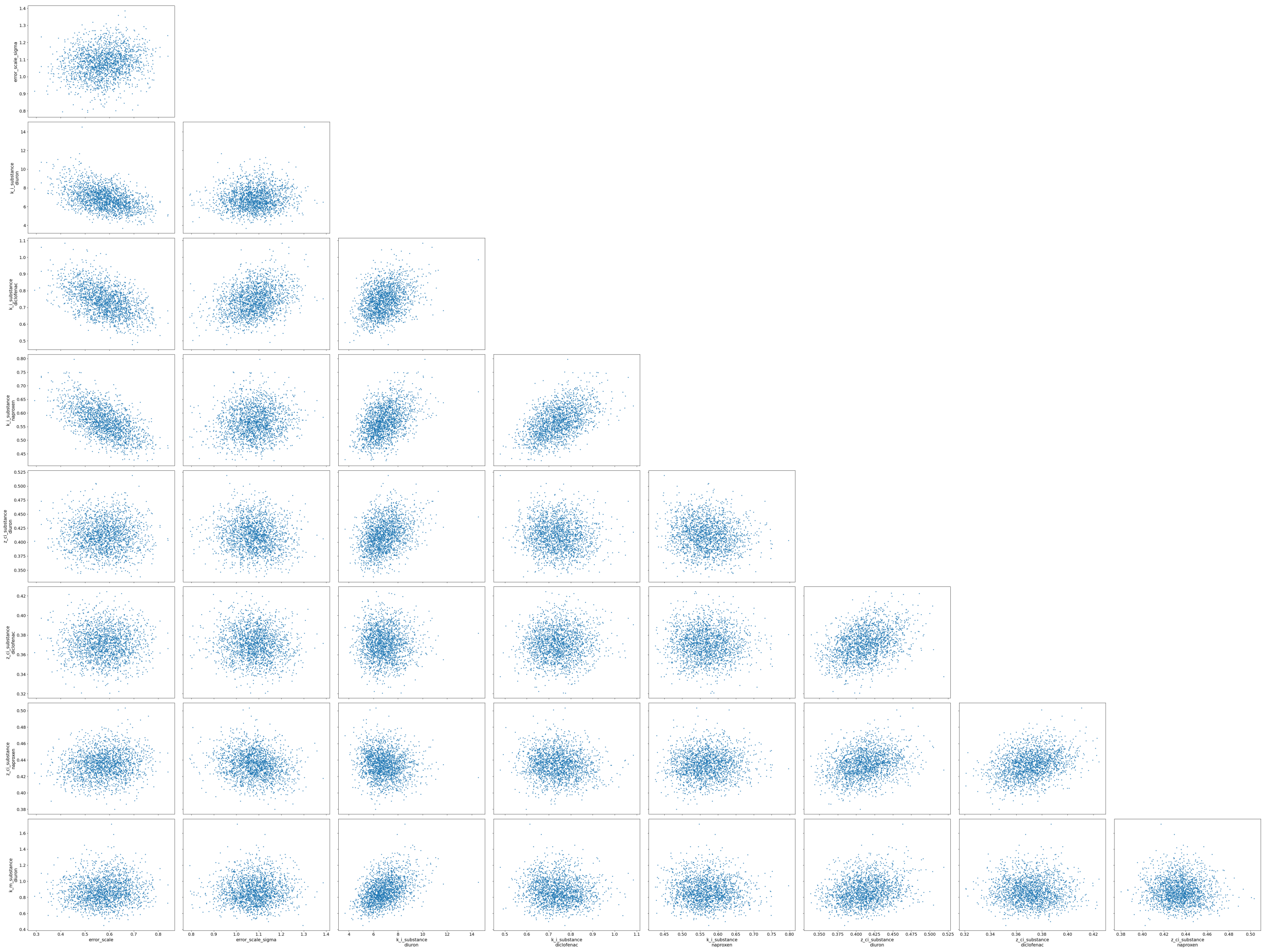

**Figure S14.** Paired parameter estimates

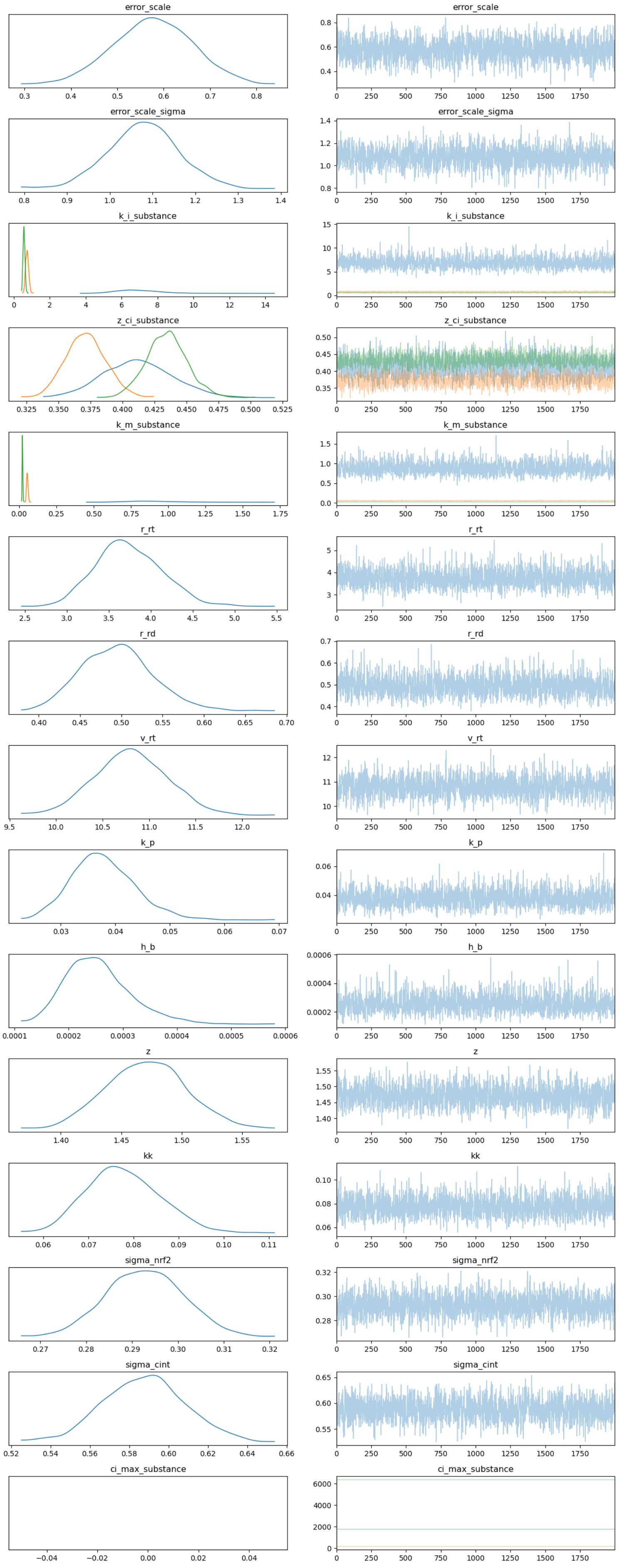


**Figure S15.** Psuedo trace, generated for draws from the optimized SVI distribution

Report ‘diagnostics’ was successfully generated and saved in ‘(’../hierarchical_molecular_tktd/results/hierarchical_
‘../hierarchical_molecular_tktd/results/hierarchical_cext_nested_sigma_hyperprior_rna_pulse_5_substance_independen

#### S4.6 Report: Visualizations ✓

#### S4.7 Report: Visualizations ✓

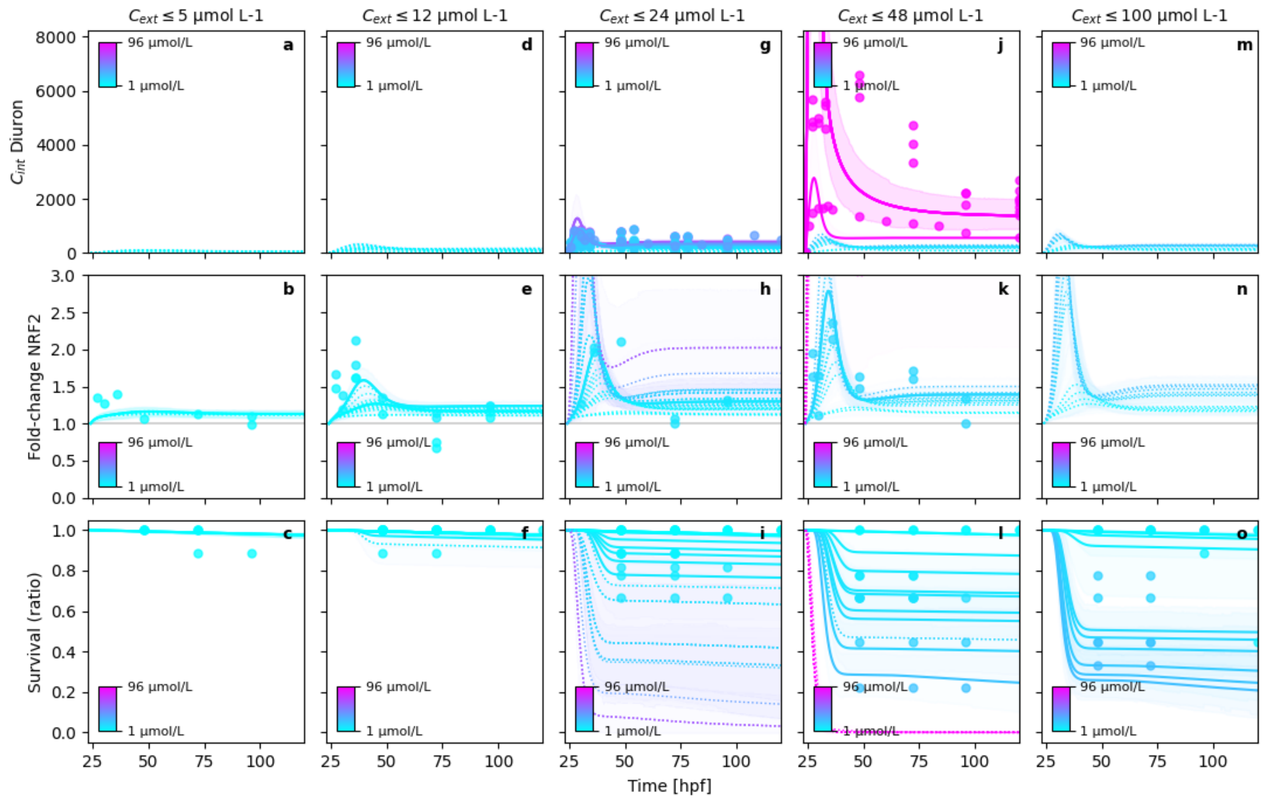


**Figure S16.** Posterior model fits

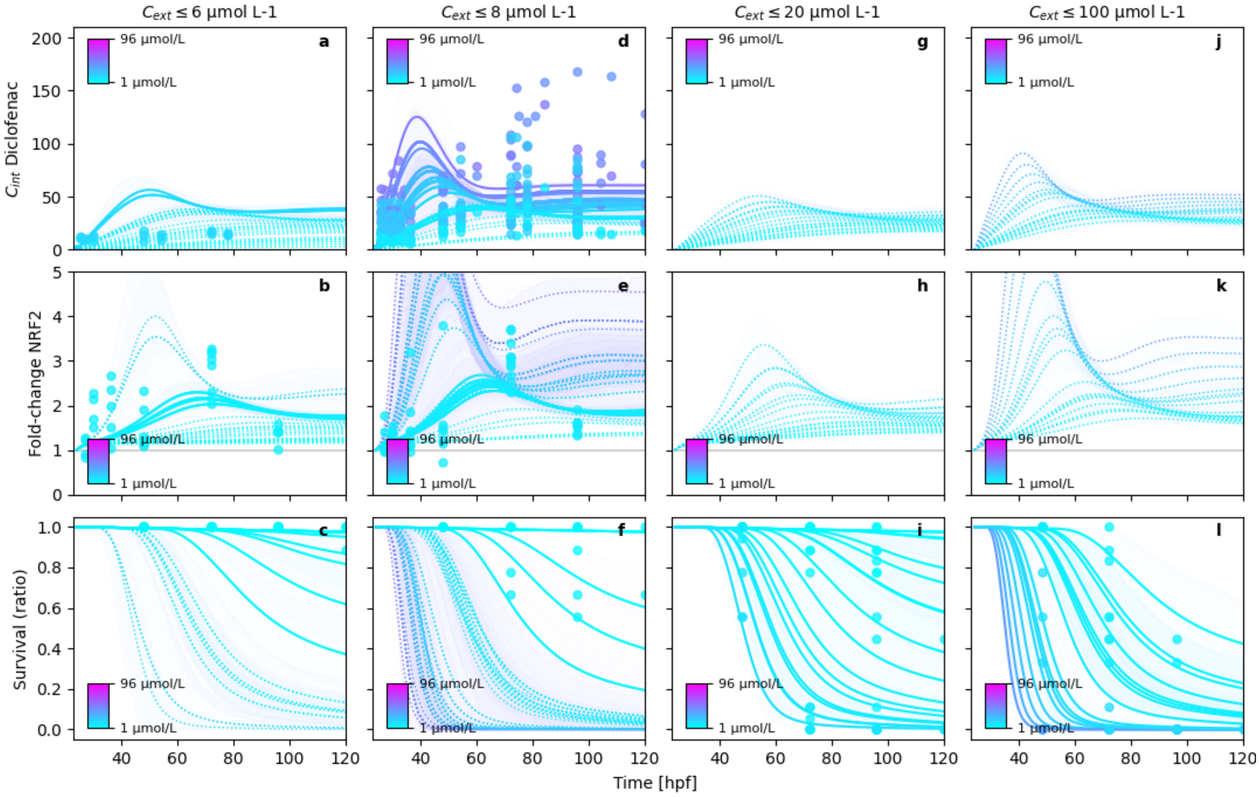


**Figure S17.** Posterior model fits

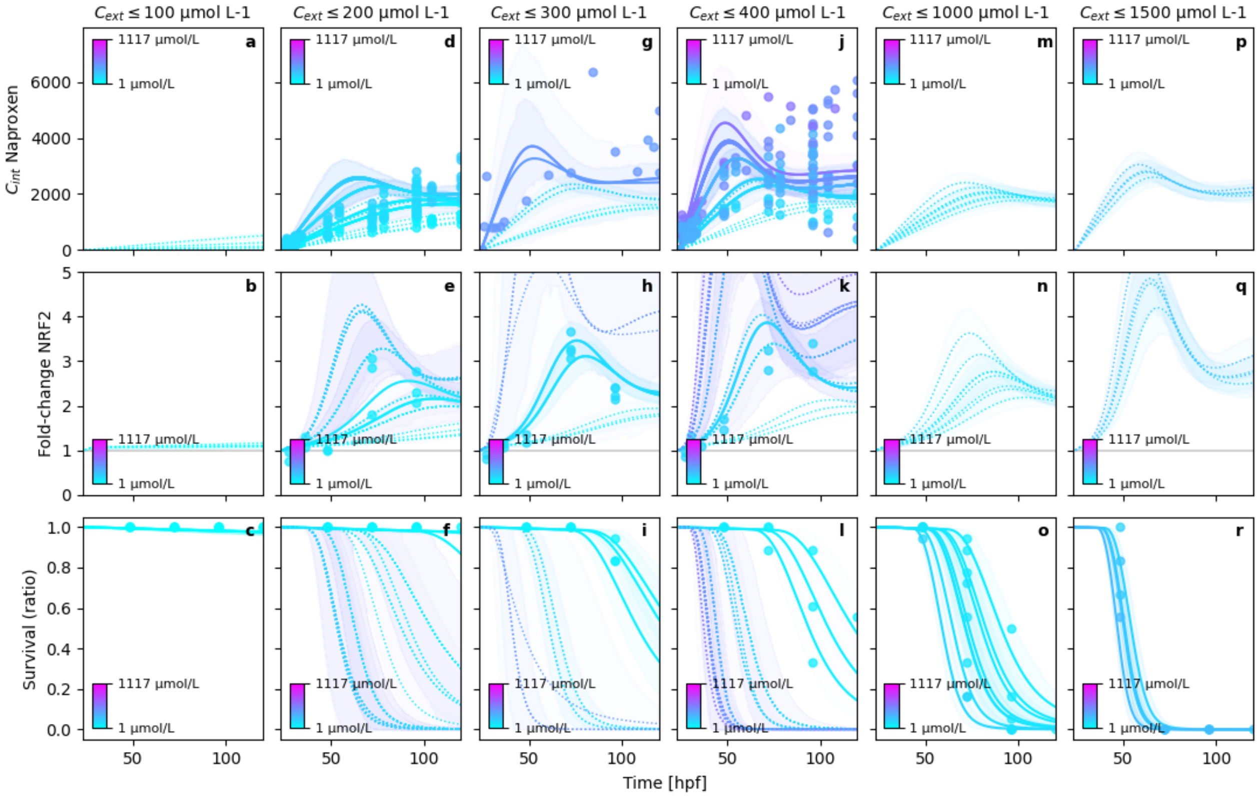


**Figure S18.** Posterior model fits

Report 'visualizations' was successfully generated and saved in '['../hierarchical_molecular_tktd/results/hierarchical_c
'../hierarchical_molecular_tktd/results/hierarchical_cext_nested_sigma_hyperprior_rna_pulse_5_substance_independen

'../hierarchical_molecular_tktd/results/hierarchical_cext_nested_sigma_hyperprior_rna_pulse_5_substance_independen

##### S4.7.1 $y_0$ estimation of external concentrations

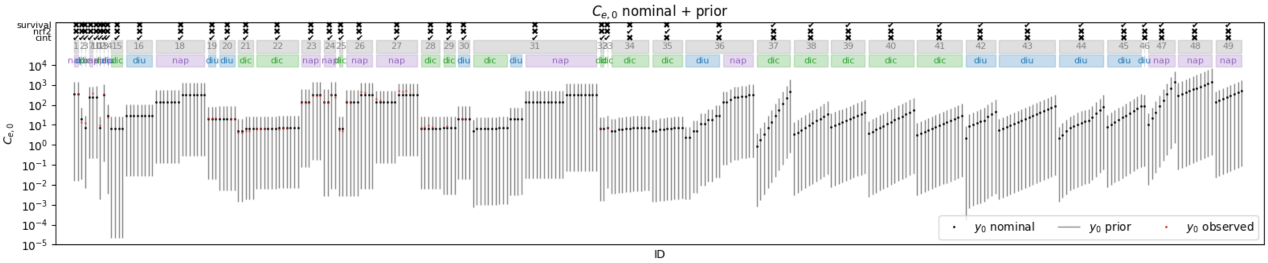


**Figure S19.** Prior $C_{ext,0}$ estimates and nominal concentrations

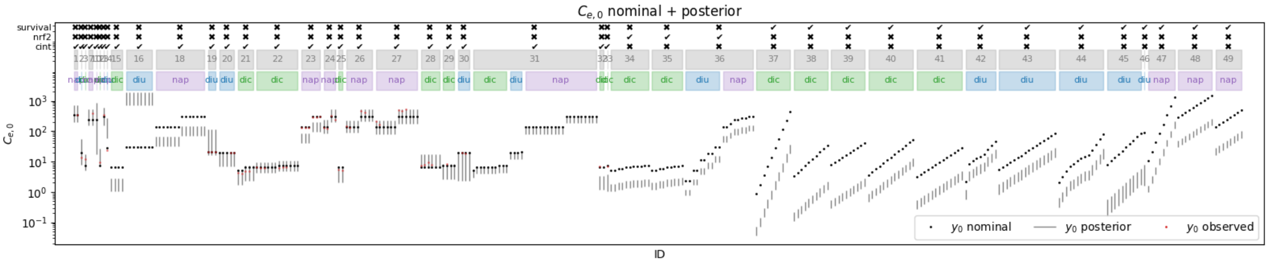


**Figure S20.** Posterior $C_{ext,0}$ estimates and nominal concentrations

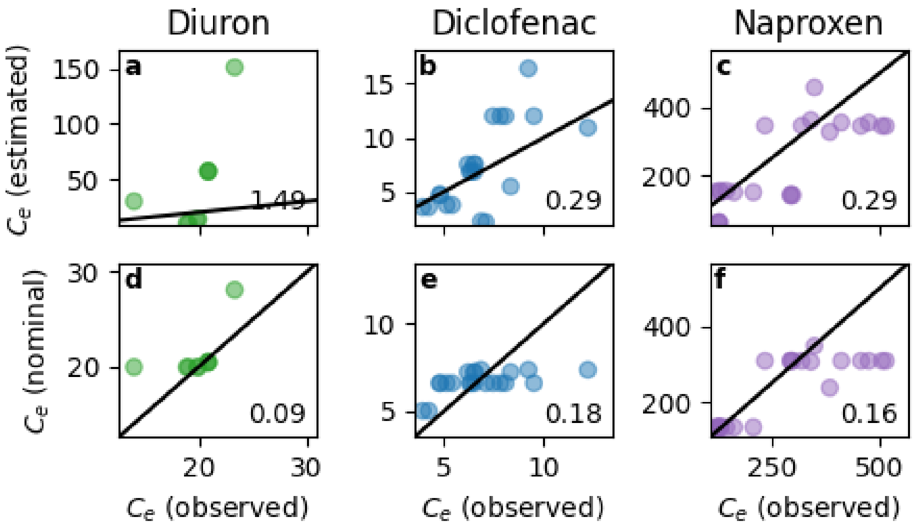


**Figure S21.** $C_{ext,0}$ comparison between nominal, measured and estimated concentrations

Report 'visualizations' was successfully generated and saved in '('../hierarchical_molecular_tktd/results/hierarchical_c
'../hierarchical_molecular_tktd/results/hierarchical_cext_nested_sigma_hyperprior_rna_pulse_5_substance_independen
'../hierarchical_molecular_tktd/results/hierarchical_cext_nested_sigma_hyperprior_rna_pulse_5_substance_independen

#### S4.8 Report: Model inadequacy metrics ✓

##### S4.8.1 Residuals

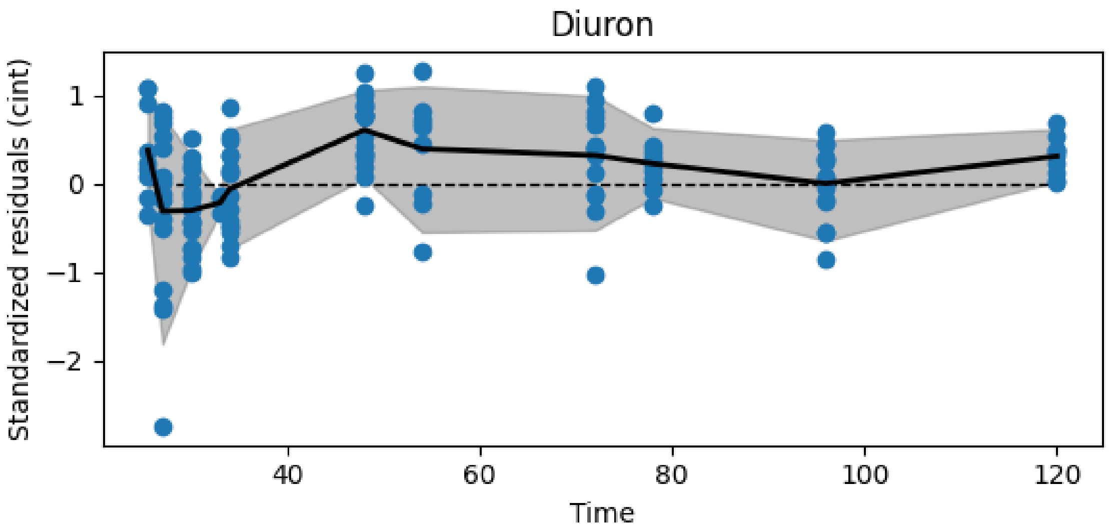


**Figure S22.** Resiudal cint dynamics of diuron

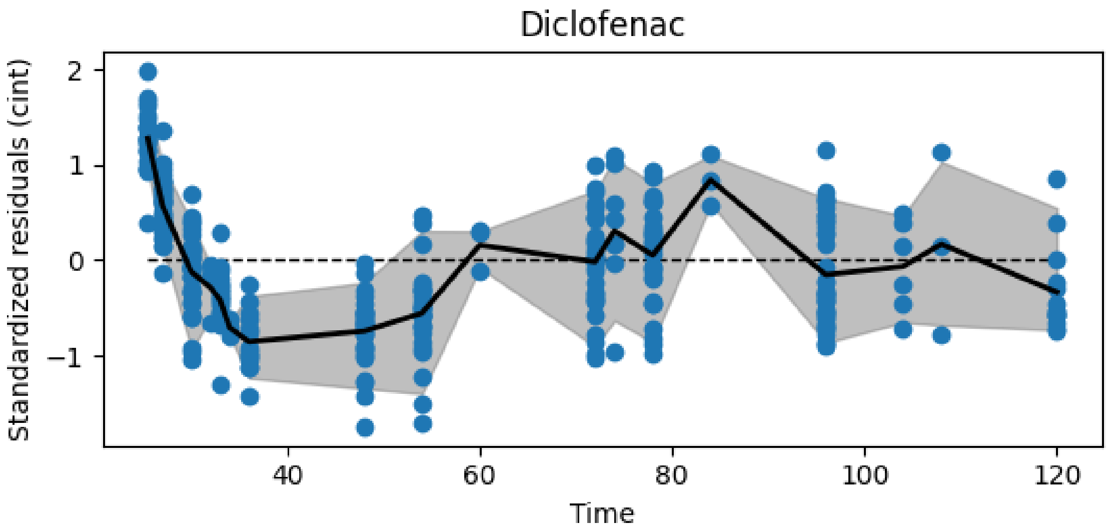


**Figure S23.** Resiudal cint dynamics of diclofenac

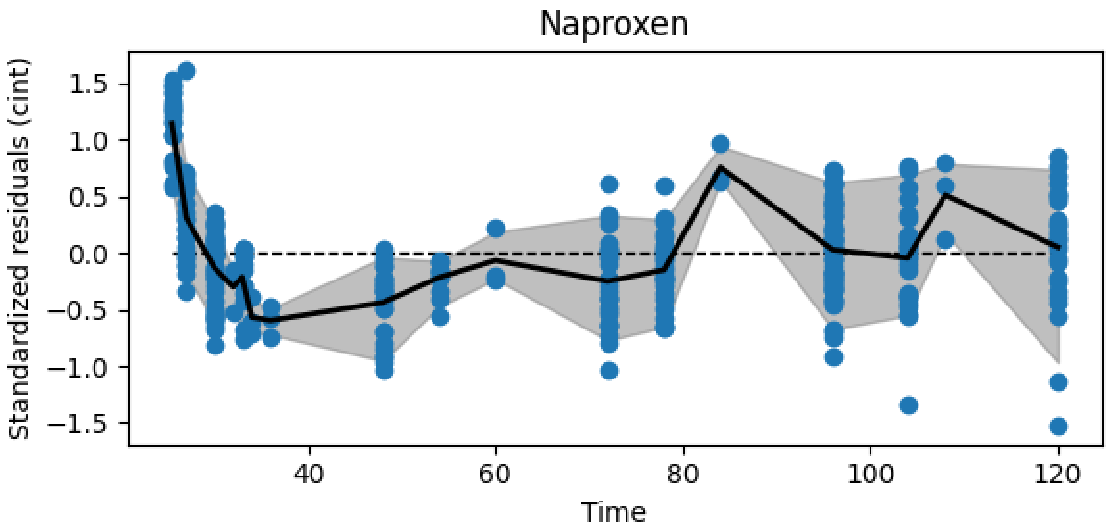


**Figure S24.** Resiudal cint dynamics of naproxen

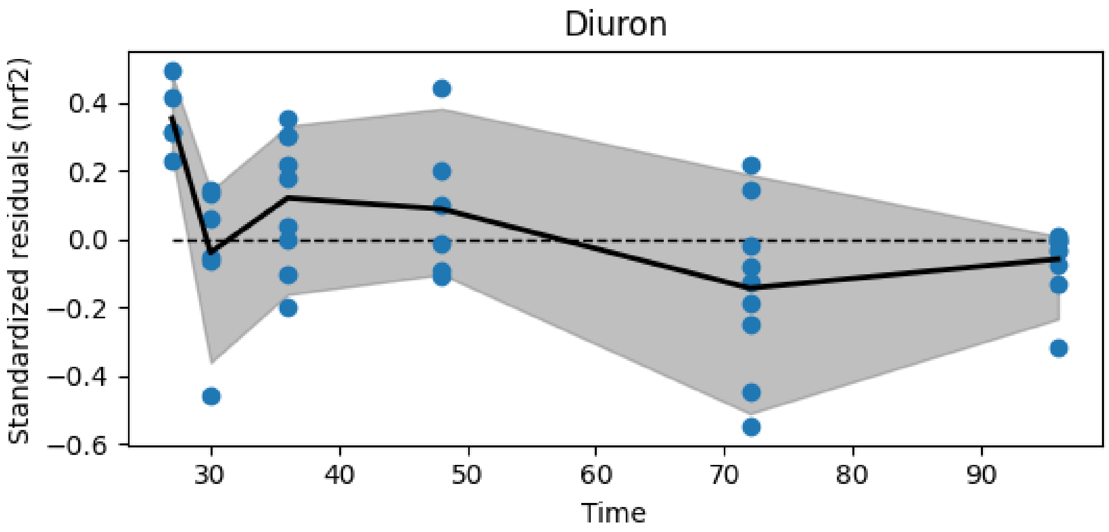


**Figure S25.** Resiudal nrf2 dynamics of diuron

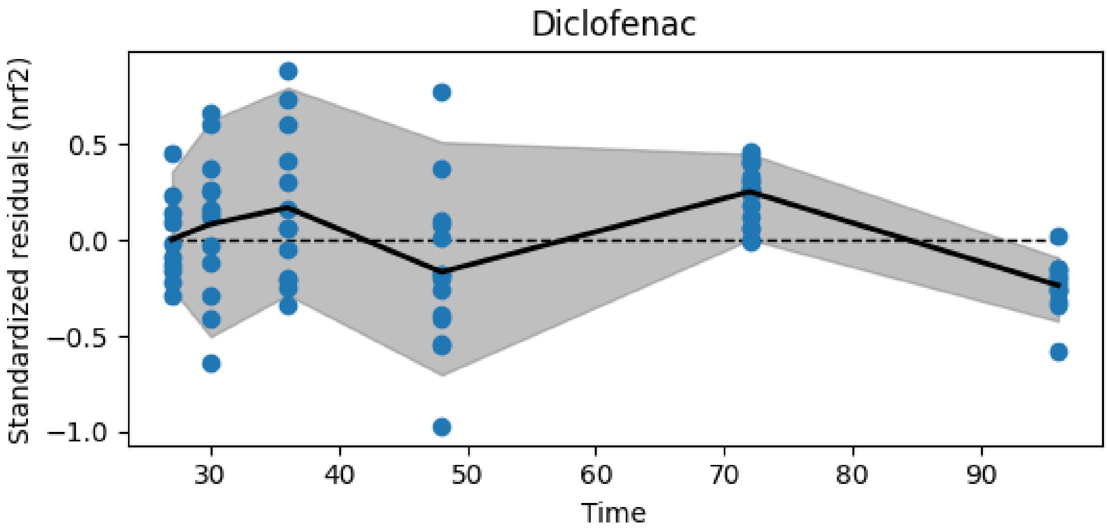


**Figure S26.** Resiudal nrf2 dynamics of diclofenac

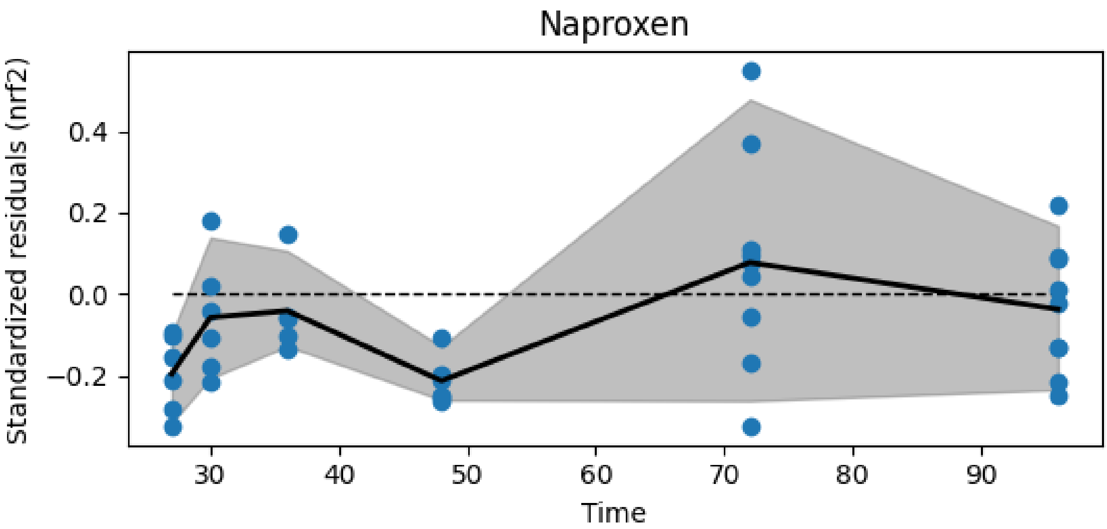


**Figure S27.** Resiudal nrf2 dynamics of naproxen

##### S4.8.2 Model inadequacy

The different metrics are measures for the model inadequacy. The comparison to `metric_value_if_normal_dist` is a simulation of normally distributed residuals that have the same data structure in terms of dimensionality (id x time) and missing values. If the `metric_value` falls within that interval, the model can be assumed as not inadequate.

- **autocorrelation**: Measures the correlation of the residuals with themselves with a lag of 1. High absolute autocorrelation means, the variable is not normally distributed. Ideal would be values close to 0.
- **deviation log-prob**: Uses a t-test to estimate the probability of the replicates at a time t being different from zero. The result is the summed log-probability. Low (negative) log probs indicate high probability for deviation.
- **significant deviations**: Uses a t-test to estimate the probability of the replicates at a time t being different from zero. The result is the number of significant deviations (for an alpha level of 0.05). High number of deviations indicate an inadequate model

- **local/global variance**: This metric calculates the local variance as a rolling variance of always 3 direct neighboring residuals. The local variances are then averaged and divided by the global averages of all residuals. The basis for the calculation is the residuals averaged by id. Values close to 1 indicate an adequate model
- **replicate/global variance**: This metric calculates the replicate variance at time t, averages it and divides the number by the global variance. The local variances are then averaged and divided by the global averages of all residuals. The basis for the calculation is the residuals averaged by id. Values close to 1 indicate an adequate model

| | metric | data_variable | index | metric_value | metric_value_if_normal_dist |
|---|---|---|---|---|---|
| 0 | autocorrelation | cint | diclofenac | 0.587241 | -0.06[-0.41,0.2] |
| 1 | autocorrelation | cint | diuron | 0.383679 | -0.1[-0.52,0.34] |
| 2 | autocorrelation | cint | naproxen | 0.459032 | -0.05[-0.37,0.26] |
| 3 | autocorrelation | nrf2 | diclofenac | -0.894373 | -0.23[-0.83,0.53] |
| 4 | autocorrelation | nrf2 | diuron | -0.117739 | -0.22[-0.84,0.6] |
| 5 | autocorrelation | nrf2 | naproxen | -0.407495 | -0.18[-0.7,0.38] |
| 6 | deviation log-prob | cint | diclofenac | -155.759 | -18.39[-26.83,-12.0] |
| 7 | deviation log-prob | cint | diuron | -41.8672 | -11.07[-16.71,-6.47] |
| 8 | deviation log-prob | cint | naproxen | -99.1327 | -16.27[-22.65,-10.75] |
| 9 | deviation log-prob | nrf2 | diclofenac | -29.8393 | -6.01[-10.22,-2.13] |
| 10 | deviation log-prob | nrf2 | diuron | -14.4741 | -6.1[-10.67,-2.72] |
| 11 | deviation log-prob | nrf2 | naproxen | -17.0252 | -6.39[-11.54,-2.95] |
| 12 | local/global variance | cint | diclofenac | 0.298292 | 0.73[0.56,0.9] |
| 13 | local/global variance | cint | diuron | 0.462983 | 0.76[0.46,0.99] |
| 14 | local/global variance | cint | naproxen | 0.359188 | 0.74[0.56,0.91] |
| 15 | local/global variance | nrf2 | diclofenac | 0.865023 | 0.82[0.45,1.2] |
| 16 | local/global variance | nrf2 | diuron | 0.517547 | 0.79[0.42,1.12] |
| 17 | local/global variance | nrf2 | naproxen | 0.986805 | 0.82[0.43,1.17] |
| 18 | replicate/global variance | cint | diclofenac | 0.373891 | 0.88[0.75,1.0] |
| 19 | replicate/global variance | cint | diuron | 0.661849 | 0.9[0.82,1.02] |
| 20 | replicate/global variance | cint | naproxen | 0.360554 | 0.86[0.76,0.99] |
| 21 | replicate/global variance | nrf2 | diclofenac | 0.746767 | 0.94[0.86,1.02] |
| 22 | replicate/global variance | nrf2 | diuron | 0.579886 | 0.88[0.73,0.99] |
| 23 | replicate/global variance | nrf2 | naproxen | 0.592473 | 0.86[0.7,1.0] |
| 24 | significant deviations | cint | diclofenac | 9 | 0.99[0.0,3.0] |
| 25 | significant deviations | cint | diuron | 5 | 0.59[0.0,2.0] |
| 26 | significant deviations | cint | naproxen | 11 | 0.66[0.0,2.0] |
| 27 | significant deviations | nrf2 | diclofenac | 2 | 0.37[0.0,1.0] |
| 28 | significant deviations | nrf2 | diuron | 1 | 0.26[0.0,1.0] |
| 29 | significant deviations | nrf2 | naproxen | 2 | 0.36[0.0,2.0] |

Report 'model_inadequacy_metrics' was successfully generated and saved in '../hierarchical_molecular_tktd/results

### S5 Report(case_study=hierarchical_molecular_tktd, scenario=hierarchical_ce

- Using `hierarchical_molecular_tktd==0.1.6`
- Using `pymob==0.5.6a3`
- Using backend: `NumpyroBackend`
- Using settings: `../hierarchical_molecular_tktd/scenarios/hierarchical_cext_nested_sigma_hyperp`

#### S5.1 Report: Model ✓

##### S5.1.1 Model

```
def tktd_rna_5(t, X, r_0, k_i, r_rt, r_rd, z_ci, v_rt, k_p, k_m, h_b, kk, z, ci_max):
    """
    A simplified RNA pulse model.

    This function models gene expression and metabolization of the internal
    concentration of a substance. The gene expression is controlled by a arctan
    step function based on the internal concentration (Ci) and switches on the
    gene's expression. The gene then translates a Protein, which metabolizes
    the internal concentration proportional to its expression level. The concept
    of protein must be understood not as a single Protein but as a collection
    of detoxification measures, which reduce the internal concentration of the
    compound and keep it at a reasonable level.

    Changes w.r.t. to RNA 4 model
    -------------------------------
    - The model does not evolve the survival probability S over time any more
      This is done more efficiently in the post processing, by simply taking the
      exponent


    Parameters
    ----------
    t : float
        Timestep at which the model is evaluated.

    X : tuple
        A tuple containing three elements:
        - Ce : float
            The external concentration.
        - Ci : float
            The internal concentration.
        - R : float
            The gene expression level.
        - P : float
            The protein level.
        - H : float
            The cumulative hazard.

    r_0 : float
        Initial value of the gene expression level.

    k_i : float
```

```
        Internal consumption rate constant.

    k_m : float
        Metabolization rate constant.

    r_rt : float
        Maximum gene expression rate constant. Termed k_rt in the paper

    r_rd : float
        Gene degradation rate constant. Termed k_rd in the paper

    z_ci : float
        The threshold for gene expression.

    v_rt : float, optional
        The slope parameter for the inverse tangent step function. This
        parameter regulates the responsiveness of the gene expression induction.

    k_p : float
        Dominant protein translation rate konstant.

    Returns
    -------
    dCe_dt : float
        The rate of change of external concentration.

    dCi_dt : float
        The rate of change of internal concentration.

    dR_dt : float
        The rate of change of gene expression level.

    dR_dt : float
        The rate of change of protein level.

    dH_dt : float
        The hazard rate h(t) = b * max(D, 0) + h_b
    """
    Ce, Ci, R, P, H = X

    # active = 0.5 + (1 / jnp.pi) * jnp.arctan(v_rt * (Ci / ci_max - z_ci))
    active = 1 / (1 + jnp.exp(- v_rt * (Ci/ci_max - z_ci)))

    dCe_dt = 0.0
    dCi_dt = Ce * k_i - Ci * P * k_m
    dR_dt = r_rt * active - (R - r_0) * r_rd
    dP_dt = k_p * ((R - r_0) - P)

    dH_dt = kk * jnp.maximum(R - z, jnp.array([0.0], dtype=float)) + h_b

    return dCe_dt, dCi_dt, dR_dt, dP_dt, dH_dt
```

##### S5.1.2 Solver post processing

```python
def survival(results, t, interpolation):
    results["survival"] = jnp.exp(-results["H"])
    return results
```

##### S5.1.3 Probability model

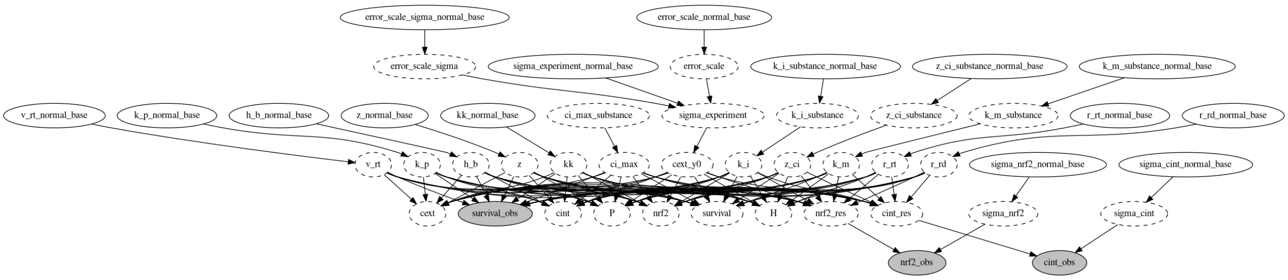


**Figure S28.** Directed acyclic graph (DAG) of the probability model.

#### S5.2 Report: Parameters ✓

### S5.2.1 $x_{in}$

No model input

### S5.2.2 $y_0$

| id | cext | cint | nrf2 | P | H |
|---|---:|---:|---:|---:|---:|
| 101_0 | 2.34 | 0 | 1 | 0 | 0 |
| 101_1 | 2.34 | 0 | 1 | 0 | 0 |
| 106_0 | 5.16 | 0 | 1 | 0 | 0 |
| 106_1 | 5.16 | 0 | 1 | 0 | 0 |
| 112_0 | 11.72 | 0 | 1 | 0 | 0 |
| 112_1 | 11.72 | 0 | 1 | 0 | 0 |
| 118_0 | 18.14 | 0 | 1 | 0 | 0 |
| 118_1 | 18.14 | 0 | 1 | 0 | 0 |
| 124_0 | 29.44 | 0 | 1 | 0 | 0 |
| 124_1 | 29.44 | 0 | 1 | 0 | 0 |
| 184_0 | 2.12727 | 0 | 1 | 0 | 0 |
| 185_0 | 8.5091 | 0 | 1 | 0 | 0 |
| 186_0 | 10.6364 | 0 | 1 | 0 | 0 |
| 187_0 | 12.7636 | 0 | 1 | 0 | 0 |
| 188_0 | 14.8909 | 0 | 1 | 0 | 0 |
| 189_0 | 17.0182 | 0 | 1 | 0 | 0 |
| 190_0 | 25.5273 | 0 | 1 | 0 | 0 |
| 191_0 | 34.0364 | 0 | 1 | 0 | 0 |
| 192_0 | 45.7364 | 0 | 1 | 0 | 0 |
| 193_0 | 5.31819 | 0 | 1 | 0 | 0 |
| 194_0 | 6.38182 | 0 | 1 | 0 | 0 |
| 195_0 | 7.78583 | 0 | 1 | 0 | 0 |
| 196_0 | 9.31746 | 0 | 1 | 0 | 0 |
| 197_0 | 11.232 | 0 | 1 | 0 | 0 |
| 198_0 | 13.4869 | 0 | 1 | 0 | 0 |

| id | cext | cint | nrf2 | P | H |
|---|---|---|---|---|---|
| 199_0 | 15.7418 | 0 | 1 | 0 | 0 |
| 200_0 | 19.3582 | 0 | 1 | 0 | 0 |
| 201_0 | 23.2724 | 0 | 1 | 0 | 0 |
| 202_0 | 27.9098 | 0 | 1 | 0 | 0 |
| 203_0 | 33.5259 | 0 | 1 | 0 | 0 |
| 204_0 | 40.2055 | 0 | 1 | 0 | 0 |
| 205_0 | 48.2466 | 0 | 1 | 0 | 0 |
| 206_0 | 57.9044 | 0 | 1 | 0 | 0 |
| 207_0 | 69.4768 | 0 | 1 | 0 | 0 |
| 208_0 | 83.3892 | 0 | 1 | 0 | 0 |
| 209_0 | 2.08473 | 0 | 1 | 0 | 0 |
| 210_0 | 3.19091 | 0 | 1 | 0 | 0 |
| 211_0 | 4.7651 | 0 | 1 | 0 | 0 |
| 212_0 | 7.19019 | 0 | 1 | 0 | 0 |
| 213_0 | 8.5091 | 0 | 1 | 0 | 0 |
| 214_0 | 10.764 | 0 | 1 | 0 | 0 |
| 215_0 | 12.7636 | 0 | 1 | 0 | 0 |
| 216_0 | 14.8909 | 0 | 1 | 0 | 0 |
| 217_0 | 16.1247 | 0 | 1 | 0 | 0 |
| 218_0 | 24.1658 | 0 | 1 | 0 | 0 |
| 219_0 | 36.2913 | 0 | 1 | 0 | 0 |
| 220_0 | 54.4582 | 0 | 1 | 0 | 0 |
| 221_0 | 81.6874 | 0 | 1 | 0 | 0 |
| 222_0 | 8.01201 | 0 | 1 | 0 | 0 |
| 223_0 | 10.4156 | 0 | 1 | 0 | 0 |
| 224_0 | 13.5403 | 0 | 1 | 0 | 0 |
| 225_0 | 17.6024 | 0 | 1 | 0 | 0 |
| 226_0 | 22.8831 | 0 | 1 | 0 | 0 |
| 227_0 | 29.748 | 0 | 1 | 0 | 0 |
| 228_0 | 38.6724 | 0 | 1 | 0 | 0 |
| 229_0 | 50.2742 | 0 | 1 | 0 | 0 |
| 230_0 | 65.3564 | 0 | 1 | 0 | 0 |
| 231_0 | 84.9634 | 0 | 1 | 0 | 0 |
| 232_0 | 91.2601 | 0 | 1 | 0 | 0 |
| 42_0 | 28.098 | 0 | 1 | 0 | 0 |
| 44_0 | 29.462 | 0 | 1 | 0 | 0 |
| 44_1 | 29.462 | 0 | 1 | 0 | 0 |
| 44_2 | 29.462 | 0 | 1 | 0 | 0 |
| 44_3 | 29.462 | 0 | 1 | 0 | 0 |
| 44_4 | 29.462 | 0 | 1 | 0 | 0 |
| 44_5 | 29.462 | 0 | 1 | 0 | 0 |
| 44_6 | 29.462 | 0 | 1 | 0 | 0 |
| 44_7 | 29.462 | 0 | 1 | 0 | 0 |
| 51_0 | 20.5062 | 0 | 1 | 0 | 0 |
| 51_1 | 20.5062 | 0 | 1 | 0 | 0 |
| 51_2 | 20.5062 | 0 | 1 | 0 | 0 |
| 52_0 | 19.9914 | 0 | 1 | 0 | 0 |
| 52_1 | 19.9914 | 0 | 1 | 0 | 0 |
| 52_2 | 19.9914 | 0 | 1 | 0 | 0 |
| 53_0 | 19.9914 | 0 | 1 | 0 | 0 |

| id | cext | cint | nrf2 | P | H |
|---|---|---|---|---|---|
| 53_1 | 19.9914 | 0 | 1 | 0 | 0 |
| 69_0 | 19.9914 | 0 | 1 | 0 | 0 |
| 70_0 | 19.9914 | 0 | 1 | 0 | 0 |
| 70_1 | 19.9914 | 0 | 1 | 0 | 0 |
| 70_2 | 19.9914 | 0 | 1 | 0 | 0 |
| 74_0 | 19.9914 | 0 | 1 | 0 | 0 |
| 74_1 | 19.9914 | 0 | 1 | 0 | 0 |
| 74_2 | 19.9914 | 0 | 1 | 0 | 0 |
| 75_0 | 20.5062 | 0 | 1 | 0 | 0 |
| 8_0 | 20 | 0 | 1 | 0 | 0 |
| 10_0 | 7.36 | 0 | 1 | 0 | 0 |
| 126_0 | 0.878067 | 0 | 1 | 0 | 0 |
| 127_0 | 1.75613 | 0 | 1 | 0 | 0 |
| 128_0 | 3.51227 | 0 | 1 | 0 | 0 |
| 129_0 | 7.02453 | 0 | 1 | 0 | 0 |
| 130_0 | 14.0491 | 0 | 1 | 0 | 0 |
| 131_0 | 28.0981 | 0 | 1 | 0 | 0 |
| 132_0 | 56.1963 | 0 | 1 | 0 | 0 |
| 133_0 | 112.393 | 0 | 1 | 0 | 0 |
| 134_0 | 224.785 | 0 | 1 | 0 | 0 |
| 136_0 | 449.57 | 0 | 1 | 0 | 0 |
| 138_0 | 3.29335 | 0 | 1 | 0 | 0 |
| 139_0 | 4.28135 | 0 | 1 | 0 | 0 |
| 140_0 | 5.56576 | 0 | 1 | 0 | 0 |
| 141_0 | 7.23549 | 0 | 1 | 0 | 0 |
| 142_0 | 9.40613 | 0 | 1 | 0 | 0 |
| 143_0 | 12.228 | 0 | 1 | 0 | 0 |
| 144_0 | 15.8964 | 0 | 1 | 0 | 0 |
| 145_0 | 20.6653 | 0 | 1 | 0 | 0 |
| 146_0 | 26.8649 | 0 | 1 | 0 | 0 |
| 147_0 | 34.9243 | 0 | 1 | 0 | 0 |
| 148_0 | 7.99998 | 0 | 1 | 0 | 0 |
| 149_0 | 9.59998 | 0 | 1 | 0 | 0 |
| 150_0 | 11.52 | 0 | 1 | 0 | 0 |
| 151_0 | 13.824 | 0 | 1 | 0 | 0 |
| 152_0 | 16.5888 | 0 | 1 | 0 | 0 |
| 153_0 | 19.9065 | 0 | 1 | 0 | 0 |
| 154_0 | 23.8878 | 0 | 1 | 0 | 0 |
| 155_0 | 28.6654 | 0 | 1 | 0 | 0 |
| 156_0 | 34.3985 | 0 | 1 | 0 | 0 |
| 157_0 | 41.2781 | 0 | 1 | 0 | 0 |
| 158_0 | 3.66326 | 0 | 1 | 0 | 0 |
| 159_0 | 4.57907 | 0 | 1 | 0 | 0 |
| 160_0 | 5.72384 | 0 | 1 | 0 | 0 |
| 161_0 | 7.1548 | 0 | 1 | 0 | 0 |
| 162_0 | 8.94349 | 0 | 1 | 0 | 0 |
| 163_0 | 11.1794 | 0 | 1 | 0 | 0 |
| 164_0 | 13.9742 | 0 | 1 | 0 | 0 |
| 165_0 | 17.4678 | 0 | 1 | 0 | 0 |
| 166_0 | 21.8347 | 0 | 1 | 0 | 0 |

| id | cext | cint | nrf2 | P | H |
|---|---|---|---|---|---|
| 167_0 | 27.2934 | 0 | 1 | 0 | 0 |
| 168_0 | 34.1167 | 0 | 1 | 0 | 0 |
| 169_0 | 42.6459 | 0 | 1 | 0 | 0 |
| 170_0 | 53.3074 | 0 | 1 | 0 | 0 |
| 171_0 | 3.14673 | 0 | 1 | 0 | 0 |
| 172_0 | 3.77607 | 0 | 1 | 0 | 0 |
| 173_0 | 4.53128 | 0 | 1 | 0 | 0 |
| 174_0 | 5.43754 | 0 | 1 | 0 | 0 |
| 175_0 | 6.52505 | 0 | 1 | 0 | 0 |
| 176_0 | 7.83006 | 0 | 1 | 0 | 0 |
| 177_0 | 9.39607 | 0 | 1 | 0 | 0 |
| 178_0 | 11.2753 | 0 | 1 | 0 | 0 |
| 179_0 | 13.5303 | 0 | 1 | 0 | 0 |
| 180_0 | 16.2364 | 0 | 1 | 0 | 0 |
| 181_0 | 19.4837 | 0 | 1 | 0 | 0 |
| 182_0 | 23.3804 | 0 | 1 | 0 | 0 |
| 183_0 | 28.0565 | 0 | 1 | 0 | 0 |
| 38_0 | 7.364 | 0 | 1 | 0 | 0 |
| 43_0 | 6.60108 | 0 | 1 | 0 | 0 |
| 43_1 | 6.60108 | 0 | 1 | 0 | 0 |
| 43_2 | 6.60108 | 0 | 1 | 0 | 0 |
| 43_3 | 6.60108 | 0 | 1 | 0 | 0 |
| 54_0 | 5.02939 | 0 | 1 | 0 | 0 |
| 54_1 | 5.02939 | 0 | 1 | 0 | 0 |
| 55_0 | 6.60108 | 0 | 1 | 0 | 0 |
| 55_1 | 6.60108 | 0 | 1 | 0 | 0 |
| 55_2 | 6.60108 | 0 | 1 | 0 | 0 |
| 56_0 | 6.60108 | 0 | 1 | 0 | 0 |
| 56_1 | 6.60108 | 0 | 1 | 0 | 0 |
| 56_2 | 6.60108 | 0 | 1 | 0 | 0 |
| 56_3 | 6.60108 | 0 | 1 | 0 | 0 |
| 56_4 | 6.60108 | 0 | 1 | 0 | 0 |
| 56_5 | 6.60108 | 0 | 1 | 0 | 0 |
| 57_0 | 7.22975 | 0 | 1 | 0 | 0 |
| 57_1 | 7.22975 | 0 | 1 | 0 | 0 |
| 57_2 | 7.22975 | 0 | 1 | 0 | 0 |
| 57_3 | 7.22975 | 0 | 1 | 0 | 0 |
| 57_4 | 7.22975 | 0 | 1 | 0 | 0 |
| 57_5 | 7.22975 | 0 | 1 | 0 | 0 |
| 62_0 | 6.60108 | 0 | 1 | 0 | 0 |
| 62_1 | 6.60108 | 0 | 1 | 0 | 0 |
| 67_0 | 6.60108 | 0 | 1 | 0 | 0 |
| 67_1 | 6.60108 | 0 | 1 | 0 | 0 |
| 67_2 | 6.60108 | 0 | 1 | 0 | 0 |
| 67_3 | 6.60108 | 0 | 1 | 0 | 0 |
| 67_4 | 6.60108 | 0 | 1 | 0 | 0 |
| 67_5 | 6.60108 | 0 | 1 | 0 | 0 |
| 68_0 | 7.22975 | 0 | 1 | 0 | 0 |
| 68_1 | 7.22975 | 0 | 1 | 0 | 0 |
| 68_2 | 7.22975 | 0 | 1 | 0 | 0 |

| id | cext | cint | nrf2 | P | H |
|---|---|---|---|---|---|
| 68_3 | 7.22975 | 0 | 1 | 0 | 0 |
| 71_0 | 5.02939 | 0 | 1 | 0 | 0 |
| 72_0 | 6.60108 | 0 | 1 | 0 | 0 |
| 72_1 | 6.60108 | 0 | 1 | 0 | 0 |
| 72_2 | 6.60108 | 0 | 1 | 0 | 0 |
| 72_3 | 6.60108 | 0 | 1 | 0 | 0 |
| 72_4 | 6.60108 | 0 | 1 | 0 | 0 |
| 72_5 | 6.60108 | 0 | 1 | 0 | 0 |
| 73_0 | 7.22975 | 0 | 1 | 0 | 0 |
| 73_1 | 7.22975 | 0 | 1 | 0 | 0 |
| 73_2 | 7.22975 | 0 | 1 | 0 | 0 |
| 78_0 | 6.60108 | 0 | 1 | 0 | 0 |
| 78_1 | 6.60108 | 0 | 1 | 0 | 0 |
| 80_0 | 7.364 | 0 | 1 | 0 | 0 |
| 82_0 | 5.1 | 0 | 1 | 0 | 0 |
| 82_1 | 5.1 | 0 | 1 | 0 | 0 |
| 84_0 | 5.77 | 0 | 1 | 0 | 0 |
| 84_1 | 5.77 | 0 | 1 | 0 | 0 |
| 86_0 | 6.52 | 0 | 1 | 0 | 0 |
| 86_1 | 6.52 | 0 | 1 | 0 | 0 |
| 88_0 | 6.93 | 0 | 1 | 0 | 0 |
| 88_1 | 6.93 | 0 | 1 | 0 | 0 |
| 88_2 | 6.93 | 0 | 1 | 0 | 0 |
| 90_0 | 7.36 | 0 | 1 | 0 | 0 |
| 90_1 | 7.36 | 0 | 1 | 0 | 0 |
| 91_0 | 5.1 | 0 | 1 | 0 | 0 |
| 91_1 | 5.1 | 0 | 1 | 0 | 0 |
| 92_0 | 5.77 | 0 | 1 | 0 | 0 |
| 92_1 | 5.77 | 0 | 1 | 0 | 0 |
| 93_0 | 6.52 | 0 | 1 | 0 | 0 |
| 93_1 | 6.52 | 0 | 1 | 0 | 0 |
| 94_0 | 6.93 | 0 | 1 | 0 | 0 |
| 94_1 | 6.93 | 0 | 1 | 0 | 0 |
| 95_0 | 7.36 | 0 | 1 | 0 | 0 |
| 102_0 | 134.58 | 0 | 1 | 0 | 0 |
| 102_1 | 134.58 | 0 | 1 | 0 | 0 |
| 107_0 | 177.57 | 0 | 1 | 0 | 0 |
| 113_0 | 234.29 | 0 | 1 | 0 | 0 |
| 113_1 | 234.29 | 0 | 1 | 0 | 0 |
| 119_0 | 269.13 | 0 | 1 | 0 | 0 |
| 119_1 | 269.13 | 0 | 1 | 0 | 0 |
| 125_0 | 309.14 | 0 | 1 | 0 | 0 |
| 125_1 | 309.14 | 0 | 1 | 0 | 0 |
| 233_0 | 10.5849 | 0 | 1 | 0 | 0 |
| 234_0 | 21.1698 | 0 | 1 | 0 | 0 |
| 235_0 | 42.3396 | 0 | 1 | 0 | 0 |
| 236_0 | 84.6793 | 0 | 1 | 0 | 0 |
| 237_0 | 169.359 | 0 | 1 | 0 | 0 |
| 238_0 | 338.717 | 0 | 1 | 0 | 0 |
| 239_0 | 677.434 | 0 | 1 | 0 | 0 |

| id | cext | cint | nrf2 | P | H |
|---|---|---|---|---|---|
| 240_0 | 1354.87 | 0 | 1 | 0 | 0 |
| 241_0 | 281.571 | 0 | 1 | 0 | 0 |
| 242_0 | 337.886 | 0 | 1 | 0 | 0 |
| 243_0 | 405.463 | 0 | 1 | 0 | 0 |
| 244_0 | 486.556 | 0 | 1 | 0 | 0 |
| 245_0 | 583.867 | 0 | 1 | 0 | 0 |
| 247_0 | 700.64 | 0 | 1 | 0 | 0 |
| 249_0 | 840.768 | 0 | 1 | 0 | 0 |
| 251_0 | 1008.92 | 0 | 1 | 0 | 0 |
| 253_0 | 1210.71 | 0 | 1 | 0 | 0 |
| 254_0 | 1452.85 | 0 | 1 | 0 | 0 |
| 255_0 | 137.422 | 0 | 1 | 0 | 0 |
| 256_0 | 164.906 | 0 | 1 | 0 | 0 |
| 257_0 | 197.888 | 0 | 1 | 0 | 0 |
| 258_0 | 237.465 | 0 | 1 | 0 | 0 |
| 259_0 | 284.958 | 0 | 1 | 0 | 0 |
| 260_0 | 341.95 | 0 | 1 | 0 | 0 |
| 261_0 | 410.34 | 0 | 1 | 0 | 0 |
| 262_0 | 492.408 | 0 | 1 | 0 | 0 |
| 27_0 | 238.256 | 0 | 1 | 0 | 0 |
| 28_0 | 238.256 | 0 | 1 | 0 | 0 |
| 33_0 | 238.256 | 0 | 1 | 0 | 0 |
| 40_0 | 306.652 | 0 | 1 | 0 | 0 |
| 48_0 | 134.792 | 0 | 1 | 0 | 0 |
| 48_1 | 134.792 | 0 | 1 | 0 | 0 |
| 48_2 | 134.792 | 0 | 1 | 0 | 0 |
| 48_3 | 134.792 | 0 | 1 | 0 | 0 |
| 48_4 | 134.792 | 0 | 1 | 0 | 0 |
| 48_5 | 134.792 | 0 | 1 | 0 | 0 |
| 48_6 | 134.792 | 0 | 1 | 0 | 0 |
| 49_0 | 309.229 | 0 | 1 | 0 | 0 |
| 49_1 | 309.229 | 0 | 1 | 0 | 0 |
| 49_2 | 309.229 | 0 | 1 | 0 | 0 |
| 49_3 | 309.229 | 0 | 1 | 0 | 0 |
| 49_4 | 309.229 | 0 | 1 | 0 | 0 |
| 49_5 | 309.229 | 0 | 1 | 0 | 0 |
| 49_6 | 309.229 | 0 | 1 | 0 | 0 |
| 58_0 | 134.792 | 0 | 1 | 0 | 0 |
| 58_1 | 134.792 | 0 | 1 | 0 | 0 |
| 58_2 | 134.792 | 0 | 1 | 0 | 0 |
| 59_0 | 309.229 | 0 | 1 | 0 | 0 |
| 59_1 | 309.229 | 0 | 1 | 0 | 0 |
| 59_2 | 309.229 | 0 | 1 | 0 | 0 |
| 5_0 | 348.834 | 0 | 1 | 0 | 0 |
| 60_0 | 134.792 | 0 | 1 | 0 | 0 |
| 60_1 | 134.792 | 0 | 1 | 0 | 0 |
| 61_0 | 309.229 | 0 | 1 | 0 | 0 |
| 61_1 | 309.229 | 0 | 1 | 0 | 0 |
| 63_0 | 134.792 | 0 | 1 | 0 | 0 |
| 63_1 | 134.792 | 0 | 1 | 0 | 0 |

| id | cext | cint | nrf2 | P | H |
|---|---|---|---|---|---|
| 63_2 | 134.792 | 0 | 1 | 0 | 0 |
| 63_3 | 134.792 | 0 | 1 | 0 | 0 |
| 64_0 | 309.229 | 0 | 1 | 0 | 0 |
| 64_1 | 309.229 | 0 | 1 | 0 | 0 |
| 64_2 | 309.229 | 0 | 1 | 0 | 0 |
| 64_3 | 309.229 | 0 | 1 | 0 | 0 |
| 65_0 | 134.792 | 0 | 1 | 0 | 0 |
| 65_1 | 134.792 | 0 | 1 | 0 | 0 |
| 65_2 | 134.792 | 0 | 1 | 0 | 0 |
| 65_3 | 134.792 | 0 | 1 | 0 | 0 |
| 65_4 | 134.792 | 0 | 1 | 0 | 0 |
| 65_5 | 134.792 | 0 | 1 | 0 | 0 |
| 66_0 | 309.229 | 0 | 1 | 0 | 0 |
| 66_1 | 309.229 | 0 | 1 | 0 | 0 |
| 66_2 | 309.229 | 0 | 1 | 0 | 0 |
| 66_3 | 309.229 | 0 | 1 | 0 | 0 |
| 66_4 | 309.229 | 0 | 1 | 0 | 0 |
| 66_5 | 309.229 | 0 | 1 | 0 | 0 |
| 6_0 | 348.834 | 0 | 1 | 0 | 0 |
| 76_0 | 134.792 | 0 | 1 | 0 | 0 |
| 76_1 | 134.792 | 0 | 1 | 0 | 0 |
| 76_10 | 134.792 | 0 | 1 | 0 | 0 |
| 76_2 | 134.792 | 0 | 1 | 0 | 0 |
| 76_3 | 134.792 | 0 | 1 | 0 | 0 |
| 76_4 | 134.792 | 0 | 1 | 0 | 0 |
| 76_5 | 134.792 | 0 | 1 | 0 | 0 |
| 76_6 | 134.792 | 0 | 1 | 0 | 0 |
| 76_7 | 134.792 | 0 | 1 | 0 | 0 |
| 76_8 | 134.792 | 0 | 1 | 0 | 0 |
| 76_9 | 134.792 | 0 | 1 | 0 | 0 |
| 77_0 | 309.229 | 0 | 1 | 0 | 0 |
| 77_1 | 309.229 | 0 | 1 | 0 | 0 |
| 77_2 | 309.229 | 0 | 1 | 0 | 0 |
| 77_3 | 309.229 | 0 | 1 | 0 | 0 |
| 77_4 | 309.229 | 0 | 1 | 0 | 0 |
| 77_5 | 309.229 | 0 | 1 | 0 | 0 |
| 77_6 | 309.229 | 0 | 1 | 0 | 0 |
| 77_7 | 309.229 | 0 | 1 | 0 | 0 |
| 77_8 | 309.229 | 0 | 1 | 0 | 0 |

###### S5.2.3 Free parameters

- error_scale ∼ lognorm(scale=1.0,s=0.1,dims=())
- error_scale_sigma ∼ halfnorm(scale=0.1,dims=())
- sigma_experiment ∼ lognorm(scale=error_scale,s=error_scale_sigma,dims=('experiment_id',))
- k_i_substance ∼ lognorm(scale=[1.0,1.0,1.0],s=2,dims=('substance',))
- z_ci_substance ∼ lognorm(scale=[0.5,0.5,0.5],s=2,dims=('substance',))
- k_m_substance ∼ lognorm(scale=[0.05,0.05,0.05],s=2,dims=('substance',))
- r_rt ∼ lognorm(scale=1.0,s=2,dims=())
- r_rd ∼ lognorm(scale=0.5,s=2,dims=())

- v_rt ~ lognorm(scale=1.0,s=2,dims=())
- k_p ~ lognorm(scale=0.02,s=2,dims=())
- h_b ~ lognorm(scale=1e-08,s=2,dims=())
- z ~ lognorm(scale=1.0,s=2,dims=())
- kk ~ lognorm(scale=0.02,s=2,dims=())
- sigma_nrf2 ~ halfnorm(scale=5.0,dims=())
- sigma_cint ~ halfnorm(scale=5.0,dims=())

###### S5.2.4 Fixed parameters

- cext_y0 = cext_y0 * sigma_experiment[experiment_id_index], dims=('id',)
- k_i = k_i_substance[substance_index], dims=('id',)
- z_ci = z_ci_substance[substance_index], dims=('id',)
- k_m = k_m_substance[substance_index], dims=('id',)
- ci_max_substance = [1757.0, 168.1, 6364.8], dims=('substance',)
- ci_max = ci_max_substance[substance_index], dims=('id',)
- r_0 = 1.0, dims=()

##### S5.3 Report: Table parameter estimates ✓

Excluding parameters: ['sigma_experiment'] for meaningful visualization

| | ('index', ' ') | ('diuron', 'mean ± std') | ('diclofenac', 'mean ± std') | ('naproxen', 'mean ± std') |
|---|---|---|---|---|
| 0 | ci_max_substance | 1757.0 ± 0.0 | 168.1 ± 0.0 | 6364.8 ± 0.0 |
| 1 | error_scale | 1.032 ± 0.083 | 1.032 ± 0.083 | 1.032 ± 0.083 |
| 2 | error_scale_sigma | 0.581 ± 0.04 | 0.581 ± 0.04 | 0.581 ± 0.04 |
| 3 | h_b | 0.0 ± 0.0 | 0.0 ± 0.0 | 0.0 ± 0.0 |
| 4 | k_i_substance | 4.899 ± 0.659 | 0.379 ± 0.033 | 0.362 ± 0.034 |
| 5 | k_m_substance | 3.776 ± 0.627 | 0.187 ± 0.022 | 0.084 ± 0.01 |
| 6 | k_p | 0.008 ± 0.001 | 0.008 ± 0.001 | 0.008 ± 0.001 |
| 7 | kk | 0.079 ± 0.008 | 0.079 ± 0.008 | 0.079 ± 0.008 |
| 8 | r_rd | 0.232 ± 0.017 | 0.232 ± 0.017 | 0.232 ± 0.017 |
| 9 | r_rt | 2.301 ± 0.252 | 2.301 ± 0.252 | 2.301 ± 0.252 |
| 10 | sigma_cint | 0.673 ± 0.024 | 0.673 ± 0.024 | 0.673 ± 0.024 |
| 11 | sigma_nrf2 | 0.278 ± 0.01 | 0.278 ± 0.01 | 0.278 ± 0.01 |
| 12 | v_rt | 3.731 ± 0.115 | 3.731 ± 0.115 | 3.731 ± 0.115 |
| 13 | z | 1.535 ± 0.033 | 1.535 ± 0.033 | 1.535 ± 0.033 |
| 14 | z_ci_substance | 1.16 ± 0.057 | 1.048 ± 0.036 | 1.172 ± 0.042 |

Report 'table_parameter_estimates' was successfully generated and saved in '../hierarchical_molecular_tktd/results

##### S5.4 Report: Goodness of fit ✓

| | cint | nrf2 | survival | model |
|---|---|---|---|---|
| NRMSE | 0.11985 | 0.158582 | 0.35749 | nan |
| NRMSE (95%-hdi[lower]) | 0.102319 | 0.15361 | 0.340377 | nan |
| NRMSE (95%-hdi[upper]) | 0.140886 | 0.163834 | 0.373375 | nan |
| Log-Likelihood | -950.513 | -25.4162 | -289.769 | -1265.7 |
| Log-Likelihood (95%-hdi[lower]) | -975.42 | -27.6206 | -307.429 | -1293.27 |
| Log-Likelihood (95%-hdi[upper]) | -925.284 | -23.3595 | -275.331 | -1237.01 |

| | cint | nrf2 | survival | model |
|---|---|---|---|---|
| n (data) | 913 | 169 | 392 | 1474 |
| k (parameters) | nan | nan | nan | 62 |
| BIC | nan | nan | nan | 2983.73 |
| BIC (95%-hdi[lower]) | nan | nan | nan | 2926.36 |
| BIC (95%-hdi[upper]) | nan | nan | nan | 3038.87 |

Report ‘goodness_of_fit’ was successfully generated and saved in ‘../hierarchical_molecular_tktd/results/hierarchica

#### S5.5 Report: Diagnostics ✓

Excluding parameters: [‘sigma_experiment’] for meaningful visualization

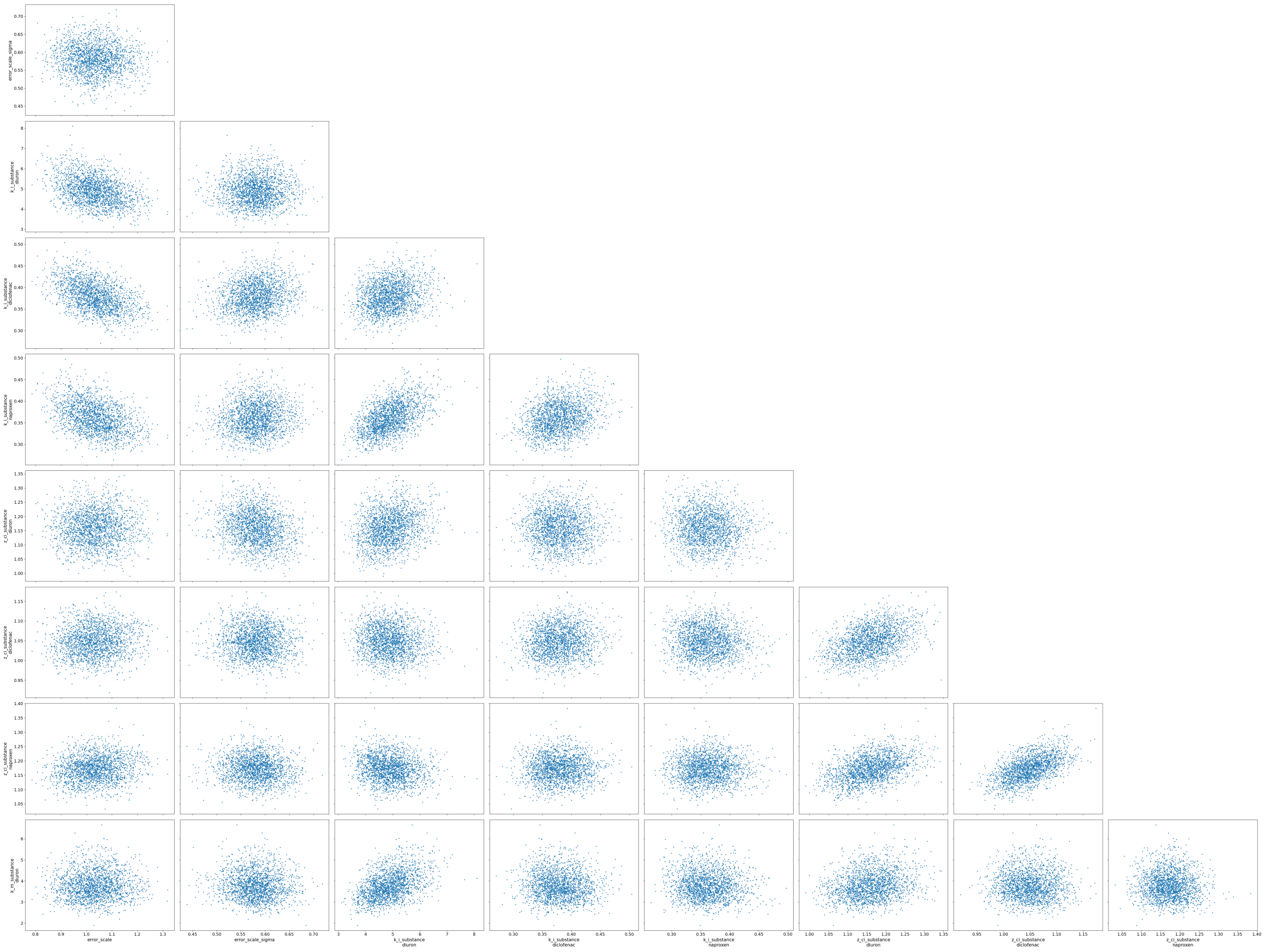

**Figure S29.** Paired parameter estimates

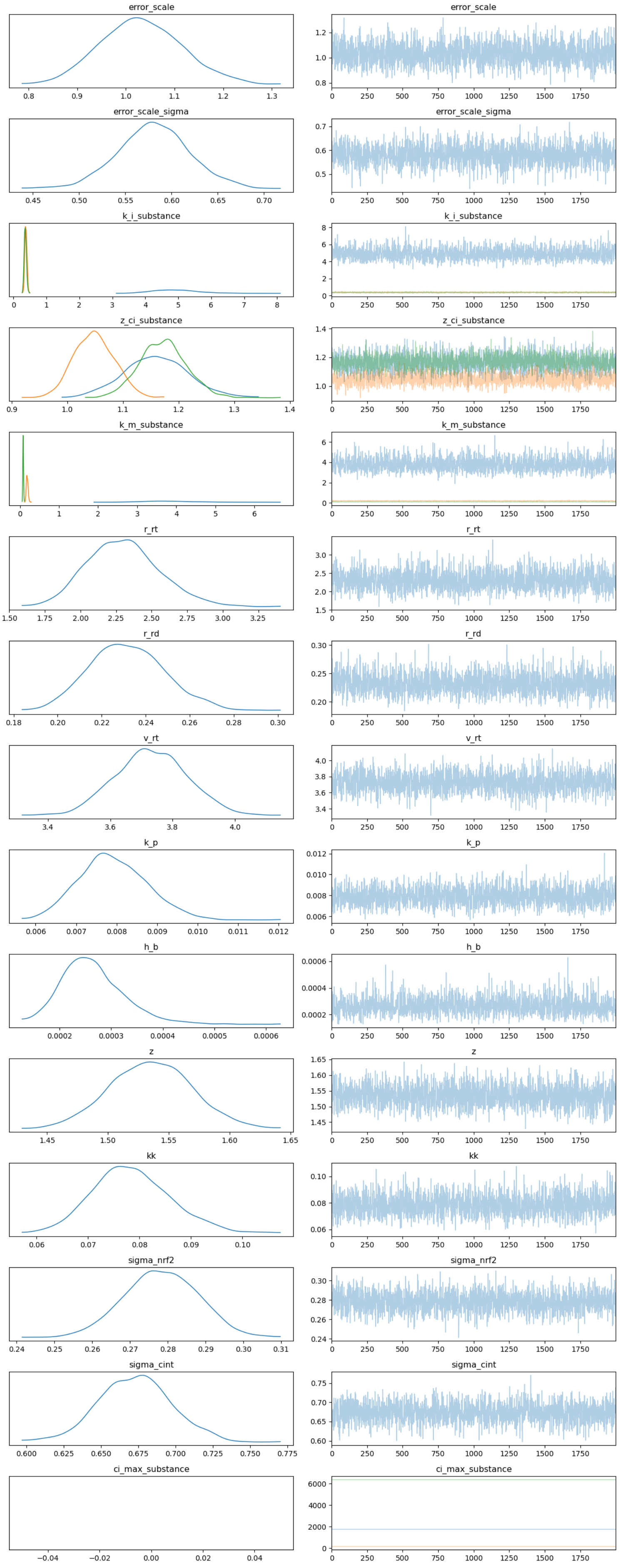


**Figure S30.** Psuedo trace, generated for draws from the optimized SVI distribution

Report ‘diagnostics’ was successfully generated and saved in ‘(’../hierarchical_molecular_tktd/results/hierarchical_[illegible]
‘../hierarchical_molecular_tktd/results/hierarchical_cext_nested_sigma_hyperprior_informed_rna_pulse_5_substance_in[illegible]

#### S5.6 Report: Visualizations ✓

#### S5.7 Report: Visualizations ✓

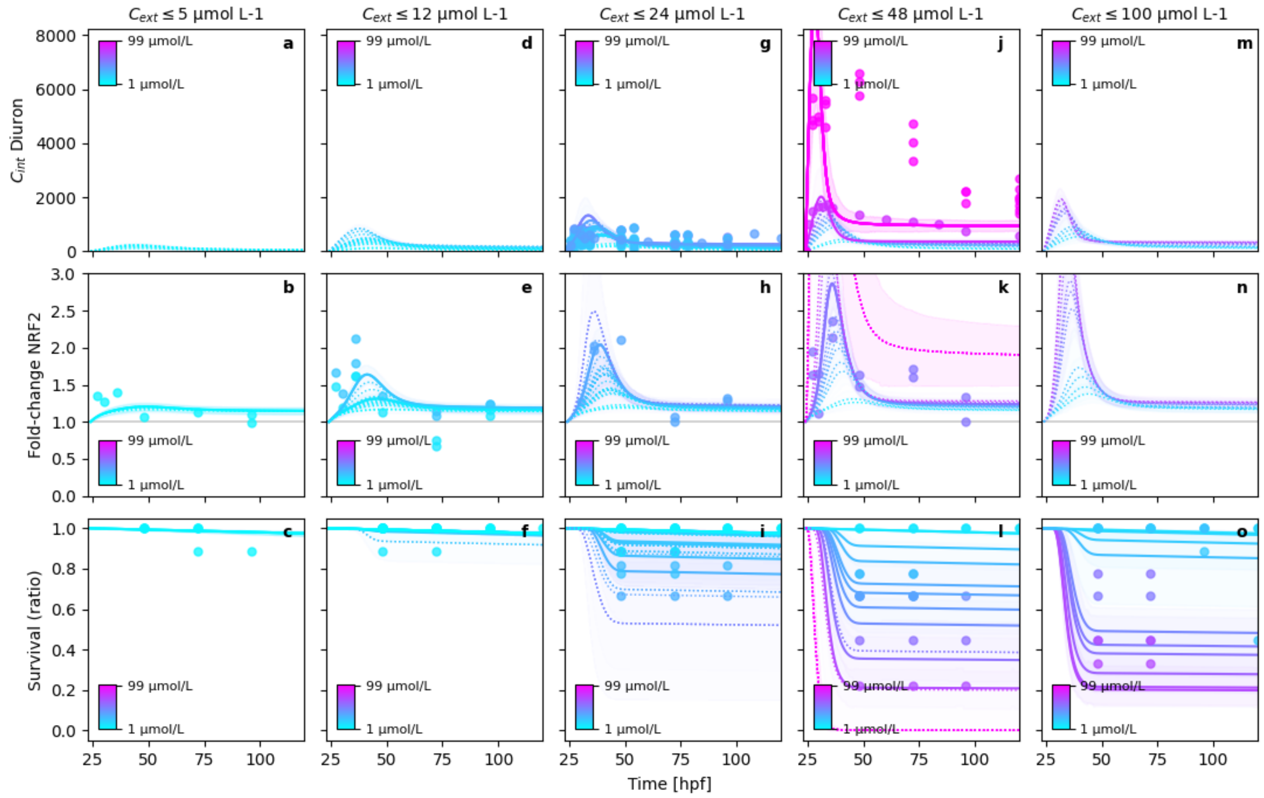


**Figure S31.** Posterior model fits

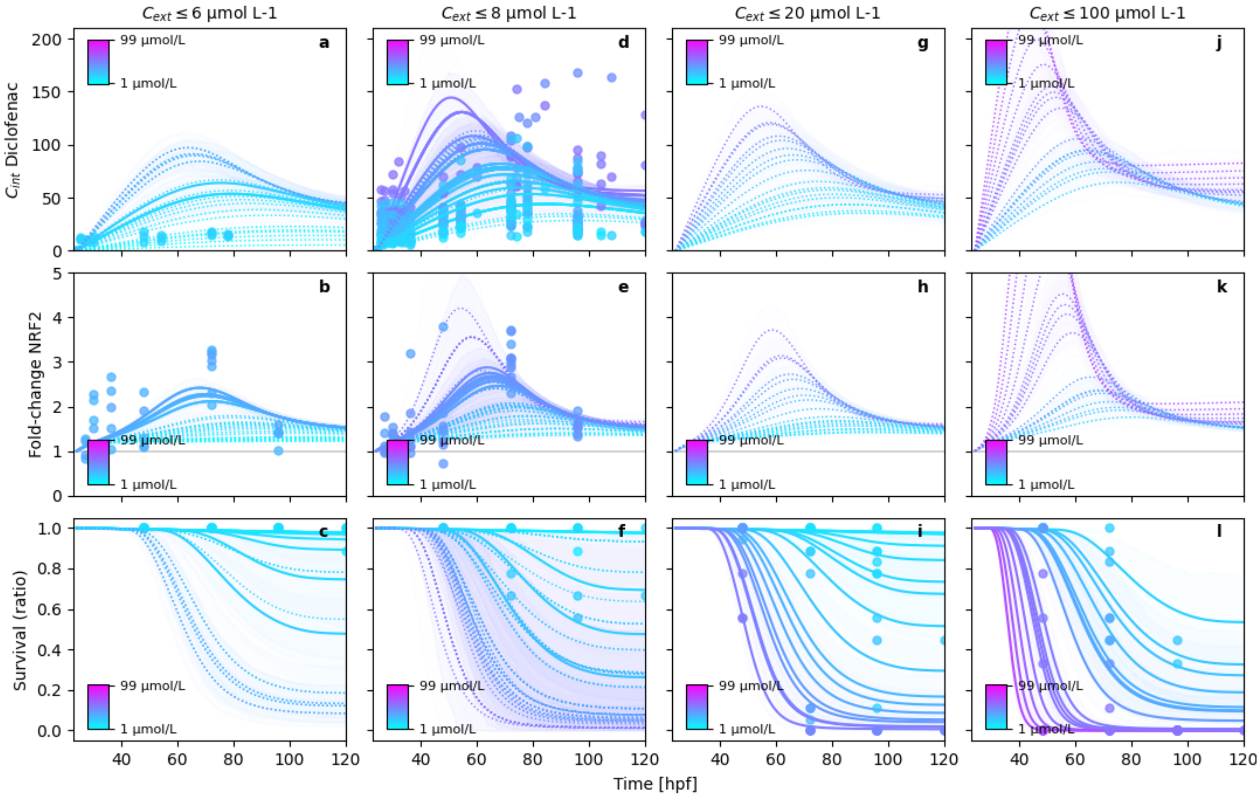


**Figure S32.** Posterior model fits

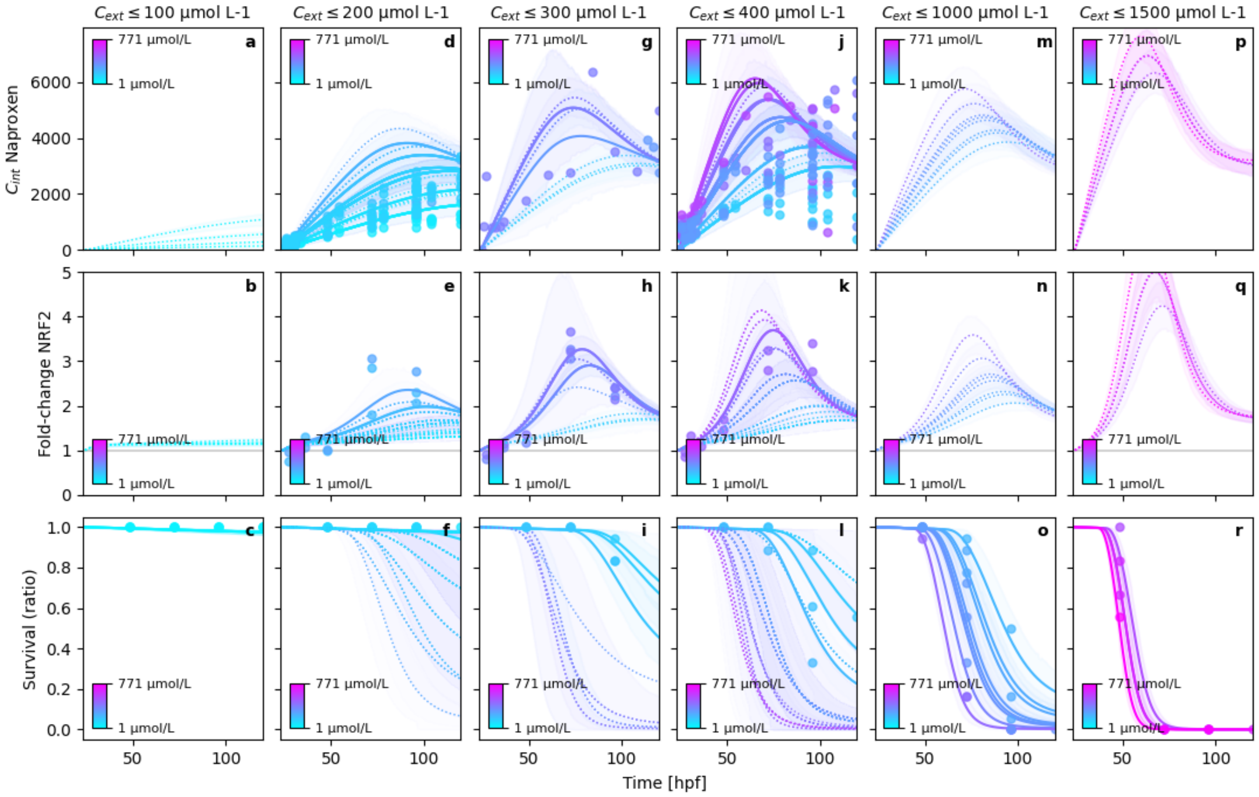


**Figure S33.** Posterior model fits

Report 'visualizations' was successfully generated and saved in '['../hierarchical_molecular_tktd/results/hierarchical_c
'../hierarchical_molecular_tktd/results/hierarchical_cext_nested_sigma_hyperprior_informed_rna_pulse_5_substance_in

’../hierarchical_molecular_tktd/results/hierarchical_cext_nested_sigma_hyperprior_informed_rna_pulse_5_substance_in

##### S5.7.1 $y_0$ estimation of external concentrations

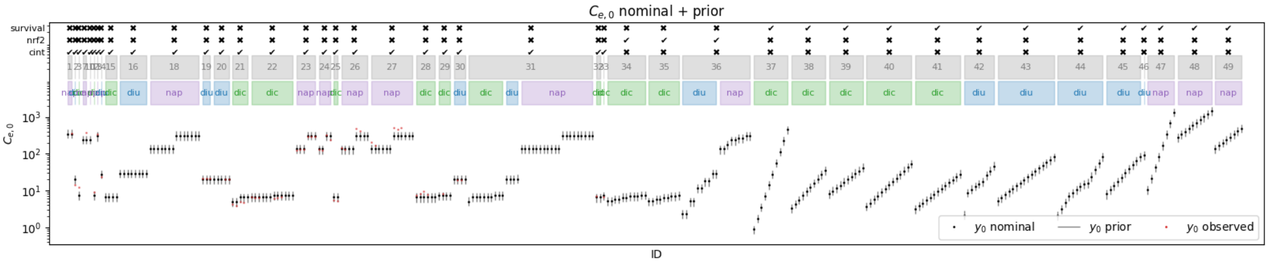


**Figure S34.** Prior $C_{ext,0}$ estimates and nominal concentrations

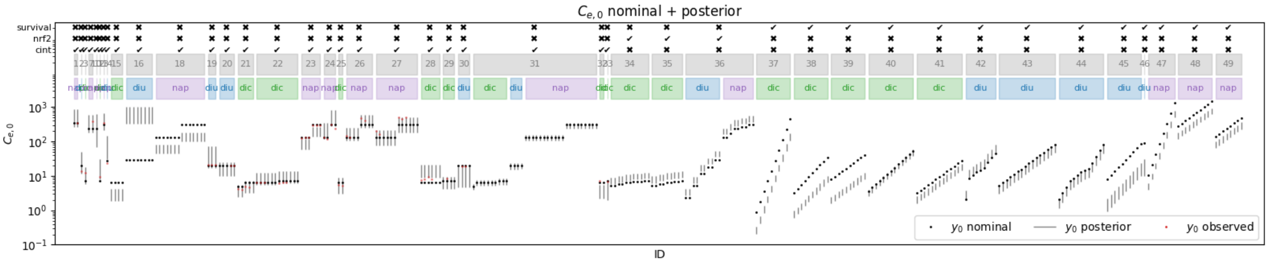


**Figure S35.** Posterior $C_{ext,0}$ estimates and nominal concentrations

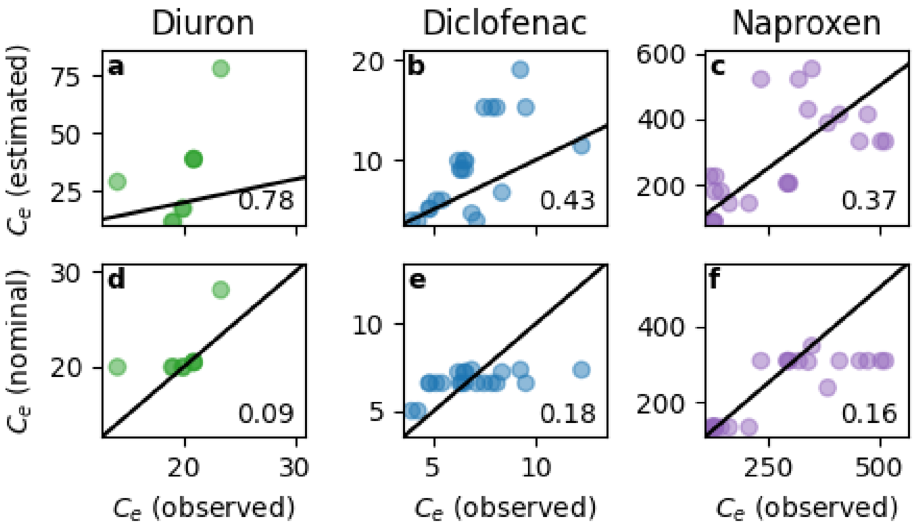


**Figure S36.** $C_{ext,0}$ comparison between nominal, measured and estimated concentrations

Report ‘visualizations’ was successfully generated and saved in ‘(’../hierarchical_molecular_tktd/results/hierarchical_c
‘../hierarchical_molecular_tktd/results/hierarchical_cext_nested_sigma_hyperprior_informed_rna_pulse_5_substance_in
‘../hierarchical_molecular_tktd/results/hierarchical_cext_nested_sigma_hyperprior_informed_rna_pulse_5_substance_in

#### S5.8 Report: Model inadequacy metrics ✓

##### S5.8.1 Residuals

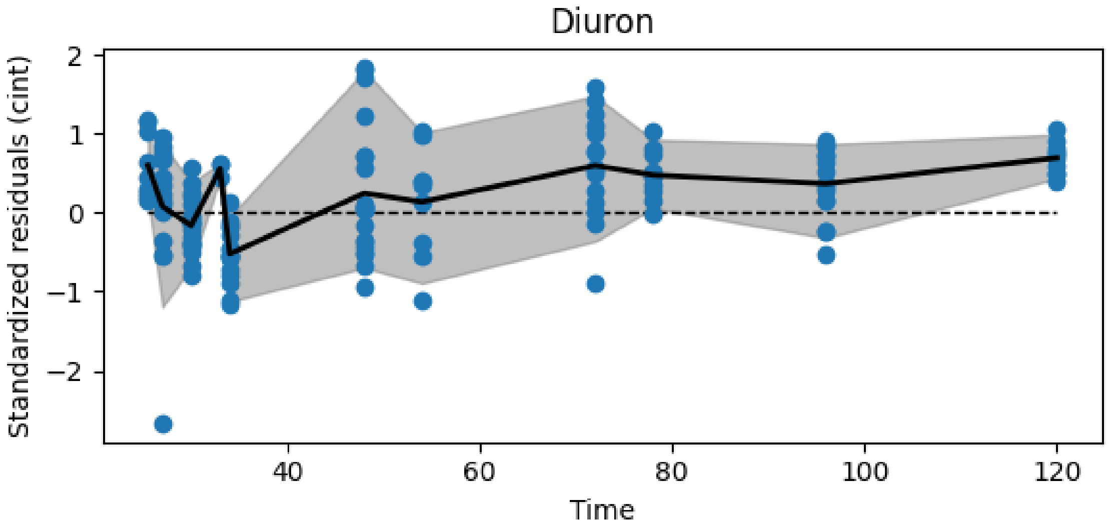


**Figure S37.** Resiudal cint dynamics of diuron

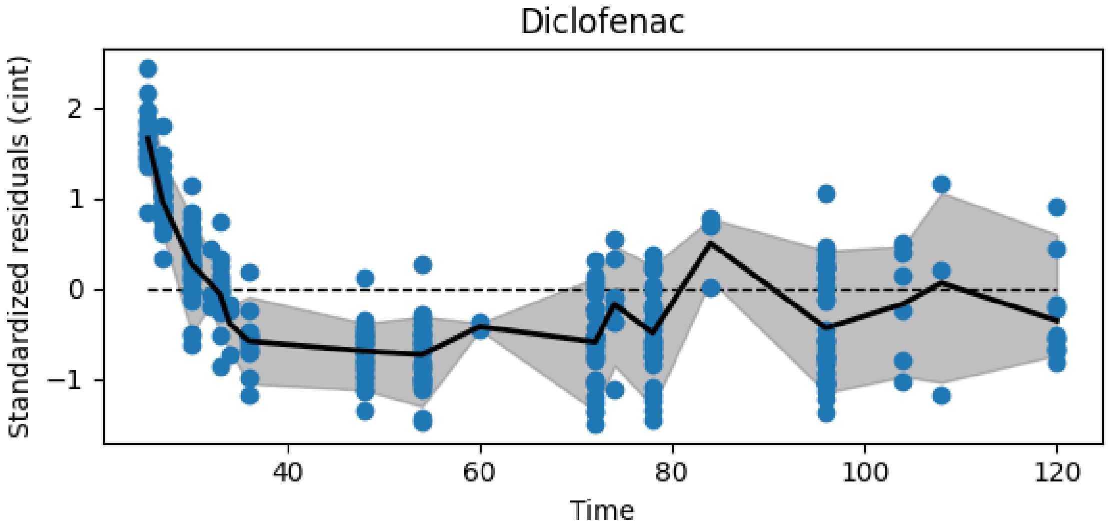


**Figure S38.** Resiudal cint dynamics of diclofenac

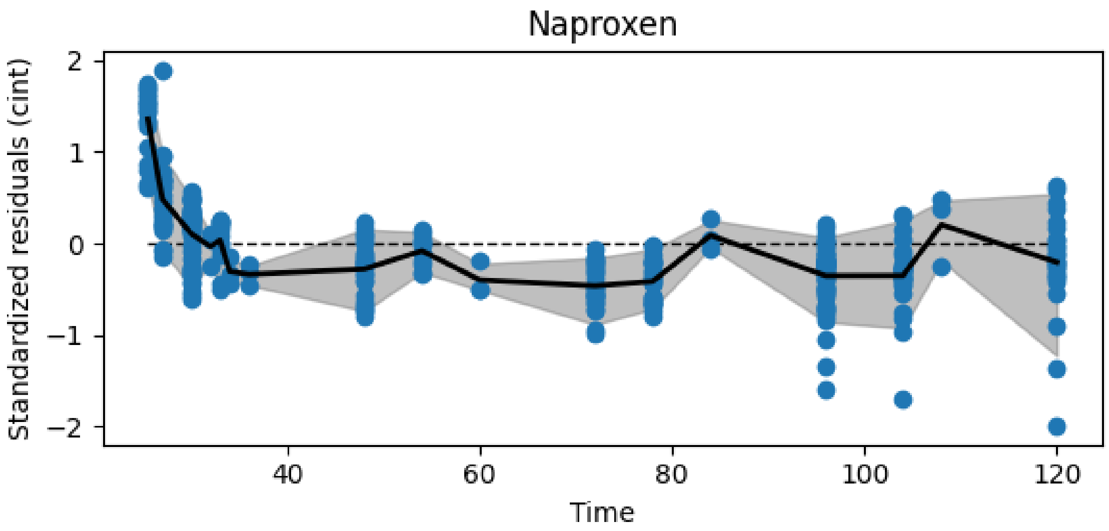


**Figure S39.** Resiudal cint dynamics of naproxen

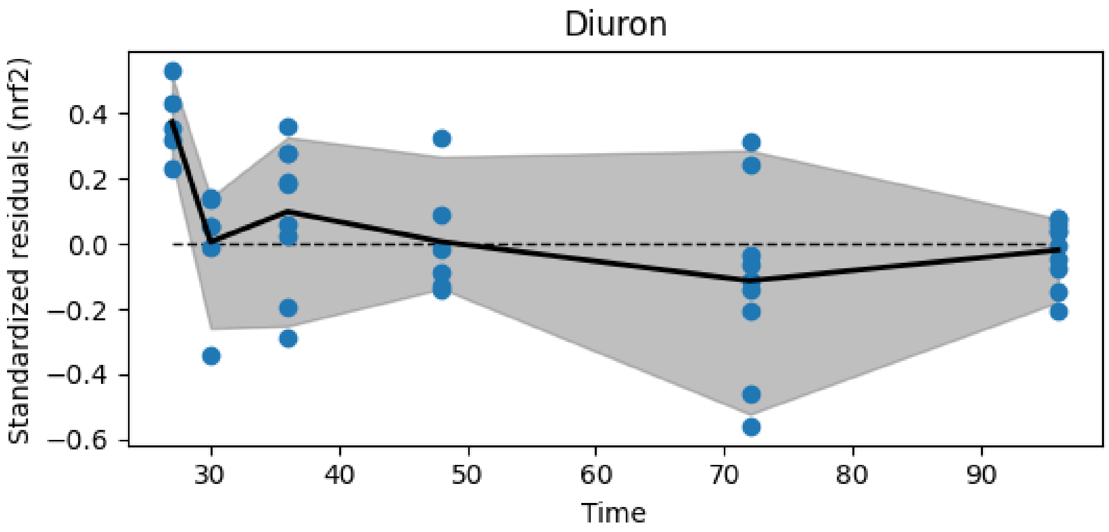


**Figure S40.** Resiudal nrf2 dynamics of diuron

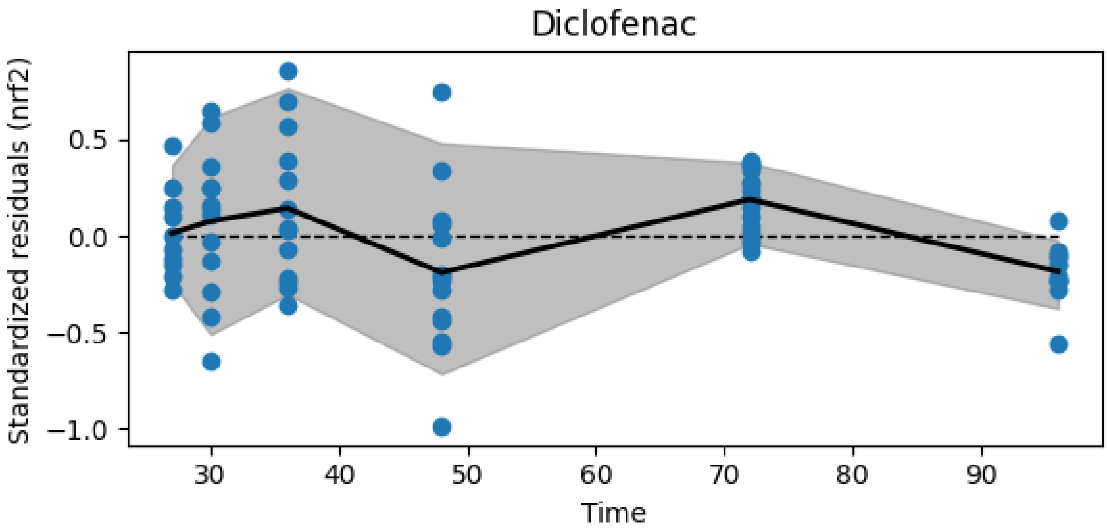


**Figure S41.** Resiudal nrf2 dynamics of diclofenac

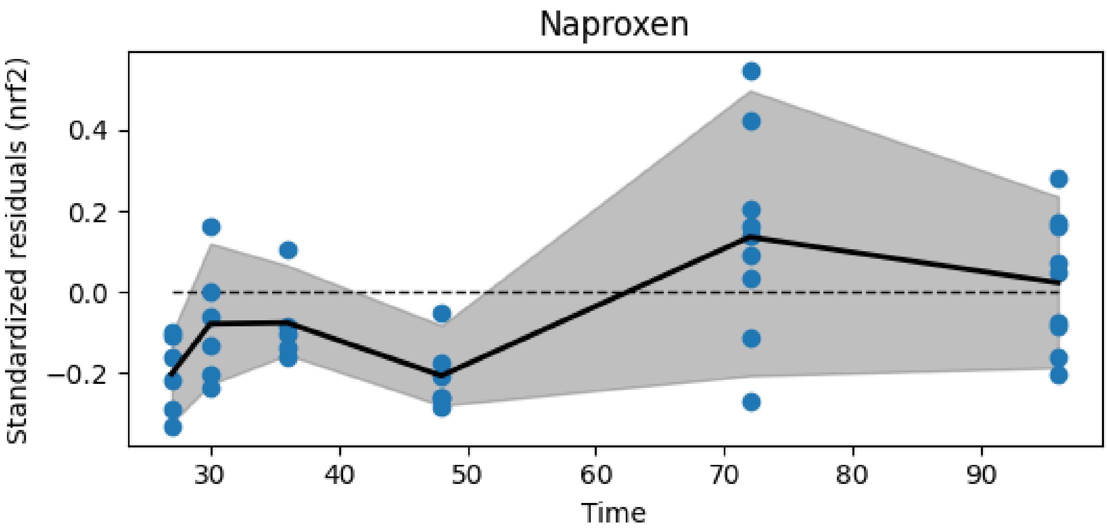


**Figure S42.** Resiudal nrf2 dynamics of naproxen

##### S5.8.2 Model inadequacy

The different metrics are measures for the model inadequacy. The comparison to `metric_value_if_normal_dist` is a simulation of normally distributed residuals that have the same data structure in terms of dimensionality (id x time) and missing values. If the `metric_value` falls within that interval, the model can be assumed as not inadequate.

- **autocorrelation**: Measures the correlation of the residuals with themselves with a lag of 1. High absolute autocorrelation means, the variable is not normally distributed. Ideal would be values close to 0.
- **deviation log-prob**: Uses a t-test to estimate the probability of the replicates at a time t being different from zero. The result is the summed log-probability. Low (negative) log probs indicate high probability for deviation.
- **significant deviations**: Uses a t-test to estimate the probability of the replicates at a time t being different from zero. The result is the number of significant deviations (for an alpha level of 0.05). High number of deviations indicate an inadequate model

- **local/global variance**: This metric calculates the local variance as a rolling variance of always 3 direct neighboring residuals. The local variances are then averaged and divided by the global averages of all residuals. The basis for the calculation is the residuals averaged by id. Values close to 1 indicate an adequate model
- **replicate/global variance**: This metric calculates the replicate variance at time t, averages it and divides the number by the global variance. The local variances are then averaged and divided by the global averages of all residuals. The basis for the calculation is the residuals averaged by id. Values close to 1 indicate an adequate model

| | metric | data_variable | index | metric_value | metric_value_if_normal_dist |
|---|---|---|---|---|---|
| 0 | autocorrelation | cint | diclofenac | 0.678054 | -0.07[-0.4,0.25] |
| 1 | autocorrelation | cint | diuron | -0.186462 | -0.1[-0.63,0.41] |
| 2 | autocorrelation | cint | naproxen | 0.620828 | -0.08[-0.42,0.21] |
| 3 | autocorrelation | nrf2 | diclofenac | -0.815108 | -0.2[-0.76,0.55] |
| 4 | autocorrelation | nrf2 | diuron | 0.116714 | -0.2[-0.83,0.56] |
| 5 | autocorrelation | nrf2 | naproxen | -0.0656427 | -0.14[-0.85,0.6] |
| 6 | deviation log-prob | cint | diclofenac | -216.49 | -17.75[-22.96,-12.42] |
| 7 | deviation log-prob | cint | diuron | -56.7554 | -10.56[-16.58,-5.91] |
| 8 | deviation log-prob | cint | naproxen | -146.351 | -16.89[-24.17,-9.83] |
| 9 | deviation log-prob | nrf2 | diclofenac | -23.0962 | -6.29[-10.42,-3.0] |
| 10 | deviation log-prob | nrf2 | diuron | -9.86738 | -5.76[-11.27,-2.64] |
| 11 | deviation log-prob | nrf2 | naproxen | -17.2778 | -5.93[-10.89,-2.35] |
| 12 | local/global variance | cint | diclofenac | 0.222328 | 0.73[0.55,0.92] |
| 13 | local/global variance | cint | diuron | 0.713998 | 0.72[0.41,0.98] |
| 14 | local/global variance | cint | naproxen | 0.251405 | 0.74[0.54,0.9] |
| 15 | local/global variance | nrf2 | diclofenac | 0.94941 | 0.82[0.44,1.19] |
| 16 | local/global variance | nrf2 | diuron | 0.385209 | 0.77[0.44,1.16] |
| 17 | local/global variance | nrf2 | naproxen | 0.815643 | 0.8[0.44,1.21] |
| 18 | replicate/global variance | cint | diclofenac | 0.288974 | 0.87[0.75,1.01] |
| 19 | replicate/global variance | cint | diuron | 0.604549 | 0.91[0.83,0.99] |
| 20 | replicate/global variance | cint | naproxen | 0.26346 | 0.87[0.72,1.01] |
| 21 | replicate/global variance | nrf2 | diclofenac | 0.810006 | 0.94[0.87,1.0] |
| 22 | replicate/global variance | nrf2 | diuron | 0.601163 | 0.88[0.72,0.99] |
| 23 | replicate/global variance | nrf2 | naproxen | 0.511345 | 0.88[0.71,1.01] |
| 24 | significant deviations | cint | diclofenac | 11 | 0.9[0.0,2.05] |
| 25 | significant deviations | cint | diuron | 7 | 0.57[0.0,2.0] |
| 26 | significant deviations | cint | naproxen | 9 | 0.87[0.0,2.0] |
| 27 | significant deviations | nrf2 | diclofenac | 2 | 0.3[0.0,1.0] |
| 28 | significant deviations | nrf2 | diuron | 1 | 0.21[0.0,1.0] |
| 29 | significant deviations | nrf2 | naproxen | 2 | 0.26[0.0,1.0] |

Report 'model_inadequacy_metrics' was successfully generated and saved in '../hierarchical_molecular_tktd/results
